# Targeted Degradation of METTL3 Against Acute Myeloid Leukemia and Gastric Cancer

**DOI:** 10.1101/2024.02.02.578521

**Authors:** Kyubin Hwang, Juhyeon Bae, Yoo-Lim Jhe, Jungmin Kim, Jae-Ho Cheong, Taebo Sim

## Abstract

Accumulating evidence reveals the oncogenic role of methyltransferase-like 3 (METTL3) in a variety of cancer types, either dependent or independent of its m^6^A methyl transferase activity. We have designed proteolysis-targeting chimeras (PROTACs) targeting METTL3 and identified **KH12** as a potent METTL3 degrader. Treatment of **KH12** on MOLM-13 cells causes more than 80% degradation of METTL3 with a half-maximal degradation concentration (DC_50_) of 220 nM in a dose-, time- and ubiquitin-dependent fashion. In addition, **KH12** reverses differentiation and possesses anti-proliferative effects surpassing the reported inhibitors in MOLM-13 cells. Furthermore, **KH12** significantly suppresses the growth of various gastric cancer (GC) cells, where the m^6^A-independent activity of METTL3 plays a crucial role in tumorigenesis. The anti-GC effect of **KH12** was further confirmed in patient-derived organoids (PDOs). This study highlights the therapeutic potential of targeted degradation of epitranscriptomic writer METTL3 as an anti-cancer strategy.

## INTRODUCTION

*N*^6^-methyladenosine (m^6^A) is the most prevalent chemical modification found on the internal mRNA of mammals. This epitranscriptomic modification plays crucial and diverse roles in regulating gene expression, generating various physiological, developmental, or pathological outcomes^1-3^. Three types of protein machinery called “writers”, “erasers”, and “readers” are respectively involved in installing, removing, and interpreting the m^6^A RNA modifications^4^. Methyltransferase-like 3 (METTL3) was originally identified as a m^6^A writer^5^. By forming a stable complex with methyltransferase-like 14 (METTL14), METTL3 installs m^6^A modification on a broad spectrum of RNA targets^6^.

METTL3 is frequently overexpressed in a variety of cancers and has been linked to the promotion of carcinogenesis^7,8^. In acute myeloid leukemia (AML), METTL3 enhances the translation of genes including BCL2 and c-MYC by inducing m^6^A modification to their transcript^9^. Consequently, METTL3 stalls the differentiation of hematopoietic stem/progenitor cells thereby establishing a leukemia state^9,10^. Despite its significant role as a methyltransferase in cancer, accumulating evidences suggest that the methyltransferase-independent function of cytoplasmic METTL3 promotes cancer progression^11-13^. In lung cancer, METTL3 promotes the translation of oncogenic mRNAs by interacting with translation initiation factor eIF3h and subsequent forming of mRNA loop^11,12^. On the other hand, the interaction between METTL3 and translation initiation factor PABPC1 was reported to elevate the translation of epigenetic factors and promote tumor progression in gastric cancer (GC)^13^. In both cases, biological knockdown of METTL3 significantly reduces the translation efficiency of mRNAs, whereas, catalytically inactive METTL3 remains in its interaction with the translation initiation factors^12,13^.

Currently, several small molecule METTL3 inhibitors targeting its m^6^A methyltransferase activity have been reported^8,14,15^. Though the inhibitors effectively reduced m^6^A levels and exerted subsequent anti-AML effects, catalytic inhibition still spares the methyltransferase-independent functions of METTL3. By employing the proteolysis targeted chimera (PROTAC) strategy, Caflisch group has reported METTL3 PROTAC degraders^16^. Despite their extensive medicinal chemistry campaign, their PROTACs achieved only about 50% degradation of METTL3. On the other hand, Du et al. have most recently reported potent METTL3 PROTACs^17^. However, their applications demonstrated anti-proliferative effects against only AML cells, where maintenance of leukemia state is associated with the conventional catalytic function of METTL3. Thus far, the anti-cancer potentials of targeted degradation of METTL3 have not been fully investigated yet. The study herein reports the discovery of a potent METTL3 PROTAC degrader and its anti-cancer potential against GC as well as against AML.

## RESULTS

### Design, synthesis, and evaluation of METTL3 PROTACs

In order to generate METTL3 PROTACs, we fused a METTL3 inhibitor UZH2 as a warhead with von Hippel-Lindau (VHL) or cereblon (CRBN) E3 ligase ligands. UZH2 is a selective METTL3 inhibitor, which targets the SAM binding domain of METTL3 and effectively impedes its methyltransferase activity with a single-digit nanomolar potency^14^ (Figure 1A). The X-ray co-crystal structure of UZH2 and METTL3 reveals the methylamino group on the pyrimidine of UZH2 is oriented toward the solvent-accessible region, while the dimethylpiperidine group is deeply embedded in the pocket (Figure 1B). Hypothesizing the position of the methylamino group is suitable for the linker attachment, we installed aliphatic or polyethylene glycol (PEG) linkers on the position in order to couple with either VHL or CRBN E3 ligase ligands, thereby producing two series of METTL3 PROTACs: VHL-based (**KH11-16**) and CRBN-based (**KH21-26**) PROTACs, respectively (Figure 1C).

**Figure 1.**
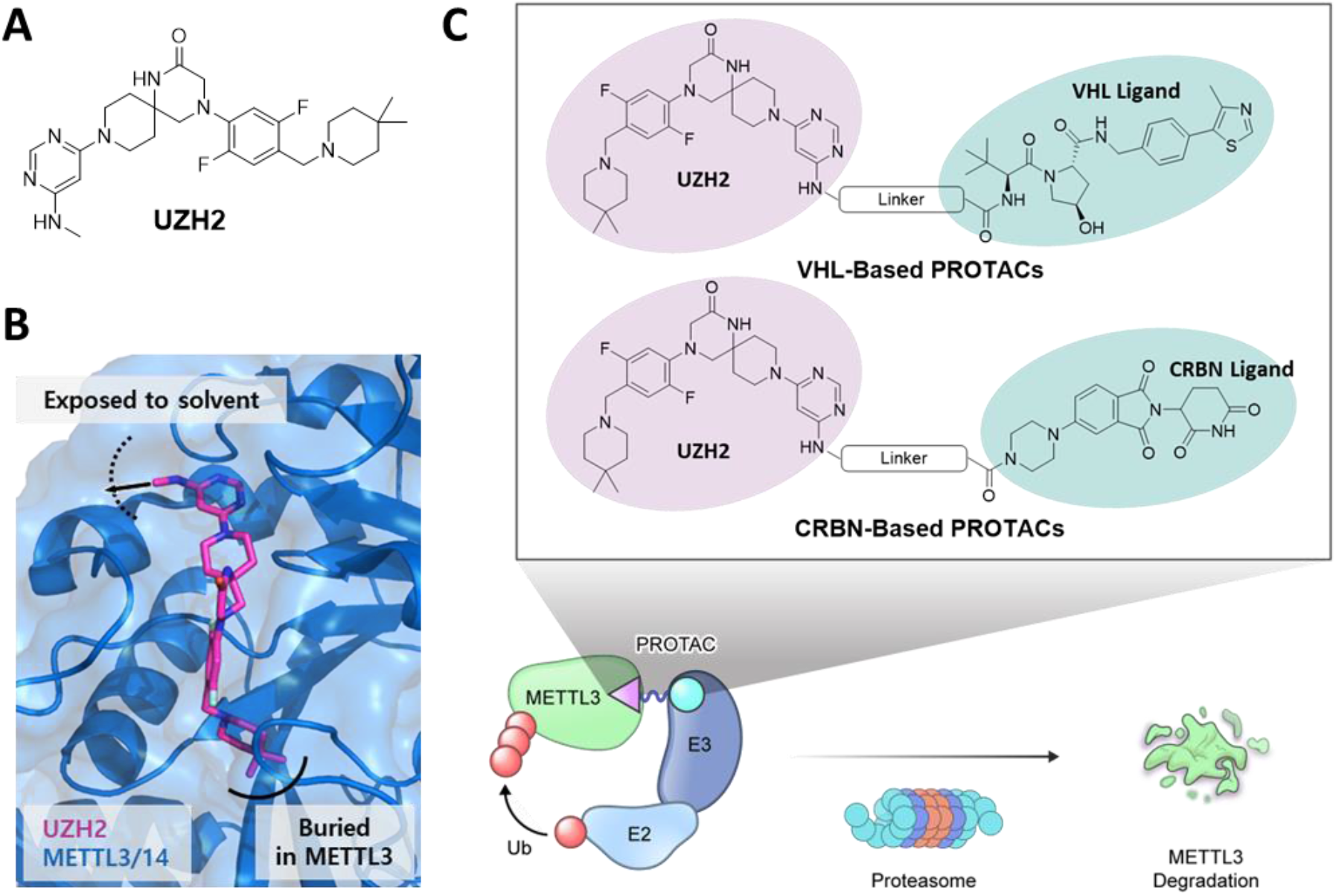
Design of METTL3 PROTACs. A) Structure of METTL3 inhibitor UZH2. B) Crystal structure showing binding mode of UZH2 on METTL3/METTL14 complex (PDB: 7O2F). UZH2 is shown in magenta and an arrow marks the linker vector for the PROTAC design. C) Structure of the designed series of METTL3 PROTACs and schematic of the targeted degradation of METTL3 using PROTAC technology.

We next carried out Western blot analysis to assess METTL3 degradation activities of the PROTACs in MOLM-13 cells. Interestingly, it was found that the composition of linkers adopted has a significant effect on the degradation activity of the PROTACs. Among the aliphatic and polyethylene glycol linkers, only aliphatic linkers could induce METTL3 degradation and replacement of aliphatic linkers by PEG linkers abolishes degradation activities (Table 1, S1 and Figure S1). With the consistent result of a recent report^16^, we concluded PEG linkers are not suitable for the METTL3 PROTAC. This might be due to their hydrophilicity or possible intramolecular hydrogen bonding, which could lead to low cell permeability or an unfavorable orientation for the stable ternary complex formation.

**Table 1.**
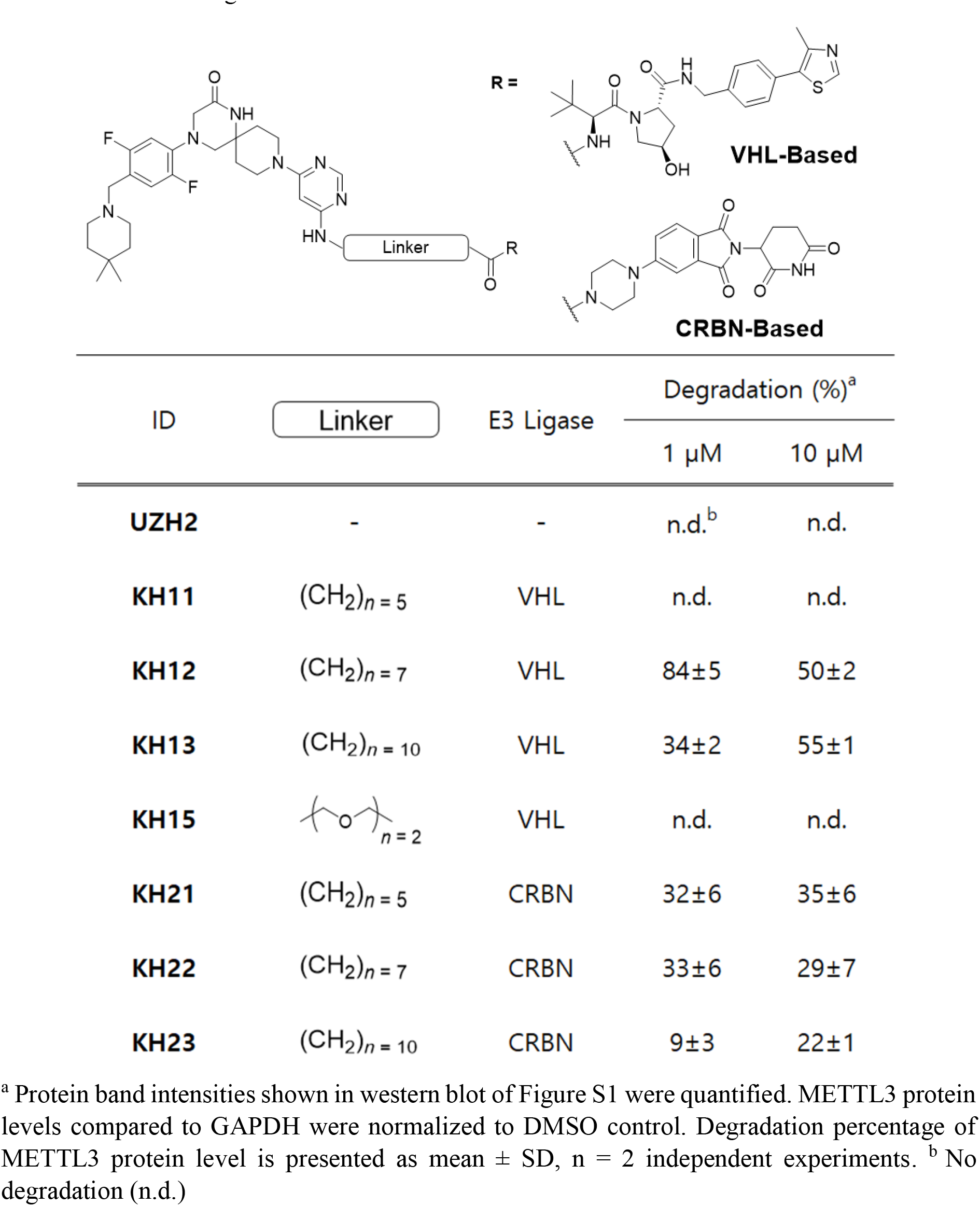
METTL3 degradation activities for PROTACs.

Within the subset of PROTACs with aliphatic linkers, the degradation activities of the VHL-based PROTACs (**KH11-13**) are substantially varied depending on linker length. Among them, **KH12**, incorporating seven-carbon linker is capable of potently inducing degradation of METTL3. We observed 84% METTL3 degradation by the treatment of 1 μM **KH12**, with characteristic hook effect at 10 μM concentration. Alteration of linker length by only 2 to 3 methylene groups (**KH11** and **KH13**) results in completely lost or impaired degradation activity toward cellular METTL3 (Table 1). The analysis of predicted ternary complex of METTL3, VHL, and **KH12** reveals that stretched orientation of the linker composed of seven-methylene group might provide optimal distance to induce METTL3-VHL interaction for productive ternary complex formation (Figure S2). It is worth noting that CRBN-based PROTACs (**KH21-23**) possess diminished degradation activities compared with VHL-based PROTACs (Table 1 and Figure S1).

On the basis of degradation activities of the two series of METTL3 PROTACs, we selected **KH12** as a representative METTL3 degrader. We further examined the degradation activities of **KH12** depending on dose range and treatment time. It was observed that **KH12** induces METTL3 degradation in a dose-dependent manner with a half-maximal degradation concentration (DC_50_) of 220 nM (Figure 2A and B). Upon treatment of **KH12 (**1 μM), the kinetics of METTL3 degradation were monitored and it was found that METTL3 is degraded by over 50% and 90% at 4 and 24 hours, respectively (Figure 2C and D). Taken together, **KH12** is capable of potently and rapidly degrading METTL3 in a dose- and time-dependent manner.

**Figure 2.**
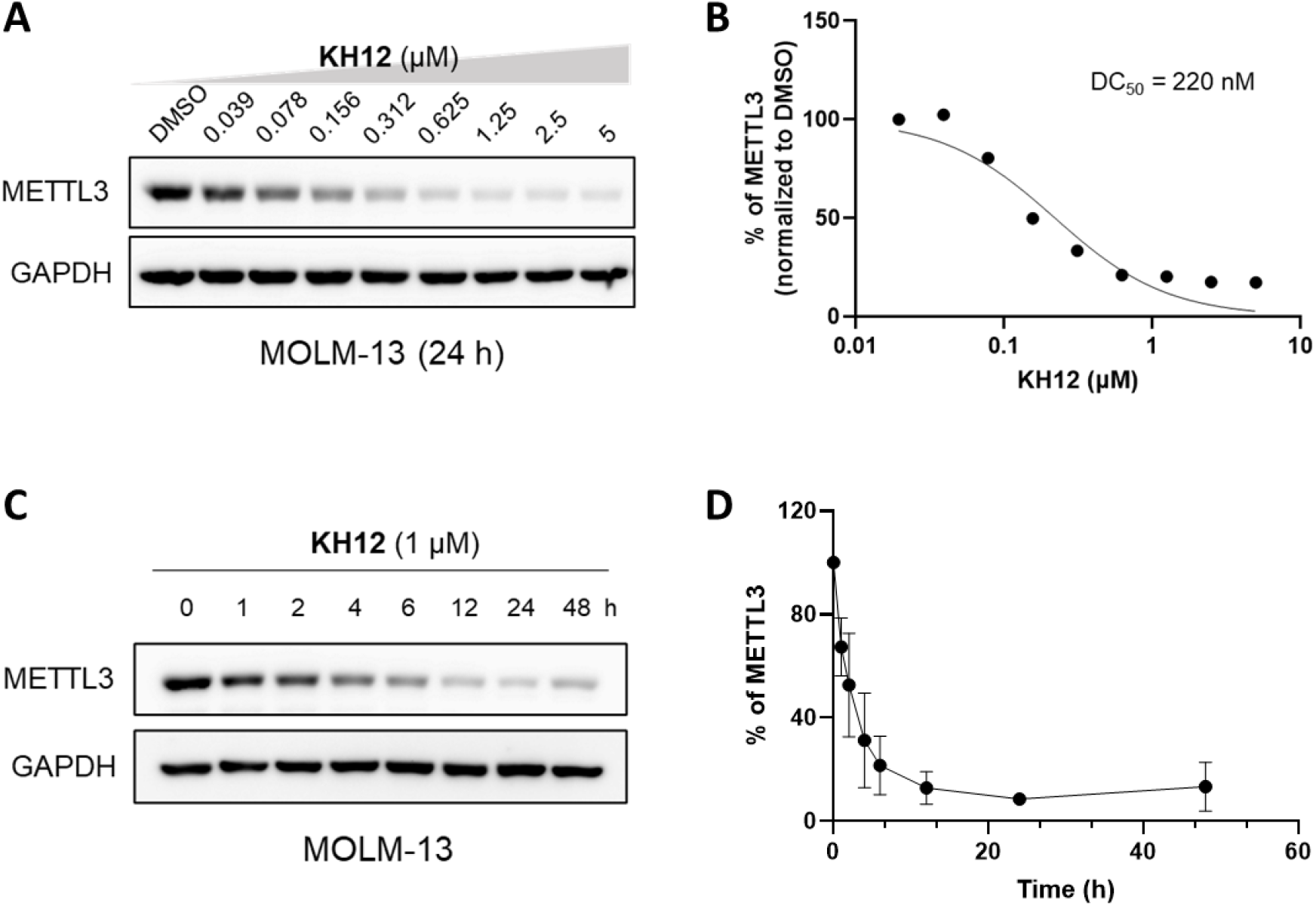
**KH12** induces degradation of METTL3 in dose- and time-dependent manner. A) Indicated concentrations of **KH12** were treated on MOLM-13 cells for 24 h. METTL3 protein levels were detected by western blot. B) **KH12** dose-response curve of METTL3 protein level in MOLM-13 cells. Protein band intensities shown in western blot (A) were quantified. METTL3 protein levels compared to GAPDH were normalized to DMSO control. Individual points are the average of two independent experiments. C) Western blot analysis of METTL3 protein levels in lysates of MOLM-13 cells treated with **KH12 (**1 μM) for indicated time points. D) Time-course curve of METTL3 protein levels in MOLM-13 cells under treatment with **KH12 (**1 μM). Protein band intensities shown in western blot (C) were quantified. METTL3 protein levels compared to GAPDH were normalized to 0 h control. Individual points are the average of two independent experiments. (Each western blot data is representative of two independent replicates)

### Biochemical characterization of KH12

We next investigated target engagement of the hetero-bifunctional molecule **KH12**. We carried out a cellular thermal shift assay (CETSA) to assess **KH12**-METTL3 engagement. Treatment of **KH12** in MOLM-13 cells at 54 °C stabilized endogenous METTL3 in a dose-dependent manner, indicating binding of **KH12** to cellular METLL3 (Figure 3B). In addition, **KH12** possessing VHL ligand was competed with MZ1 on HeLa cells stably expressing GFP-fused BRD4 (Figure 3C). MZ1 has previously been reported as a potent BRD4 PROTAC consisting of VHL ligand^18^. While MZ1 alone effectively reduced BRD4 levels, the co-treatment of MZ1 and **KH12** resulted in the rescue of BRD4 levels depending on dose of **KH12** (Figure 3D and S3). This implies that **KH12** effectively binds to VHL and competes with MZ1 for the VHL binding site.

**Figure 3.**
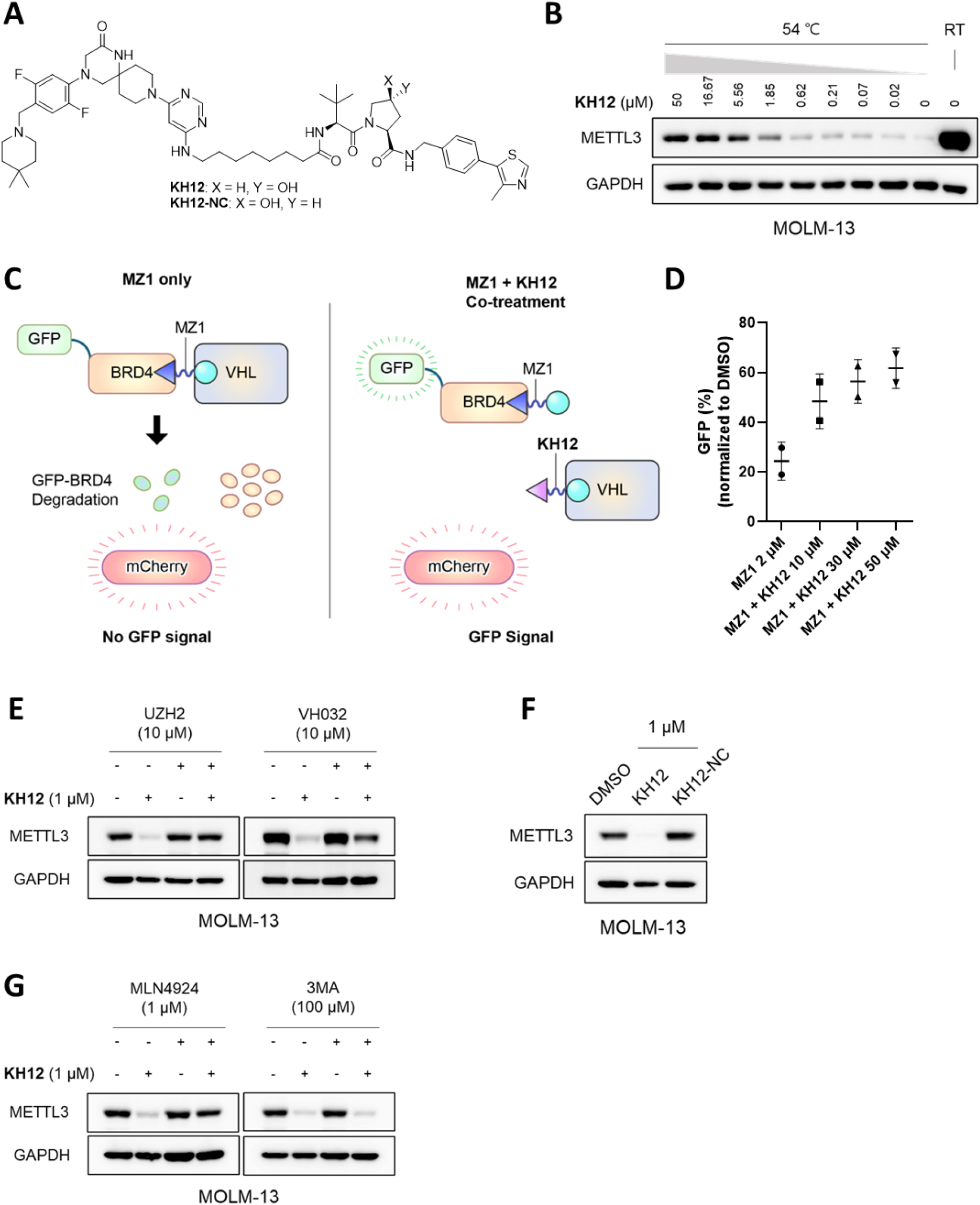
Biochemical characterization of **KH12**. A) Structure of **KH12** and its negative control, **KH12-NC**. B) CETSA at 54 °C in MOLM-13 cells. **KH12** dose-response METTL3 protein levels were detected by western blot. C) Schematic of VHL target engagement assay. D) VHL target engagement assay in BRD4-eGFP_mCherry expressing HeLa cells. Cells were treated either with MZ1 alone (2 μM) or cotreated with indicated concentrations of **KH12**. GFP signals were detected by flow cytometry and normalized to DMSO control. Quantification of n = 2 independent experiments. E) Competition assay using excess amount of UZH2 or VH032 with **KH12**. MOLM-13 cells were pre-treated either with UZH2 (10 μM) or VH032 (10 μM) for 2 h, followed by treatment of **KH12** (1 μM) for 24 h. F) Western blot analysis of METTL3 protein levels in lysates of MOLM-13 cells treated with **KH12** (1 μM) or **KH12-NC** (1 μM) for 24 h. G) Effects of NAE inhibitor and autophagy inhibitor on the **KH12-**induced METTL3 degradation. MOLM-13 cells were pre-treated with MLN4924 (1 μM) or 3MA (100 μM) for 2 h, followed by treatment of **KH12** (1 μM) for 24 h. (Each western blot data is representative of two independent replicates)

We proceeded to explore the mechanism by which **KH12** induces METTL3 degradation in MOLM-13 cells. Degradation activity of **KH12** was remarkably perturbed when either METTL3 inhibitor UZH2 or VHL ligand VH032 was pre-treated (Figure 3E). In addition, we prepared **KH12-NC** as a negative control of **KH12** by conjugating inactive VHL ligand and observed that **KH12-NC** is unable to reduce METTL3 levels on MOLM-13 cells (Figure 3A and F). Taken together, our findings indicate that METTL3 degradation occurs only when both the warhead and VHL ligand of **KH12** are actively engaged, which is necessary for productive ternary complex formation. Moreover, it was found that pre-treatment of Neddylation Activating Enzyme (NAE) inhibitor MLN4924 rescues METTL3 degradation induced by **KH12** while autophagy inhibitor 3MA does not (Figure 3G). This observation implies that **KH12** induces METTL3 degradation via ubiquitin-proteasome system, not by autophagy-lysosomal pathway.

### Anti-proliferative effect of KH12 against AML cells

The importance of m^6^A modification in hematopoietic cell differentiation and the pivotal role of METTL3 in AML cells have previously been reported^9,10^. The use of small molecule inhibitors to suppress the methyltransferase activity of METTL3 has been demonstrated to reduce m^6^A levels and exert anti-proliferative effect against AML cells^14,15^. Establishing the capability of **KH12** to induce METTL3 degradation on MOLM-13 cells, we explored antiproliferative activity of **KH12** on MOLM-13 cells. Compared to METTL3 inhibitors UZH2 and STM2457, **KH12** remarkably suppresses growth of MOLM-13 cells with IC_50_ value of 1.04 μM (Figure 4A). In MOLM-13 cells, translation of c-MYC is promoted by m^6^A modification on its transcripts^9^. It was observed that **KH12** is capable of reducing c-MYC protein levels in a dose-dependent fashion (Figure 4B). The result indicates that **KH12-**induced depletion of METTL3 efficiently reduces the translation of genes reliant on m^6^A modification. Treatment of **KH12** also resulted in a significant induction of myeloid differentiation^10,19^ and apoptosis on MOLM-13 cells (Figure 4C, D, and S4). Overall, it was concluded that trigger degradation of METTL3 is an efficient strategy against MOLM-13 cells. Targeted degradation of METTL3 surpassed the catalytic inhibition in inducing apoptosis and exerting antiproliferative effect, which might be involved with the deletion of m^6^A-independent functions.

**Figure 4.**
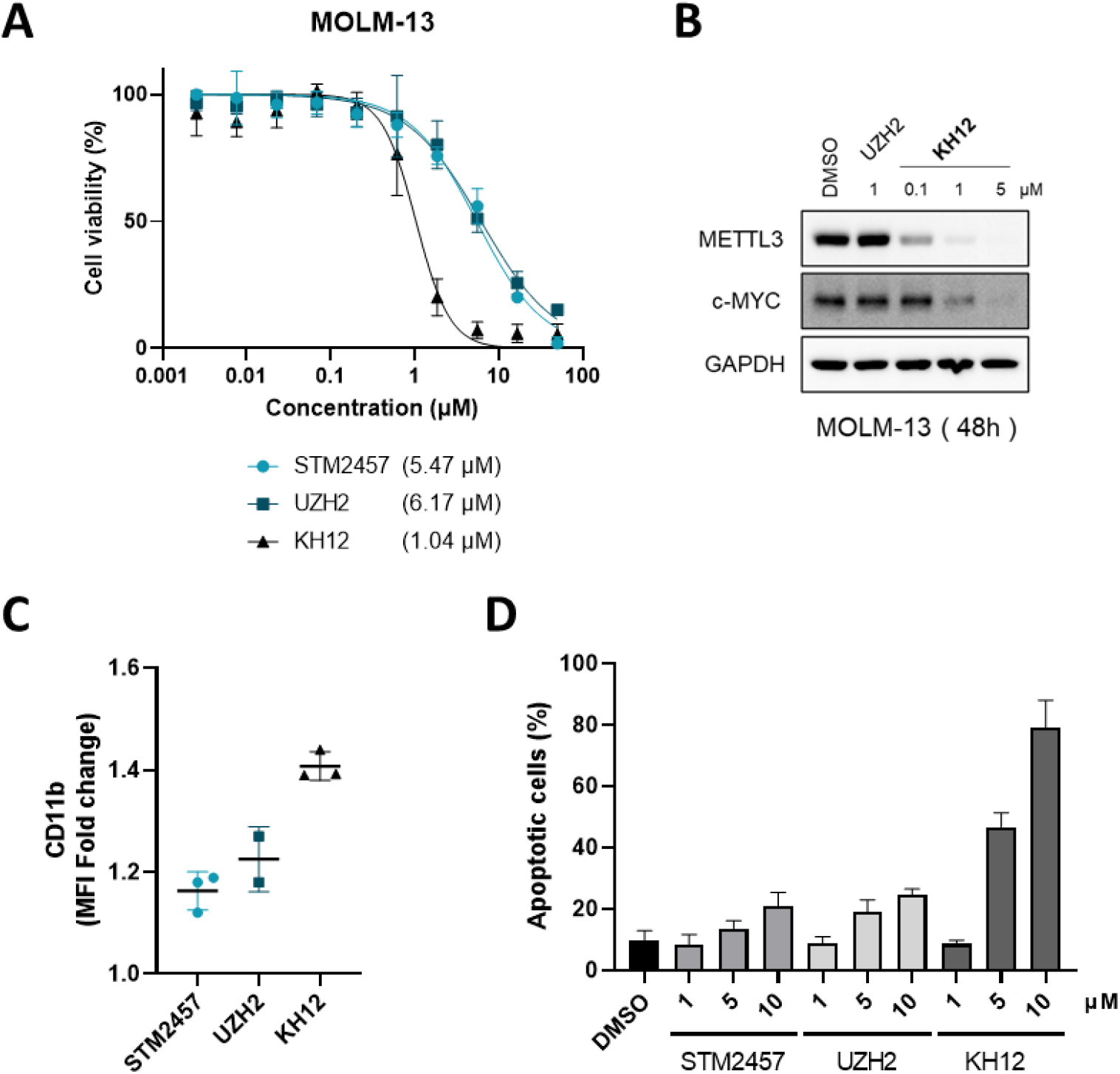
Anti-proliferative effect of **KH12** against MOLM-13 cells. A) Cell viability assays were conducted in MOLM-13 cells treated with STM2457, UZH2, or **KH12** at indicated dose range for 72 h. IC_50_ value for each compound is shown in brackets. The experiments were performed in three independent replicates with duplicate. B) Western blot analysis of METTL3 and c-MYC protein levels in lysates of MOLM-13 cells treated with indicated concentration of UZH2 or **KH12** for 48 h. (Data is representative of two independent replicates) C) MOLM-13 cells were treated with 5 μM of STM2457, UZH2, or **KH12** for 48 h. CD11b levels were assessed by flow cytometry. Quantification of n = 3 independent experiments. D) Apoptosis assay in MOLM-13 cells treated with STM2457, UZH2, or **KH12** at indicated concentration for 48 h. Annexin V and PI staining cells were detected by flow cytometry. Quantification of n = 2 independent experiments.

### Anti-proliferative effect of KH12 against GC cells and organoids

Abnormal cytoplasmic localization of METTL3 and its m^6^A-independent enhancement of the translation have been associated with promotion of tumorigenesis in gastric cancer (GC)^13^. Confirming the capability of **KH12** to induce METTL3 degradation in different cancer cell types including AGS GC cells, we then continued to explore anti-GC potential of **KH12** (Figure 5A and S5). We observed a significant growth-inhibitory activity of **KH12** on AGS cells, which greatly exceeds the effects of small molecule METTL3 inhibitors, STM2457 and UZH2 (Figure 5B). We further investigated effects of **KH12** on different types of GC cells. It is worthwhile to note that **KH12** suppresses growth of most gastric cancer cells explored while small molecule METTL3 inhibitors have little to no effects (Figure 5C and S6). Based on the previous report^13^, it is assumed the anti-proliferative effects of **KH12** on GC cells are associated with m^6^A-independent function of METTL3. Furthermore, we explored anti-GC effects using patient-derived organoids (PDOs) and it was found that **KH12** is also capable of significantly suppressing growth of PDOs. While growth of GA352T organoids was suppressed by both **KH12** and the small molecule inhibitors, GA215T organoids are much more sensitive to **KH12** rather than to the small molecule inhibitors (Figure 5D-G). Overall, our findings suggest that **KH12** efficiently attenuates progression of GC in both cell and organoid levels. We assume anti-GC effect of **KH12** is mainly associated with the deletion of m^6^A-independent function of METTL3.

**Figure 5.**
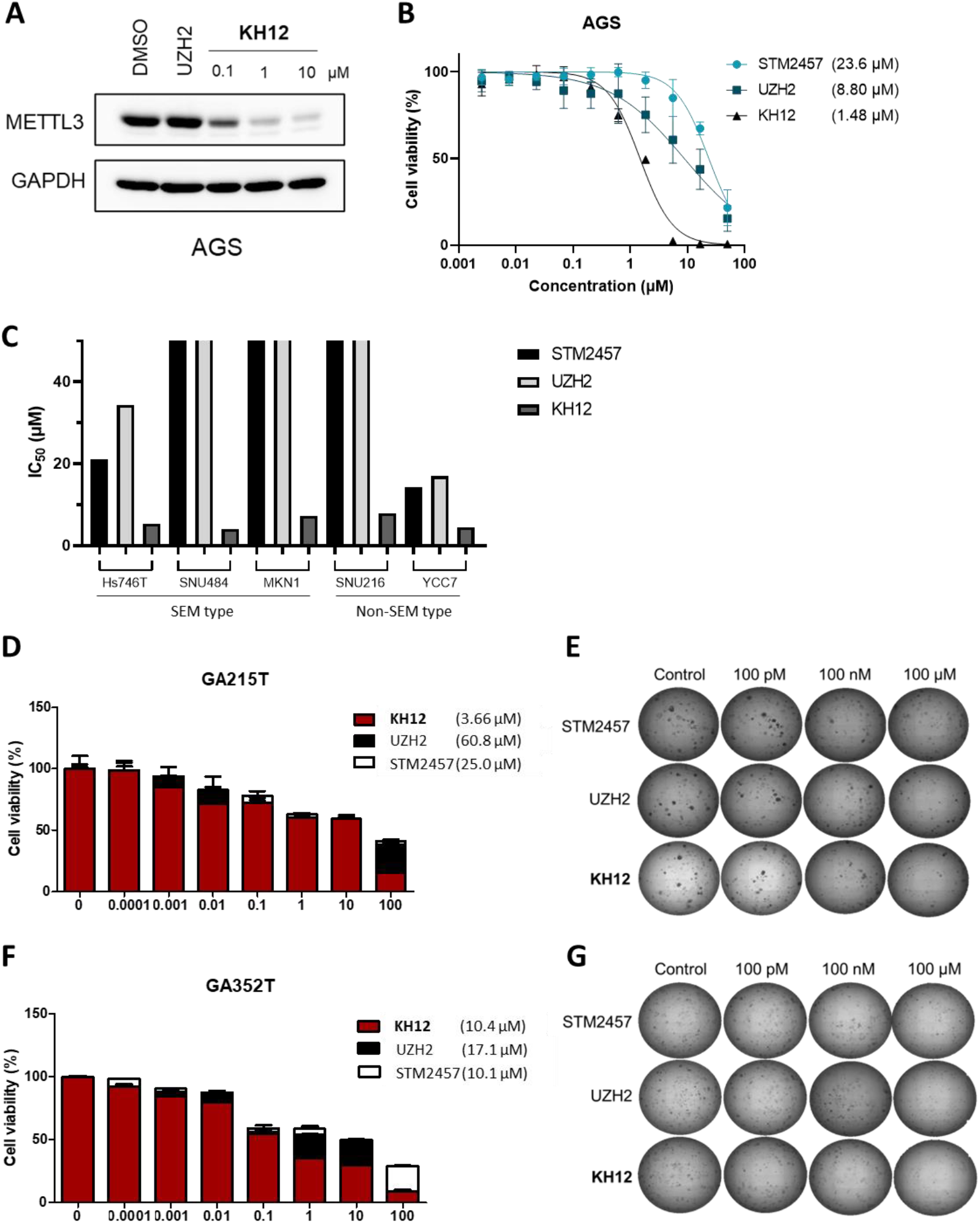
Anti-GC effects of **KH12** A) Western blot analysis of METTL3 protein levels in lysates of AGS cells treated with indicated concentration of UZH2 or **KH12** for 24 h. (Data is representative of two independent replicates) B) Cell viability assay in AGS cells treated with STM2457, UZH2, or **KH12** at indicated does range for 72 h. IC_50_ value for each compound is shown in brackets. The experiments were performed in two independent replicates with duplicate. C) Cell viability assay in GC cell lines treated with STM2457, UZH2, or **KH12** for 72 h. Each analysis was performed in two independent replicates with triplicate and IC_50_ values of compounds are indicated with bar graph. Organoid viability assay in GA215T (D) and GA352T (F) treated with STM2457, UZH2, or **KH12** at indicated does range for 5 days. IC_50_ value for each compound is shown in brackets. Organoid morphology at indicated concentration in GA215T (E) and GA352T (G).

## DISCUSSION

Through SAR study with two series of PROTACs, we have successfully identified VHL-based PROTAC **KH12** as a potent METTL3 degrader. **KH12** has DC_50_ value of 220 nM in MOLM-13 cells and inhibits METTL3 enzyme with IC_50_ value of 341 nM in biochemical assay (Figure S7). Based on our SAR study, it was found that degradation activities of METTL3 PROTACs are significantly affected by composition and length of linker moiety and the aliphatic linker composed of seven-methylenes in **KH12** is optimal for degradation activity. Coupled with the rapid induction of **KH12** for METTL3 degradation, it was hypothesized that VHL-mediated METTL3 degradation undergoes in a highly efficient way that only a modest recruitment of METTL3 to VHL would be sufficient for degradation. Nevertheless, we expect further optimization of the linker should improve the degradation activity of **KH12**. On the other hand, it was found that the CRBN-based PROTAC series induces limited degradation of METTL3 relative to VHL-based PROTACs. Consistent with the report by Caflsich group^16^, it was assumed that CRBN E3 ligase may not be optimal for METTL3 degradation.

Upon treatment of **KH12** on MOLM-13 cells, it was observed that an equivalent level of METTL14 reduction along with METTL3 degradation (Figure S8). It was reported that METTL3 stabilizes METTL14 and STUB1 mediates METTL14 degradation in the absence of METTL3^20^. However, we lack insight into whether METTL3/14 complex is degraded altogether by the PROTAC or if METTL14 degradation is a consequent result of destabilization by METTL3 depletion. In the METTL3/14 complex, METTL14 provides an RNA-binding platform, facilitates recognition of RNA substrate, and thereby promotes the methyltransferase activity of METTL3^21^. Although METTL14 alone has no catalytic activity, the deletion of METTL14 might contribute to the cellular effects of **KH12**. However, the study presented here did not investigate any direct effect of METTL14 deletion.

SEM (Stem-like/Epithelial-to-mesenchymal transition/Mesenchymal) type GC is the most aggressive subtype with no targeted therapies^22^. It is of note that progression of both SEM-type and non-SEM-type GC cells was remarkably suppressed by **KH12** (Figure 5C and S6). Despite the high heterogeneity of GC, it is anticipated that targeted degradation of METTL3 offers therapeutic opportunity to treat GC including its most aggressive subtypes. Also, it is hypothesized that anti-GC effects of **KH12** are contributed by the deletion of cytoplasmic METTL3. Abnormal cytoplasmic METTL3 is implicated with GC progression by complexing with translation initiation factors and enhancing translation of epigenetic factors independent of its methyltransferase function^13^. Nevertheless, additional investigations are required to elucidate the mechanism by which **KH12** perturbs the progression of GC.

Herein, we designed and synthesized METTL3 PROTACs by utilizing either VHL or CRBN E3 ligase. We demonstrated that **KH12** suppressed growth of MOLM-13 AML cells, induced differentiation and apoptosis. Moreover, we observed its anti-proliferative effects on various types of GC cells and organoids. While transient inhibition of METTL3 catalytic function by small molecule inhibitors has little effect on growth of GC cells, deletion of METTL3 by **KH12** results in strongly impeded progression of GC, which is an agreement with the previous report^13^. Altogether, this study provides insight into design of METTL3 PROTAC and therapeutic potentials of targeted degradation of METTL3.

## Acknowledgements

This research was supported by the following grants: the Brain Korea 21 FOUR Project for Medical Science, Yonsei University College of Medicine, Seoul, Republic of Korea and NRF grant funded by the Korean government (MSIP) (NRF 2018R1A5A2025079). We thank Dr. Ha-Soon Choi, Magicbullettherapeutics Inc. for the advice on synthesis. We thank Prof. Yongchul Kim, Gwangju Institute of Science and Technology (GIST) for generously providing MOLM-13 cells.

## References

1 He, P. C. & He, C. m(6) A RNA methylation: from mechanisms to therapeutic potential. EMBO J 40, e105977 (2021).

2 Livneh, I., Moshitch-Moshkovitz, S., Amariglio, N., Rechavi, G. & Dominissini, D. The m(6)A epitranscriptome: transcriptome plasticity in brain development and function. Nat Rev Neurosci 21, 36–51 (2020).

3 Zhang, M., Zhai, Y., Zhang, S., Dai, X. & Li, Z. Roles of N6-Methyladenosine (m(6)A) in Stem Cell Fate Decisions and Early Embryonic Development in Mammals. Front Cell Dev Biol 8, 782 (2020).

4 Schaefer, M. R. The Regulation of RNA Modification Systems: The Next Frontier in Epitranscriptomics? Genes (Basel) 12 (2021).

5 Bokar, J. A., Shambaugh, M. E., Polayes, D., Matera, A. G. & Rottman, F. M. Purification and cDNA cloning of the AdoMet-binding subunit of the human mRNA (N6-adenosine)-methyltransferase. RNA 3, 1233–1247 (1997).

6 Liu, J. et al. A METTL3-METTL14 complex mediates mammalian nuclear RNA N6-adenosine methylation. Nat Chem Biol 10, 93–95 (2014).

7 Lan, Q. et al. The Critical Role of RNA m(6)A Methylation in Cancer. Cancer Res 79, 1285–1292 (2019).

8 Xu, P. & Ge, R. Roles and drug development of METTL3 (methyltransferase-like 3) in anti-tumor therapy. Eur J Med Chem 230, 114118 (2022).

9 Vu, L. P. et al. The N(6)-methyladenosine (m(6)A)-forming enzyme METTL3 controls myeloid differentiation of normal hematopoietic and leukemia cells. Nat Med 23, 1369–1376 (2017).

10 Barbieri, I. et al. Promoter-bound METTL3 maintains myeloid leukaemia by m(6)A-dependent translation control. Nature 552, 126–131 (2017).

11 Choe, J. et al. mRNA circularization by METTL3-eIF3h enhances translation and promotes oncogenesis. Nature 561, 556–560 (2018).

12 Lin, S., Choe, J., Du, P., Triboulet, R. & Gregory, R. I. The m(6)A Methyltransferase METTL3 Promotes Translation in Human Cancer Cells. Mol Cell 62, 335–345 (2016).

13 Wei, X. et al. METTL3 preferentially enhances non-m(6)A translation of epigenetic factors and promotes tumourigenesis. Nat Cell Biol 24, 1278–1290 (2022).

14 Dolbois, A. et al. 1,4,9-Triazaspiro[5.5]undecan-2-one Derivatives as Potent and Selective METTL3 Inhibitors. J Med Chem 64, 12738–12760 (2021).

15 Yankova, E. et al. Small-molecule inhibition of METTL3 as a strategy against myeloid leukaemia. Nature 593, 597–601 (2021).

16 Francesco Errani, A. I., Marcin Herok, Elena Bochenkova, Fiona Stamm, Ivan Corbeski, Valeria Romanucci, Giovanni Di Fabio, František Zálešák, Amedeo Caflisch PROTAC degraders of the METTL3-14 m6A-RNA methyltransferase. ChemRxiv (2023).

17 Du, W. et al. Discovery of a PROTAC degrader for METTL3-METTL14 complex. Cell Chem Biol (2024).

18 Zengerle, M., Chan, K. H. & Ciulli, A. Selective Small Molecule Induced Degradation of the BET Bromodomain Protein BRD4. ACS Chem Biol 10, 1770–1777 (2015).

19 Ripperger, T. et al. The heteromeric transcription factor GABP activates the ITGAM/CD11b promoter and induces myeloid differentiation. Biochim Biophys Acta 1849, 1145–1154 (2015).

20 Zeng, Z. C. et al. METTL3 protects METTL14 from STUB1-mediated degradation to maintain m(6) A homeostasis. EMBO Rep 24, e55762 (2023).

21 Liu, X. et al. Insights into roles of METTL14 in tumors. Cell Prolif 55, e13168 (2022).

22 Yoon, B. K. et al. PHGDH preserves one-carbon cycle to confer metabolic plasticity in chemoresistant gastric cancer during nutrient stress. Proc Natl Acad Sci U S A 120, e2217826120 (2023).

