## Supplementary information for "Targeted Degradation of METTL3 Against Acute Myeloid Leukemia and Gastric Cancer"

**Supporting Information**

**Table of Contents:**

Tables and Figures	**-** S1

Chemistry	**-** S8

Biology and Biochemistry Methods	**-** S23

^1^H and ^13^C NMR Spectra of Intermediates	**-** S26

^1^H and ^13^C NMR Spectra of METTL3 PROTACs	**-** S45

HPLC Traces of METTL3 PROTACs	**-** S58

ESI-HRMS Spectra of METTL3 PROTACs	**-** S71

References	**-** S75

**Tables and Figures**

**Table S1.** METTL3 degradation activity for PROTACs.

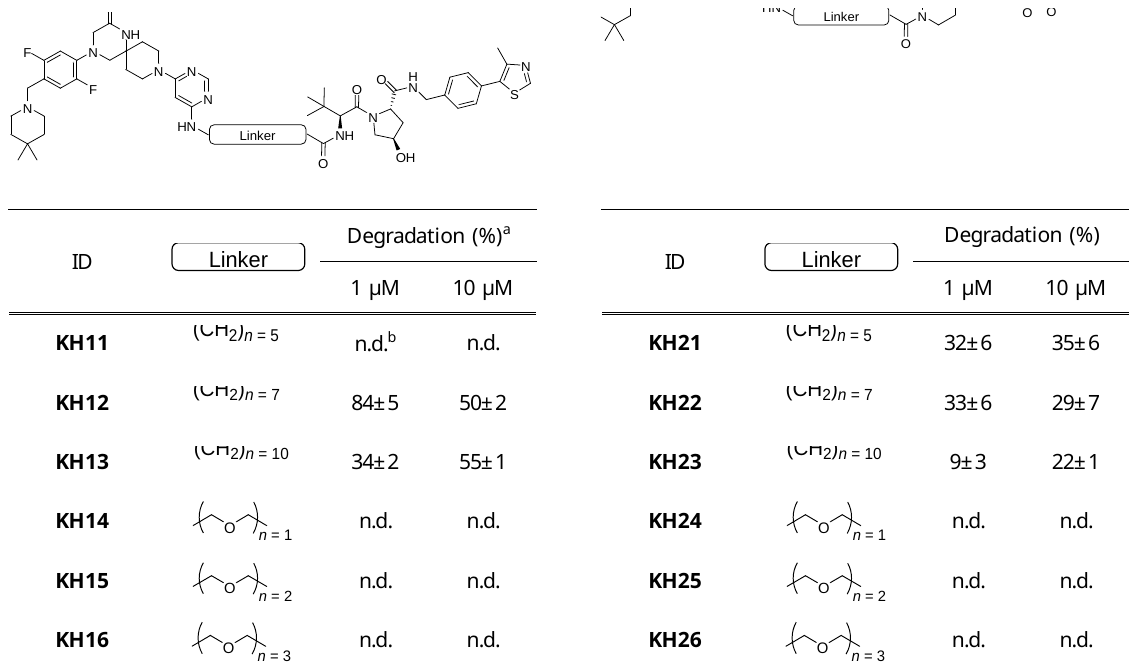

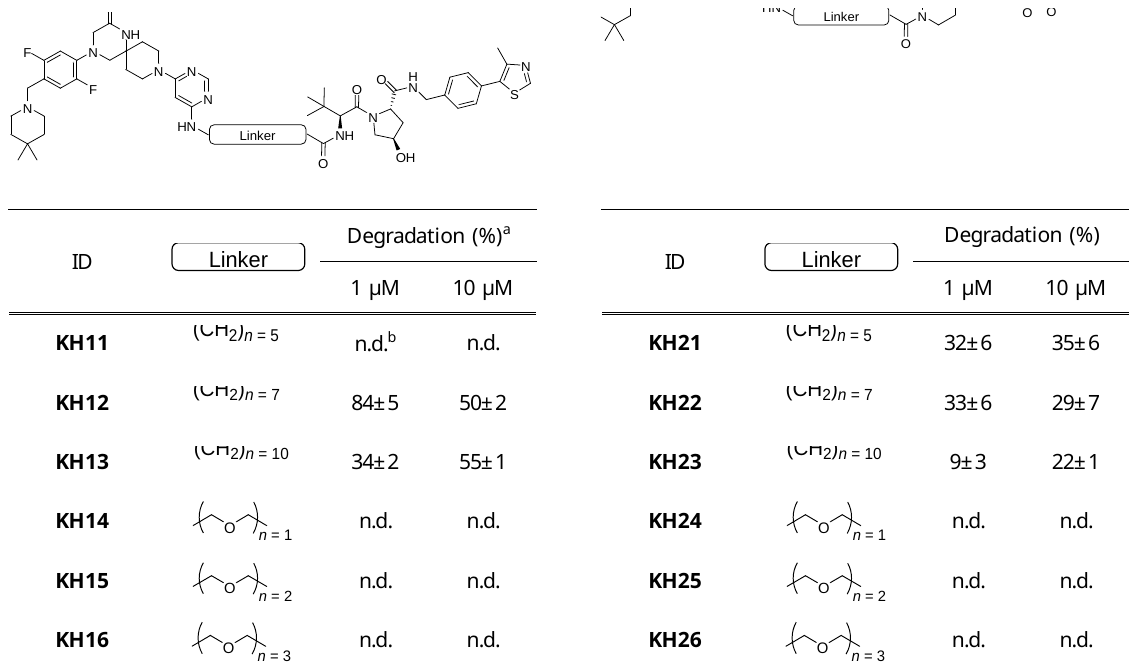

^a^ Protein band intensities shown in western blot of Figure S1 were quantified. METTL3 protein levels compared to GAPDH were normalized to DMSO control. Degradation percentage of METTL3 protein level is presented as mean ± SD, n = 2 independent experiments. ^b^ No degradation (n.d.)

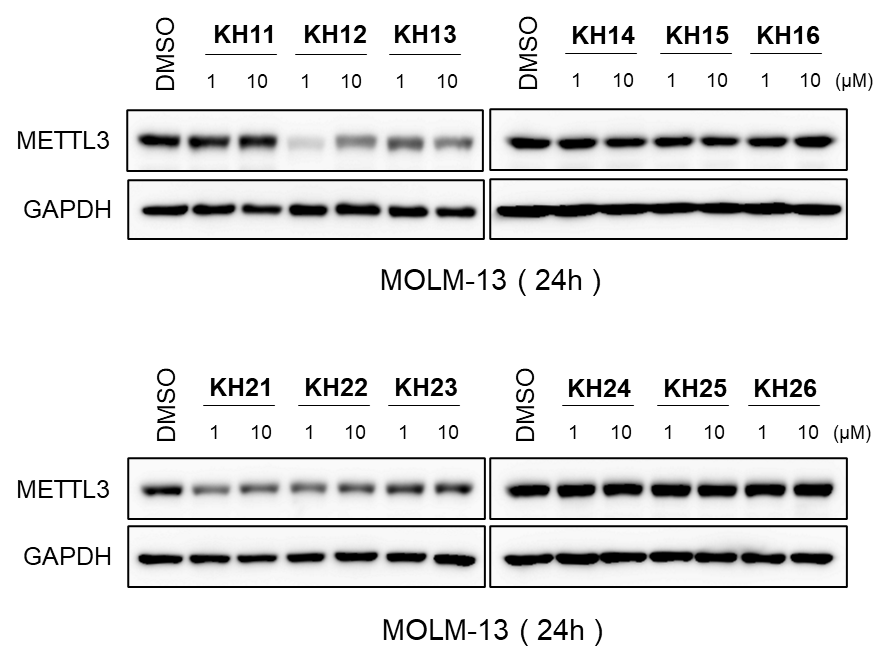

**Figure S1.** METTL3 degradation activity for PROTACs.

Western blot analysis of METTL3 protein levels in lysates of MOLM-13 cells treated with

indicated dose of PROTAC compounds for 24 h. (Each western blot data is representative of two independent replicates)

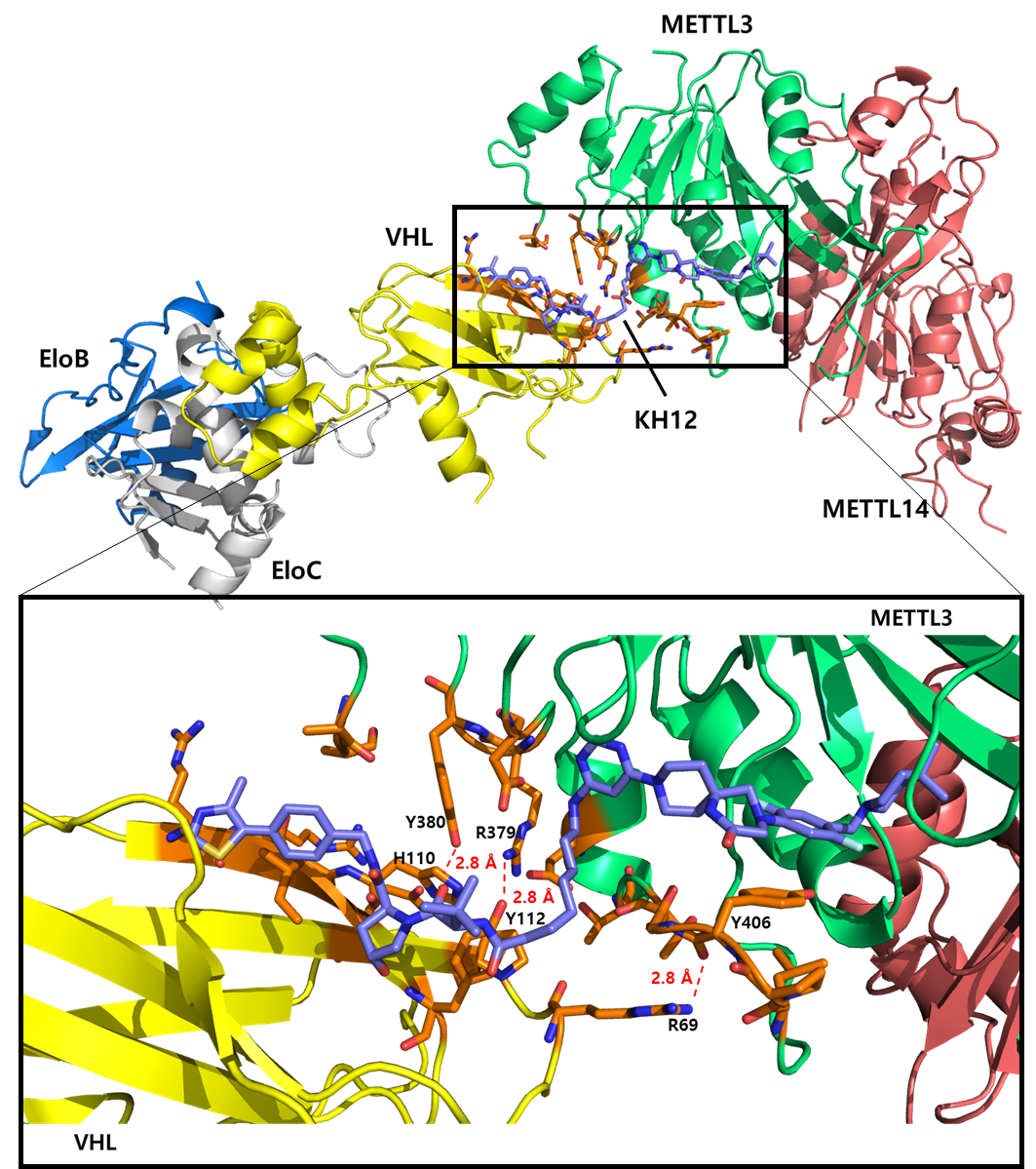

**Figure S2.** Predicted model of EloB/C-VHL-**KH12**-METTL3/14 complex.

Docking studies were performed with ICM-Pro using structures PDB: 5LLI and PDB: 7O2F and further prepared using PyMOL.

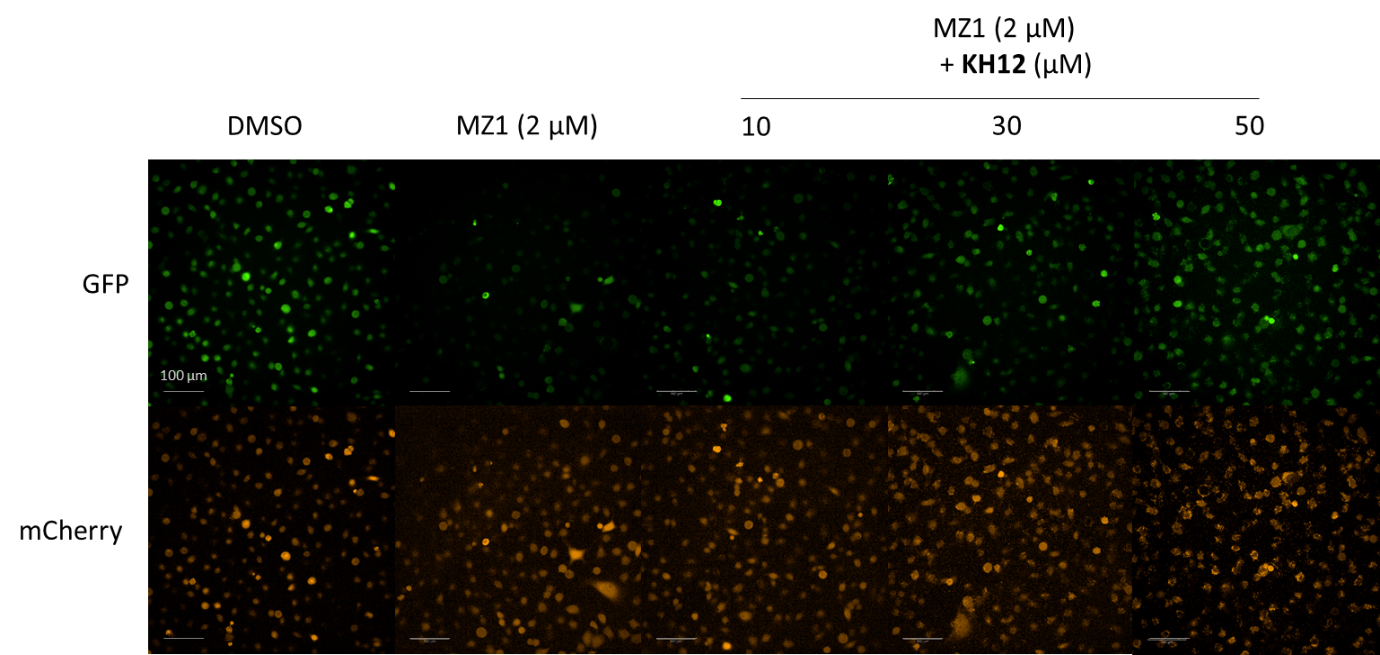

**Figure S3.** VHL target engagement assay in BRD4-GFP_mCherry transduced Hela cells.

Cells were treated either with MZ1 alone (2 μM) or co-treated with indicated concentration of **KH12**. GFP and mCherry signals were detected by HCS imaging. (Data is representative of two independent replicates)

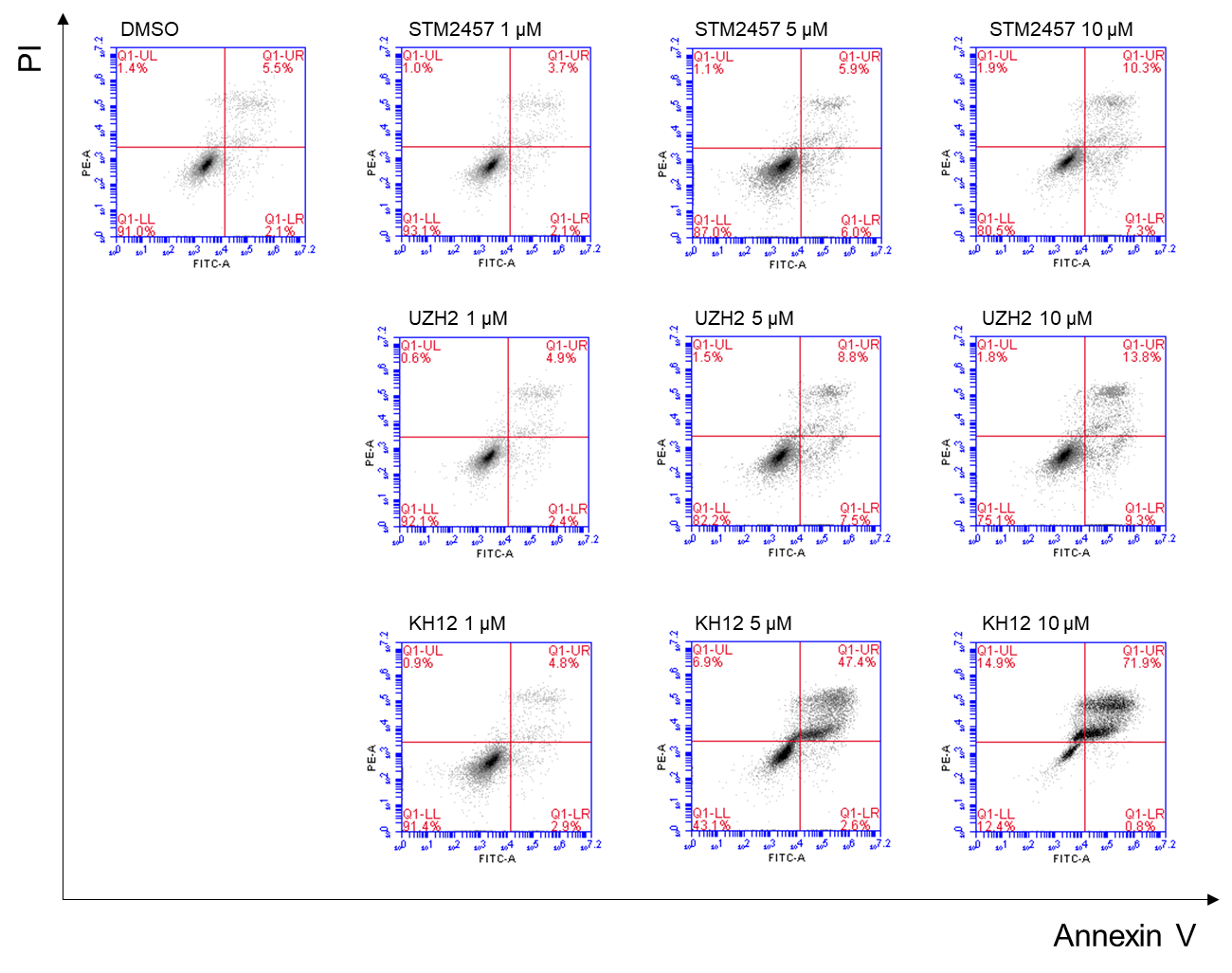

**Figure S4.** Apoptosis assay in MOLM-13 cells.

MOLM-13 cells treated with STM2457, UZH2, or **KH12** at indicated concentration for 48 h. Annexin V and PI staining cells were detected by flow cytometry. (Data is representative of two independent replicates)

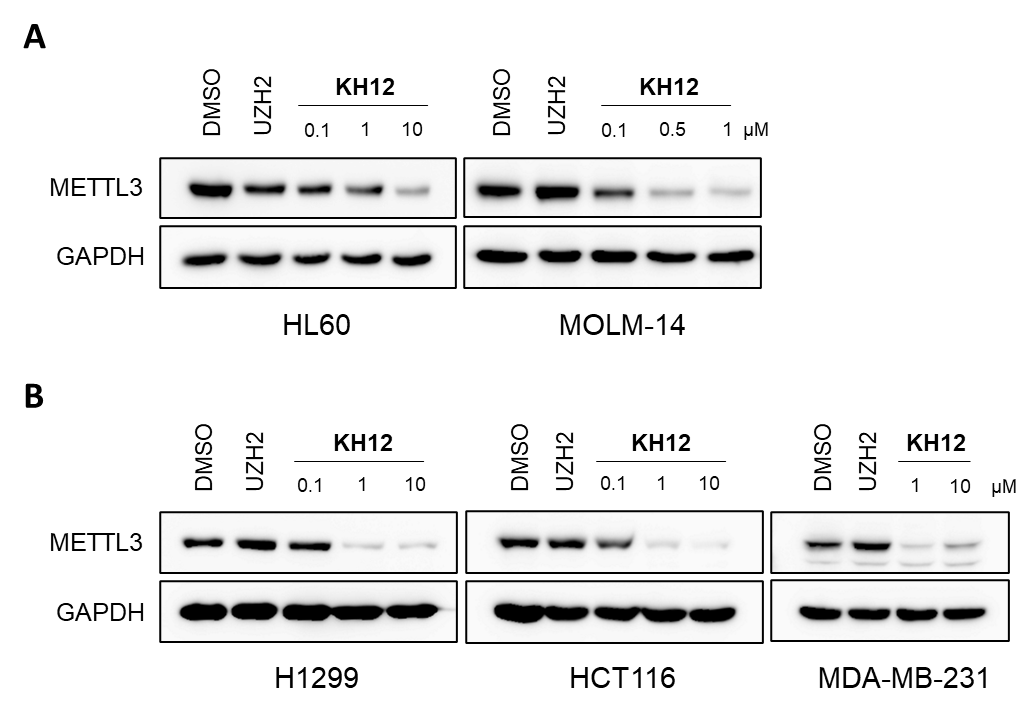

**Figure S5.** **KH12** efficiently induces METTL3 degradation in different types of cancer cells.

Western blot analysis of METTL3 protein levels in lysates of A) AML cell lines, HL60 and MOLM-14 cells or B) H1299 (lung cancer), HCT116 (colorectal cancer) and MDA-MB-231 (breast cancer) cells treated with UZH2 or indicated concentration of **KH12** for 24 h. (Data is representative of two independent replicates)

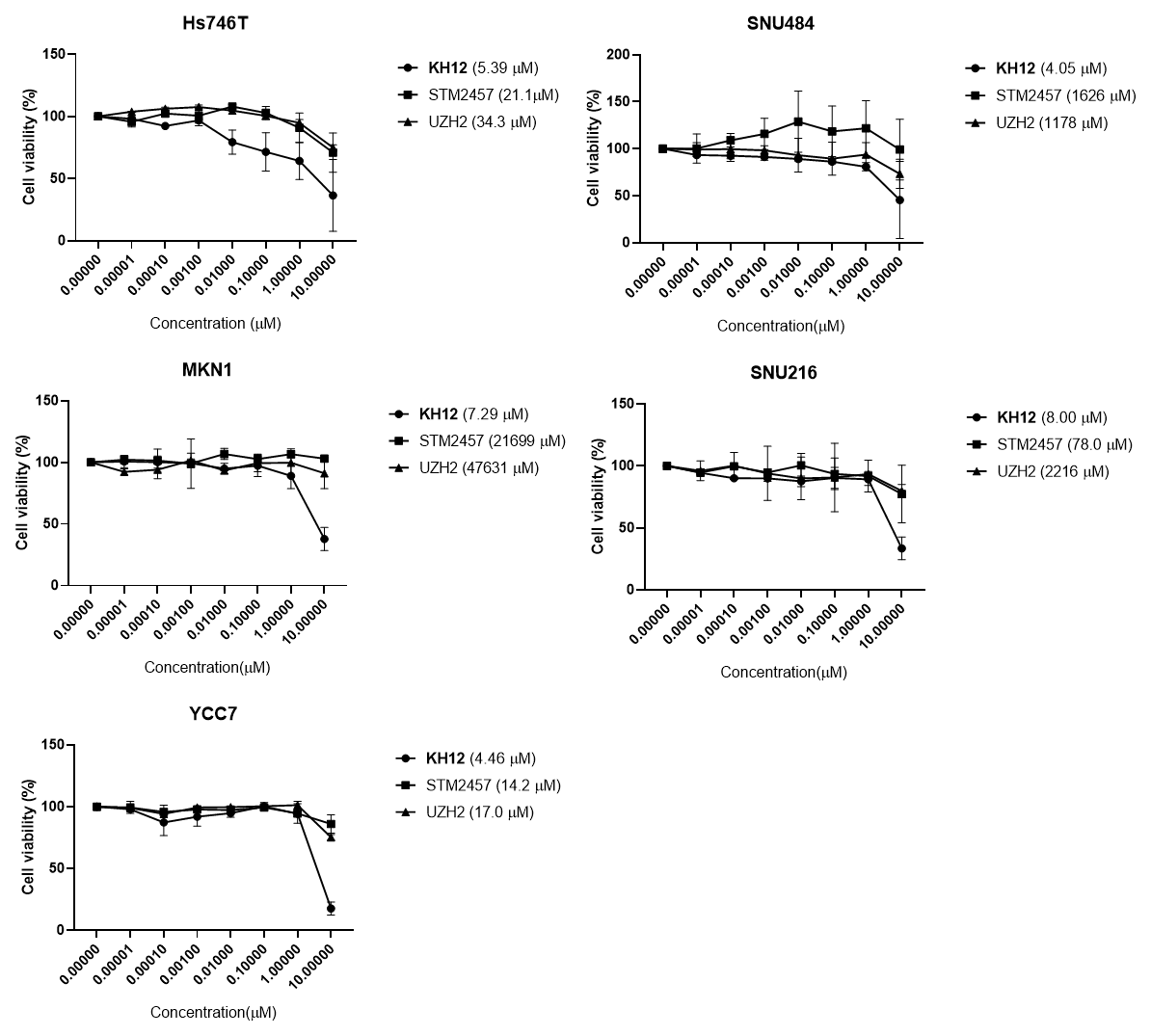

**Figure S6.** GC cell viability assay.

Cell viability assay in GC cell lines treated with STM2457, UZH2, or **KH12** at indicated dose range for 72 h. Each analysis was performed in two independent replicates with triplicate and IC_50_ values of each compound are shown in brackets.

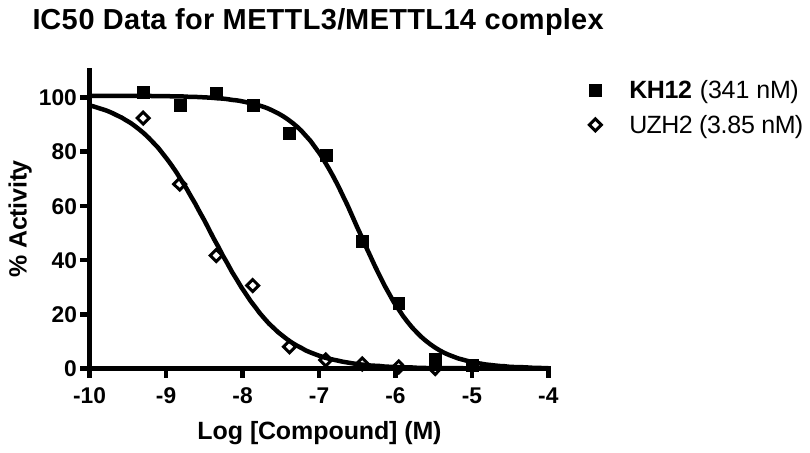

**Figure S7.** Methyltransferase assay showing inhibition of the METTL3/14 complex using a dose-range of UZH2 and **KH12**.

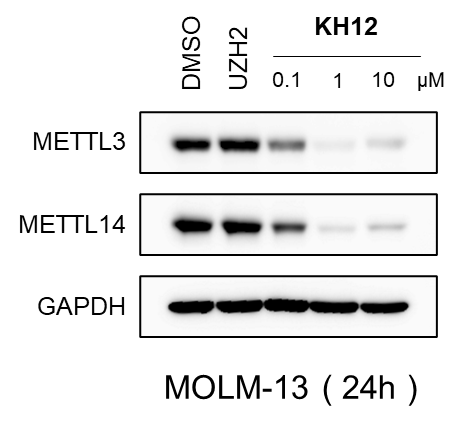

**Figure S8. KH12** induces METTL14 degradation along with METTL3 degradation.

Western blot analysis of METTL3 and METTL14 protein levels in lysates of MOLM-13 cells treated with 1 μM of UZH2 or indicated concentration of **KH12** for 24 h. (Data is representative of two independent replicates)

**Chemistry**

**General information.** Unless otherwise described, all reagents and solvents were purchased from commercial suppliers and used without further purification. All reactions were performed in flame-dried glassware under N_2_ atmosphere. Reactions were monitored by using TLC with 0.25 mm E. Merck pre-coated silica gel plates (60 F254). TLC was analyzed with UV, ninhydrin, or *p*-anisaldehyde stain for detection. Purification of reaction mixture was carried out by using silica gel column chromatography with Kieselgel 60 Art. 9385 (230–400 mesh) or by using reverse phase column chromatography with RediSep Gold® C18 Reversed Phase Columns (20-40 *μ*m particle size). Purities of all compounds were ≥ 95%. Mass spectra and purities of all compounds were assessed using LC/MS analysis with Waters LC/MS system (Waters 2998 Photodiode Array Detector, Waters 3100 Mass Detector, Waters SFO System Fluidics Organizer, Water 2545 Binary Gradient Module, Waters Reagent Manager, and Waters 2767 Sample Manager) using SunFire^TM^ C18 column (4.6 × 50 mm, 5 *μ*m particle size): solvent gradient = 30% B at 0.00 min, 100% B at 7.00 min, 100% B at 8.50 min, 30% B at 8.51 min, 30% B at 10.00 min. Solvent A = 0.1% formic acid in H_2_O; Solvent B = 0.1% formic acid in methanol; flow rate = 0.8 mL/min. ^1^H and ^13^C NMR spectra were obtained using Bruker 400 MHz FT-NMR (400 MHz for ^1^H, and 100 MHz for ^13^C) spectrometer. Standard abbreviations are used for denoting the signal multiplicities.

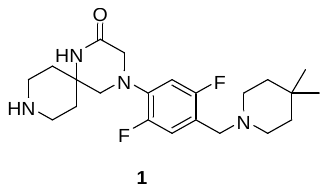

Compound **1** were prepared according to the known literature procedures^1^.

Synthesis of focused series of METTL3 PROTACs are outlined in **Scheme 1-3**. For the synthesis of PROTACs, we prepared UZH2 warhead incorporating ester linkers by employing two sequential S_N_Ar reactions of 4,6-dichloropyrimidine (**2**) (**Scheme 1 and 3**). Esters were further hydrolyzed to carboxylic acid and following HATU-mediated amide coupling with the VHL or CRBN ligands provided METTL3 PROTACs (**Scheme 3**).

The synthetic route begins with generating chloropyrimidine linker blocks. Synthesis of alkyl linker blocks **6-8** and PEG linker blocks **12**-**14** are depicted in **Scheme 1**. For the preparation of alkyl linker blocks, commercially available amines with carboxylic acid ends were subjected to undergo S_N_Ar reaction with 4,6-dichloropyrimidine (**2**). Following Fischer esterification of carboxylic acids provided intermediates **3-5** (62-78%), and subsequent acetylation of amine groups using acetic anhydride produced alkyl linker blocks **6-8** (82-88%). S_N_Ar reaction of **2** with commercially available amino-PEG-*tert*-butyl esters, followed by acetylation provided PEG linker blocks **12-14** (84-86%).

The Synthesis of piperazine-installed CRBN binder is outlined in **Scheme 2**. Aromatic halides of **15** were substituted by *tert*-butyl piperazine-1-carboxylate and following Boc deprotection yielded **17** (94%)

The preparation of METTL3 degraders and the negative compound is shown in **Scheme 3**. Secondary S_N_Ar reactions of chloropyrimidine linker blocks **6-8** and **12-14** with compound **1** were carried out to generate linker-installed METTL3 binders **18-20** and **21-23** (65-75%). Hydrolysis of the ester ends and acetyl deprotection of the amine groups were simultaneously achieved through either a base or acid treatment. The resulting carboxylic acid functionalized METTL3 binders were coupled with VHL or CRBN ligands by amidation reaction to provide METTL3 PROTACs.

**Scheme 1.** Synthesis of chloropyrimidine linker blocks **6-8** and **12**-**14**.

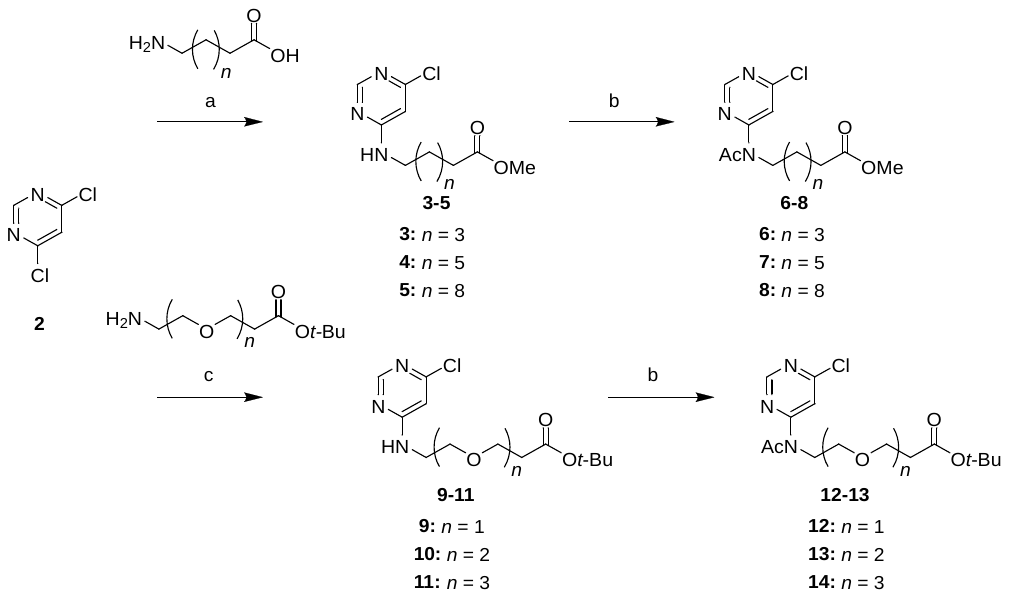

Reagents and conditions: (a) i) Corresponding amines, Et_3_N, EtOH, 70 ^o^C, 12 h; ii) H_2_SO_4_, MeOH, 70 ^o^C, 3 h, 62-78% over two steps; (b) Et_3_N, Ac_2_O, 100 °C, 16 h, 82-88%; (c) Corresponding amines, Et_3_N, EtOH, 70 ^o^C, 12 h, 74-78%.

**Scheme 2.** Synthesis of CRBN E3 ligase binder **17**.

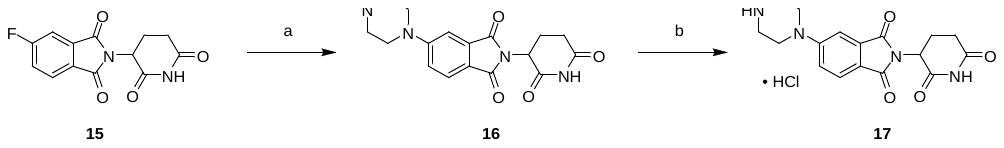

Reagents and conditions: (a) *tert*-butyl piperazine-1-carboxylate, DIPEA, DMSO, 90 ^o^C, 6 h, 94%; (b) HCl, 1,4-dioxane, rt, 3 h, 94%.

**Scheme 3.** Synthesis of METTL3 PROTACs.

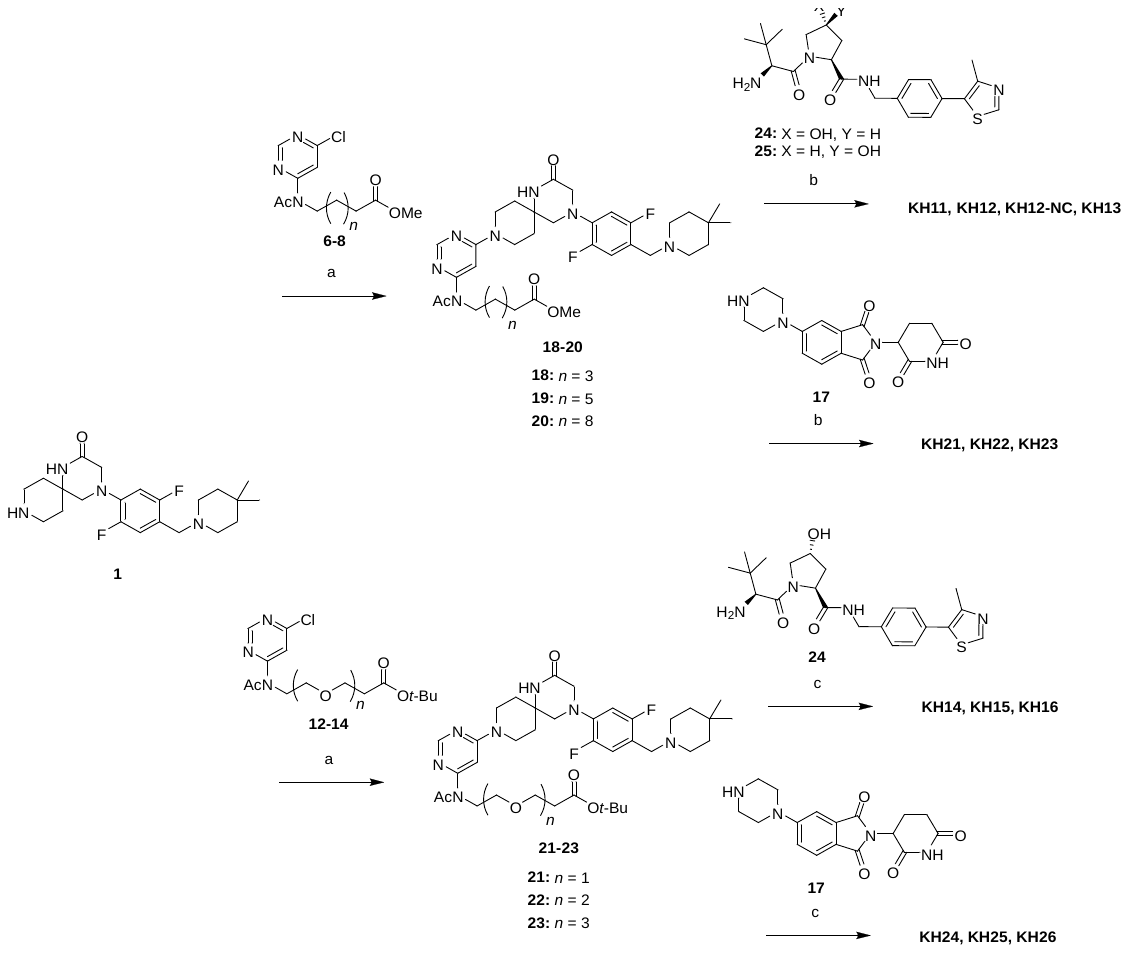

Reagents and conditions: (a) **1**, Chloropyrimidines, Et_3_N, DMF, 70 ^o^C, 5 h, 65-75%; (b) i) LiOH, THF/H_2_O, 50 ^o^C, 16 h; ii) Corresponding amine, HATU, DIPEA, rt, 3 h, 23-36% over two steps; (c) i) HCl, MeOH/1,4-dioxane, 50 ^o^C, 16 h; ii) Corresponding amine, HATU, DIPEA, rt, 3 h, 18-40% over two steps.

**General Procedure A for S_N_Ar with 4,6-Dichloropyrimidine.**

To a solution of the corresponding amine (1.0 equiv) in EtOH (0.5 M), Et_3_N (3.0 equiv) and 4,6-dichloropyrimidine (1.5 equiv) were added. The reaction mixture was stirred at room temperature for 12 h, quenched by addition of saturated aqueous NH_4_Cl solution, and extracted with ethyl acetate. The combined organic layers were washed with brine, dried over Na_2_SO_4_, filtered, and concentrated under reduced pressure. The obtained residues were either subjected to flash column chromatography on silica gel or advanced to the next step without further purification.

**General Procedure B for Esterification.**

To a solution of the corresponding carboxylic acid (1.0 equiv) in MeOH (0.5 M), surfuric acid (98%, 0.2 equiv) was added dropwise at room temperature. The reaction mixture was stirred at 70 ^o^C for 3 h, cooled to 0 ^o^C, and saturated aqueous NaHCO_3_ solution was gently added. The mixture was then extracted with ethyl acetate, washed with brine, dried over Na_2_SO_4_, filtered, and concentrated under reduced pressure. The obtained residues were subjected to flash column chromatography on silica gel.

**General Procedure C for Acetylation.**

To a solution of the corresponding amines (1.0 equiv) in acetic anhydride (0.5 M), Et_3_N (3.0 equiv) was added. The reaction mixture was stirred at 110 ^o^C for 16 h, cooled to 0 ^o^C and methanol and a saturated aqueous NaHCO_3_ solution were gently added. The mixture was stirred for 30 min at room temperature and extracted with ethyl acetate. The combined organic layers were washed with brine, dried over Na_2_SO_4_, filtered, and concentrated under reduced pressure. The obtained residues were subjected to flash column chromatography on silica gel.

Methyl 6-((6-chloropyrimidin-4-yl)amino)hexanoate (**3**).

6-aminohexanoic acid (2.00 g, 15.3 mmol) was converted to compound **3** using general procedure A followed by general procedure B. The crude mixture was purified by flash column chromatography on silica gel (0% to 30% ethyl acetate/hexane) to afford **3** (2.43 g, 9.43 mmol, 62% over two steps.) as a white solid. ^1^H NMR (400 MHz, Methanol-*d*_4_) δ 8.20 (s, 1H), 6.46 (s, 1H), 4.90 (s, 1H), 3.64 (s, 3H), 3.41 – 3.13 (m, 2H), 2.32 (t, *J* = 7.4 Hz, 2H), 1.68 – 1.54 (m, 4H), 1.43 – 1.27 (m, 2H); ^13^C NMR (101 MHz, Methanol-*d*_4_) δ 175.66, 164.82, 159.30, 158.23, 105.12, 51.99, 41.57, 34.60, 29.78, 27.38, 25.64. LRMS (ESI) *m/z* calculated for C_11_H_17_ClN_3_O_2_^+^ [M + H]^+^: 258.1. Found: 258.

Methyl 8-((6-chloropyrimidin-4-yl)amino)octanoate (**4**).

8-aminooctanoic acid (2.00 g, 12.6 mmol) was converted to compound **4** using general procedure A followed by general procedure B. The crude mixture was purified by flash column chromatography on silica gel (0% to 30% ethyl acetate/hexane) to afford **4** (2.82 g, 9.87 mmol, 78% over two steps.) as a white solid. 1H NMR (400 MHz, Methanol-d4) δ 8.20 (s, 1H), 6.46 (s, 1H), 4.90 (s, 1H), 3.63 (s, 3H), 3.40 – 3.10 (m, 2H), 2.29 (t, J = 7.5 Hz, 2H), 1.66 – 1.50 (m, 4H), 1.40 – 1.24 (m, 6H); ^13^C NMR (101 MHz, Methanol-*d*_4_) δ 175.70, 164.77, 159.29, 158.19, 105.11, 51.96, 41.75, 34.68, 30.05, 30.02, 27.75, 25.87. LRMS (ESI) *m/z* calculated for C_13_H_21_ClN_3_O_2_^+^ [M + H]^+^: 286.1. Found: 286.

Methyl 11-((6-chloropyrimidin-4-yl)amino)undecanoate (**5**).

11-aminoundecanoic acid (2.00 g, 9.94 mmol) was converted to compound **5** using general procedure A followed by general procedure B. The crude mixture was purified by flash column chromatography on silica gel (0% to 30% ethyl acetate/hexane) to afford **4** (2.50 g, 7.62 mmol, 77% over two steps.) as a white solid. ^1^H NMR (400 MHz, Methanol-*d*_4_) δ 8.20 (s, 1H), 6.47 (s, 1H), 4.91 (s, 1H), 3.64 (s, 3H), 3.42 – 3.10 (m, 2H), 2.30 (t, *J* = 7.4 Hz, 2H), 1.64 – 1.50 (m, 4H), 1.40 – 1.26 (m, 12H); ^13^C NMR (101 MHz, Methanol-*d*_4_) δ 175.82, 164.83, 159.30, 158.22, 105.13, 51.96, 41.83, 34.76, 30.60, 30.50, 30.40, 30.36, 30.17, 27.96, 26.00. LRMS (ESI) *m/z* calculated for C_16_H_27_ClN_3_O_2_^+^ [M + H]^+^: 328.2. Found: 328.

Methyl 6-(N-(6-chloropyrimidin-4-yl)acetamido)hexanoate (**6**).

Compound **3** (2.00 g, 7.76 mmol) was converted to compound **6** using general procedure C. The crude mixture was purified by flash column chromatography on silica gel (0% to 30% ethyl acetate/hexane) to afford **6** (1.97 g, 6.57 mmol, 85%) as a white solid. ^1^H NMR (400 MHz, DMSO-*d*_6_) δ 8.85 (s, 1H), 7.96 (s, 1H), 3.95 – 3.88 (m, 2H), 3.56 (s, 3H), 2.32 (s, 3H), 2.28 (t, *J* = 7.4 Hz, 2H), 1.60 – 1.47 (m, 4H), 1.31 – 1.21 (m, 2H); ^13^C NMR (101 MHz, DMSO-*d*_6_) δ 173.25, 171.84, 161.38, 160.28, 158.11, 113.93, 51.19, 46.05, 33.13, 27.68, 25.65, 24.50, 24.05. LRMS (ESI) *m/z* calculated for C_13_H_18_ClN_3_O_3_Na^+^ [M + Na]^+^: 322.1 Found: 322.

Methyl 8-(N-(6-chloropyrimidin-4-yl)acetamido)octanoate (**7**).

Compound **4** (2.00 g, 6.70 mmol) was converted to compound **7** using general procedure C. The crude mixture was purified by flash column chromatography on silica gel (0% to 30% ethyl acetate/hexane) to afford **7** (1.94 g, 5.92 mmol, 88%) as a white solid. ^1^H NMR (400 MHz, DMSO-*d*_6_) δ 8.86 (s, 1H), 7.97 (s, 1H), 3.95 – 3.89 (m, 2H), 3.56 (s, 3H), 2.32 (s, 3H), 2.27 (t, *J* = 7.4 Hz, 2H), 1.60 – 1.44 (m, 4H), 1.28 – 1.20 (m, 6H); ^13^C NMR (101 MHz, DMSO-*d*_6_) δ 173.36, 171.85, 161.46, 160.26, 158.16, 113.98, 51.19, 46.19, 33.23, 28.37, 28.27, 27.94, 26.02, 24.52, 24.35. LRMS (ESI) *m/z* calculated for C_15_H_22_ClN_3_O_3_Na^+^ [M + Na]^+^: 350.1 Found: 350.

Methyl 11-(N-(6-chloropyrimidin-4-yl)acetamido)undecanoate (**8**).

Compound **5** (2.00 g, 6.10 mmol) was converted to compound **8** using general procedure C. The crude mixture was purified by flash column chromatography on silica gel (0% to 30% ethyl acetate/hexane) to afford **8** (1.85 g, 5.00 mmol, 82%) as a white solid. ^1^H NMR (400 MHz, DMSO-*d*_6_) δ 8.86 (s, 1H), 7.97 (s, 1H), 3.95 – 3.89 (m, 2H), 3.57 (s, 3H), 2.32 (s, 3H), 2.27 (t, *J* = 7.4 Hz, 2H), 1.59 – 1.43 (m, 4H), 1.28 – 1.17 (m, 12H); ^13^C NMR (101 MHz, DMSO-*d*_6_) δ 173.38, 171.83, 161.49, 160.26, 158.16, 114.02, 51.18, 46.20, 33.27, 28.88, 28.79, 28.68, 28.59, 28.46, 27.99, 26.16, 24.51, 24.44. LRMS (ESI) *m/z* calculated for C_18_H_28_ClN_3_O_3_Na^+^ [M + Na]^+^: 392.2 Found: 392.

*tert*-Butyl 3-(2-((6-chloropyrimidin-4-yl)amino)ethoxy)propanoate (**9**).

*tert*-Butyl 3-(2-aminoethoxy)propanoate (2.00 g, 10.6 mmol) was converted to compound **9** using general procedure A. The crude mixture was purified by flash column chromatography on silica gel (10% to 50% ethyl acetate/hexane) to afford **9** (2.36 g, 7.82 mmol, 74%) as a colorless oil. ^1^H NMR (400 MHz, Methanol-*d*_4_) δ 8.22 (s, 1H), 6.50 (s, 1H), 4.88 (s, 1H), 3.68 (t, *J* = 6.1 Hz, 2H), 3.62 – 3.35 (m, 4H), 2.47 (t, *J* = 6.1 Hz, 2H), 1.42 (s, 9H); ^13^C NMR (101 MHz, Methanol-*d*_4_) δ 172.74, 164.84, 159.25, 158.37, 105.29, 81.67, 70.10, 67.60, 41.59, 37.08, 28.33. LRMS (ESI) *m/z* calculated for C_13_H_21_ClN_3_O_3_^+^ [M + H]^+^: 302.1. Found: 302.

*tert*-Butyl 3-(2-(2-((6-chloropyrimidin-4-yl)amino)ethoxy)ethoxy)propanoate (**10**).

*tert*-Butyl 3-(2-(2-aminoethoxy)ethoxy)propanoate (2.00 g, 8.57 mmol) was converted to compound **10** using general procedure A. The crude mixture was purified by flash column chromatography on silica gel (10% to 50% ethyl acetate/hexane) to afford **10** (2.30 g, 6.65 mmol, 78%) as a colorless oil. ^1^H NMR (400 MHz, Methanol-*d*_4_) δ 8.22 (s, 1H), 6.53 (s, 1H), 4.84 (s, 1H), 3.68 (t, *J* = 6.2 Hz, 2H), 3.65 – 3.52 (m, 8H), 2.47 (t, *J* = 6.2 Hz, 2H), 1.43 (s, 9H); ^13^C NMR (101 MHz, Methanol-*d*_4_) δ 172.50, 164.74, 159.24, 158.33, 105.27, 81.54, 71.30, 71.23, 70.30, 67.81, 41.65, 37.05, 28.36. LRMS (ESI) *m/z* calculated for C_15_H_25_ClN_3_O_4_^+^ [M + H]^+^: 346.2. Found: 346.

*tert*-Butyl 3-(2-(2-(2-((6-chloropyrimidin-4-yl)amino)ethoxy)ethoxy)ethoxy)propanoate (**11**).

*tert*-Butyl 3-(2-(2-(2-aminoethoxy)ethoxy)ethoxy)propanoate (2.00 g, 7.21 mmol) was converted to compound **11** using general procedure A. The crude mixture was purified by flash column chromatography on silica gel (10% to 50% ethyl acetate/hexane) to afford **11** (2.20 g, 5.64 mmol, 78%) as a colorless oil. ^1^H NMR (400 MHz, Methanol-*d*_4_) δ 8.23 (s, 1H), 6.53 (s, 1H), 4.83 (s, 1H), 3.68 (t, *J* = 6.2 Hz, 2H), 3.66 – 3.54 (m, 12H), 2.47 (t, *J* = 6.3 Hz, 2H), 1.43 (s, 9H); ^13^C NMR (101 MHz, Methanol-*d*_4_) δ 172.48, 164.78, 159.26, 158.34, 105.27, 81.50, 71.51, 71.41, 71.26, 70.31, 67.79, 41.66, 37.09, 28.37. LRMS (ESI) *m/z* calculated for C_17_H_29_ClN_3_O_5_^+^ [M + H]^+^: 390.2. Found: 390.

*tert*-Butyl 3-(2-(N-(6-chloropyrimidin-4-yl)acetamido)ethoxy)propanoate (**12**).

Compound **9** (2.00 g, 6.63 mmol) was converted to compound **12** using general procedure C. The crude mixture was purified by flash column chromatography on silica gel (10% to 50% ethyl acetate/hexane) to afford **12** (1.85 g, 5.38 mmol, 84%) as a colorless oil. ^1^H NMR (400 MHz, Methanol-*d*_4_) δ 8.74 (s, 1H), 8.07 (s, 1H), 4.24 (t, *J* = 5.2 Hz, 2H), 3.69 (t, *J* = 5.2 Hz, 2H), 3.62 (t, *J* = 6.0 Hz, 2H), 2.42 (s, 3H), 2.40 (t, *J* = 5.9 Hz, 2H), 1.41 (s, 9H); ^13^C NMR (101 MHz, Methanol-*d*_4_) δ 174.77, 172.43, 162.89, 162.10, 158.71, 115.54, 81.57, 70.03, 67.91, 48.07, 37.15, 28.34, 25.37. LRMS (ESI) *m/z* calculated for C_15_H_22_ClN_3_O_4_Na^+^ [M + Na]^+^: 366.1 Found: 366.

*tert-*Butyl 3-(2-(2-(N-(6-chloropyrimidin-4-yl)acetamido)ethoxy)ethoxy)propanoate (**13**).

Compound **10** (2.00 g, 5.78 mmol) was converted to compound **13** using general procedure C. The crude mixture was purified by flash column chromatography on silica gel (10% to 50% ethyl acetate/hexane) to afford **13** (1.91 g, 4.92 mmol, 85%) as a colorless oil. ^1^H NMR (400 MHz, Methanol-*d*_4_) δ 8.75 (s, 1H), 8.07 (s, 1H), 4.24 (t, *J* = 5.3 Hz, 2H), 3.72 (t, *J* = 5.2 Hz, 2H), 3.63 (t, *J* = 6.2 Hz, 2H), 3.57 – 3.48 (m, 4H), 2.43 (s, 3H), 2.44 (t, *J* = 6.4 Hz, 2H), 1.44 (s, 9H); ^13^C NMR (101 MHz, Methanol-*d*_4_) δ 174.76, 172.54, 163.15, 162.15, 158.79, 115.76, 81.57, 71.60, 71.38, 70.27, 67.85, 48.15, 37.12, 28.36, 25.26. LRMS (ESI) *m/z* calculated for C_17_H_26_ClN_3_O_5_Na^+^ [M + Na]^+^: 410.1 Found: 410.

*tert*-Butyl 3-(6-chloropyrimidin-4-yl)-2-oxo-6,9,12-trioxa-3-azapentadecan-15-oate (**14**).

Compound **11** (2.00 g, 5.13 mmol) was converted to compound **14** using general procedure C. The crude mixture was purified by flash column chromatography on silica gel (10% to 50% ethyl acetate/hexane) to afford **14** (1.90 g, 4.40 mmol, 86%) as a colorless oil. ^1^H NMR (400 MHz, Methanol-*d*_4_) δ 8.76 (s, 1H), 8.08 (s, 1H), 4.25 (t, *J* = 5.2 Hz, 2H), 3.73 (t, *J* = 5.2 Hz, 2H), 3.67 (t, *J* = 6.2 Hz, 2H), 3.55 (d, *J* = 0.9 Hz, 8H), 2.46 (t, *J* = 6.2 Hz, 2H), 2.44 (s, 3H), 1.44 (s, 9H); ^13^C NMR (101 MHz, Methanol-*d*_4_) δ 174.79, 172.61, 163.19, 162.15, 158.82, 115.79, 81.59, 71.62, 71.55, 71.47, 71.36, 70.27, 67.83, 48.17, 37.16, 28.36, 25.25. LRMS (ESI) *m/z* calculated for C_19_H_30_ClN_3_O_6_Na^+^ [M + Na]^+^: 454.2 Found: 454.

*tert*-butyl 4-(2-(2,6-dioxopiperidin-3-yl)-1,3-dioxoisoindolin-5-yl)piperazine-1-carboxylate (**16**).

To a solution of **15** (2.00 g, 7.24 mmol) in 15 mL of DMSO, *tert*-butyl piperazine-1-carboxylate (1.35 g , 7.24 mmol) and DIPEA (1.89 mL, 10.7 mmol) were added. The mixture was stirred at 90 ^o^C for 6 h, cooled to room temperature, poured into ice water and extracted with ethyl acetate. The combined organic layers were washed with brine, dried over Na_2_SO_4_, filtered, and concentrated under reduced pressure. The crude mixture was purified by flash column chromatography on silica gel (0% to 50% ethyl acetate/hexane) to afford **16** (2.37 g, 5.36 mmol, 94%) as a yellow solid. ^1^H NMR (400 MHz, DMSO-*d*_6_) δ 11.12 (s, 1H), 7.67 (d, *J* = 8.5 Hz, 1H), 7.32 (d, *J* = 2.3 Hz, 1H), 7.21 (dd, *J* = 8.6, 2.3 Hz, 1H), 5.08 (dd, *J* = 12.9, 5.4 Hz, 1H), 3.51 – 3.40 (m, 8H), 2.97 – 2.82 (m, 1H), 2.65 – 2.47 (m, 2H), 2.06 – 1.98 (m, 1H), 1.42 (s, 9H); ^13^C NMR (101 MHz, DMSO-*d*_6_) δ 172.90, 170.16, 167.57, 167.02, 154.99, 153.89, 133.87, 124.96, 118.59, 117.91, 108.11, 79.20, 54.97, 48.83, 46.62, 31.06, 28.10, 22.24. LRMS (ESI) *m/z* calculated for C_22_H_27_N_4_O_6_^+^ [M + H]^+^: 443.2 Found: 443.

2-(2,6-dioxopiperidin-3-yl)-5-(piperazin-1-yl)isoindoline-1,3-dione hydrochloride (**17**).

To a solution of **16** (2.00 g, 4.52 mmol) in DCM (4 mL) was added HCl (4.0 M in 1,4-dioxane, 12.0 mL) at 0 °C. The mixture was stirred at room temperature for 3 h and concentrated under reduced pressure. The residue was solidified by addition of diethyl ether, filtered and dried to yield **17** (1.61 g, 4.25 mmol, 94%) as a yellow solid. The obtained solid proceeded to the next step without any further purification. LRMS (ESI) *m/z* calculated for C_17_H_19_N_4_O_4_^+^ [M + H]^+^: 343.1 Found: 343.

**General Procedure D for S_N_Ar of Compound 1 with Chloropyrimidine Intermediates.**

To a solution of compound **1** (1.0 equiv) in DMF (0.2 M), corresponding chloropyrimidine (1.0 equiv) and Et_3_N (3.0 equiv) were added. The mixture was stirred at 70 ^o^C for 5 h, cooled to room temperature, saturated aqueous NaHCO_3_ solution was added and extracted with ethyl acetate. The combined organic layers were washed with brine, dried over Na_2_SO_4_, filtered, and concentrated under reduced pressure. The obtained residues were subjected to flash column chromatography on silica gel.

Methyl 6-(N-(6-(4-(4-((4,4-dimethylpiperidin-1-yl)methyl)-2,5-difluorophenyl)-2-oxo-1,4,9-triazaspiro[5.5]undecan-9-yl)pyrimidin-4-yl)acetamido)hexanoate (**18**).

Compound **18** was prepared by S_N_Ar reaction of compound **1** (100 mg, 0.23 mmol) with compound **6** (69 mg, 0.23 mmol) using general procedure D. The crude mixture was purified by flash column chromatography on silica gel (0% to 10% methanol/CH_2_Cl_2_) to afford **18** (115 mg, 0.17 mmol, 75%) as a white solid. ^1^H NMR (400 MHz, DMSO-*d*_6_) δ 8.41 (s, 1H), 8.27 (s, 1H), 7.13 (dd, *J* = 13.0, 6.7 Hz, 1H), 6.92 (dd, *J* = 11.5, 7.3 Hz, 1H), 6.87 (s, 1H), 3.99 (s, 2H), 3.74 (t, *J* = 7.4 Hz, 2H), 3.60 (s, 2H), 3.56 (s, 3H), 3.56 – 3.36 (m, 4H), 3.27 (s, 2H), 2.37 – 2.26 (m, 4H), 2.24 (t, *J* = 7.4 Hz, 2H), 2.06 (s, 3H), 1.88 – 1.77 (m, 2H), 1.75 – 1.65 (m, 2H), 1.53 – 1.39 (m, 4H), 1.32 – 1.25 (m, 4H), 1.25 – 1.17 (m, 2H), 0.85 (s, 6H); ^13^C NMR (101 MHz, DMSO-*d*_6_) δ 173.29, 169.59, 166.65, 162.50, 161.54, 157.89, 157.15 (d, *J* = 239.7 Hz), 150.43 (d, *J* = 239.7 Hz), 138.02 (t, *J* = 9.62 Hz), 118.64 – 118.12 (m), 117.89 – 117.38 (m), 106.44 (d, *J* = 27.3 Hz), 97.93, 55.03, 54.07, 52.86, 52.57, 51.19, 49.12, 45.81, 38.27, 34.76, 33.18, 28.12, 27.57, 25.69, 24.13, 23.38. LRMS (ESI) *m/z* calculated for C_35_H_50_F_2_N_7_O_4_^+^ [M + H]^+^: 670.4 Found: 670.

Methyl 8-(N-(6-(4-(4-((4,4-dimethylpiperidin-1-yl)methyl)-2,5-difluorophenyl)-2-oxo-1,4,9-triazaspiro[5.5]undecan-9-yl)pyrimidin-4-yl)acetamido)octanoate (**19**).

Compound **19** was prepared by S_N_Ar reaction of compound **1** (100 mg, 0.23 mmol) with compound **7** (75 mg, 0.23 mmol) using general procedure D. The crude mixture was purified by flash column chromatography on silica gel (0% to 10% methanol/CH_2_Cl_2_) to afford **19** (113 mg, 0.16 mmol, 70%) as a white solid. ^1^H NMR (400 MHz, DMSO-*d*_6_) δ 8.41 (s, 1H), 8.29 (s, 1H), 7.12 (dd, *J* = 12.9, 6.6 Hz, 1H), 6.92 (dd, *J* = 11.5, 7.3 Hz, 1H), 6.87 (s, 1H), 4.00 (s, 2H), 3.74 (t, *J* = 7.3 Hz, 2H), 3.60 (s, 2H), 3.55 (s, 3H), 3.57 – 3.36 (m, 4H), 3.26 (s, 2H), 2.37 – 2.26 (m, 4H), 2.25 (t, *J* = 7.4 Hz, 2H), 2.06 (s, 3H), 1.87 – 1.78 (m, 2H), 1.75 – 1.65 (m, 2H), 1.52 – 1.37 (m, 4H), 1.32 – 1.25 (m, 4H), 1.24 – 1.16 (m, 6H), 0.85 (s, 6H); ^13^C NMR (101 MHz, DMSO-*d*_6_) δ = 173.33, 169.56, 166.64, 162.50, 161.58, 157.85, 157.13 (d, *J* = 239.8 Hz), 150.03 (d, *J*= 239.7 Hz), 137.96 (t, *J* = 9.5 Hz), 118.59 – 118.20, 117.64 (dd, *J*= 22.5, 5.6 Hz), 106.41 (d, *J* = 27.2 Hz), 97.85, 55.07, 54.06, 52.86, 52.56, 51.16, 49.13, 45.98, 38.27, 34.77, 33.25, 28.43, 28.38, 28.10, 27.86, 26.10, 23.40. LRMS (ESI) *m/z* calculated for C_37_H_54_F_2_N_7_O_4_^+^ [M + H]^+^: 698.4 Found: 698.

Methyl 11-(N-(6-(4-(4-((4,4-dimethylpiperidin-1-yl)methyl)-2,5-difluorophenyl)-2-oxo-1,4,9-triazaspiro[5.5]undecan-9-yl)pyrimidin-4-yl)acetamido)undecanoate (**20**).

Compound **20** was prepared by S_N_Ar reaction of compound **1** (100 mg, 0.23 mmol) with compound **8** (85 mg, 0.23 mmol) using general procedure D. The crude mixture was purified by flash column chromatography on silica gel (0% to 10% methanol/CH_2_Cl_2_) to afford **20** (122 mg, 0.16 mmol, 72%) as a white solid. ^1^H NMR (400 MHz, DMSO-*d*_6_) δ 8.40 (s, 1H), 8.30 (s, 1H), 7.11 (dd, *J* = 13.0, 6.6 Hz, 1H), 6.91 (dd, *J* = 11.5, 7.3 Hz, 1H), 6.86 (s, 1H), 3.99 (s, 2H), 3.74 (t, *J* = 7.4 Hz, 2H), 3.60 (s, 2H), 3.55 (s, 3H), 3.59 – 3.31 (m, 4H), 3.26 (s, 2H), 2.28 (d, *J* = 18.1 Hz, 4H), 2.23 (d, *J* = 7.4 Hz, 2H), 2.06 (s, 3H), 1.87 – 1.77 (m, 2H), 1.75 – 1.64 (m, 2H), 1.52 – 1.36 (m, 4H), 1.33 – 1.24 (m, 4H), 1.24 – 1.11 (m, 12H), 0.84 (s, 6H); ^13^C NMR (101 MHz, DMSO-*d*_6_) δ = 173.30, 169.53, 166.62, 162.50, 161.58, 157.82, 156.76 (d, *J* = 240.8 Hz), 150.43 (d, *J* = 239.7 Hz), 137.97 (t, *J* = 9.67 Hz), 118.68 – 118.17 (m), 117.84 – 117.25 (m), 106.37 (d, *J* = 27.8 Hz), 97.84, 55.12, 54.06, 52.85, 52.54, 51.13, 49.13, 45.99, 38.28, 34.76, 33.28, 28.96, 28.87, 28.74, 28.71, 28.52, 28.09, 27.90, 26.25, 24.47, 23.39. LRMS (ESI) *m/z* calculated for C_40_H_60_F_2_N_7_O_4_^+^ [M + H]^+^: 740.5 Found: 740.

*tert*-Butyl 3-(2-(N-(6-(4-(4-((4,4-dimethylpiperidin-1-yl)methyl)-2,5-difluorophenyl)-2-oxo-1,4,9-triazaspiro[5.5]undecan-9-yl)pyrimidin-4-yl)acetamido)ethoxy)propanoate (**21**).

Compound **21** was prepared by S_N_Ar reaction of compound **1** (100 mg, 0.23 mmol) with compound **9** (79 mg, 0.23 mmol) using general procedure D. The crude mixture was purified by flash column chromatography on silica gel (0% to 10% methanol/CH_2_Cl_2_) to afford **21** (107 mg, 0.15 mmol, 65%) as a white solid. ^1^H NMR (400 MHz, Methanol-*d*_4_) δ 8.42 (s, 1H), 7.18 (dd, *J* = 13.0, 6.7 Hz, 1H), 6.96 (s, 1H), 6.89 (dd, *J* = 11.2, 7.3 Hz, 1H), 4.21 – 4.06 (m, 2H), 3.96 (t, *J* = 5.3 Hz, 2H), 3.73 (s, 2H), 3.65 – 3.56 (m, 6H), 3.55 (s, 2H), 3.41 (s, 2H), 2.54 – 2.44 (m, 4H), 2.41 (t, *J* = 6.0 Hz, 2H), 2.12 (s, 3H), 2.09 – 2.01 (m, 2H), 1.90 – 1.80 (m, 2H), 1.45 – 1.42 (m, 4H), 1.41 (s, 9H), 0.93 (s, 6H); ^13^C NMR (101 MHz, Methanol-*d*_4_) δ 173.04, 172.67, 170.29, 164.45, 162.98, 159.13, 158.99 (d, *J* = 242.5 Hz), 152.39 (d, *J* = 241.5 Hz), 139.85 (t, *J* = 9.90 Hz), 119.84 (dd, *J* = 23.3, 5.9 Hz), 118.67 (dd, *J* = 17.2, 6.5 Hz), 107.11 (d, *J* = 28.4 Hz), 100.80, 81.70, 69.49, 67.73, 56.19, 55.37, 54.55, 53.76, 50.49, 41.61, 39.19, 37.26, 35.98, 29.13, 28.38, 23.39. LRMS (ESI) m/z calculated for C_37_H_54_F_2_N_7_O_5_^+^ [M + H]^+^: 714.4 Found: 714.

*tert*-Butyl 3-(2-(2-(N-(6-(4-(4-((4,4-dimethylpiperidin-1-yl)methyl)-2,5-difluorophenyl)-2-oxo-1,4,9-triazaspiro[5.5]undecan-9-yl)pyrimidin-4-yl)acetamido)ethoxy)ethoxy)propanoate (**22**).

Compound **22** was prepared by S_N_Ar reaction of compound **1** (100 mg, 0.23 mmol) with compound **10** (89 mg, 0.23 mmol) using general procedure D. The crude mixture was purified by flash column chromatography on silica gel (0% to 10% methanol/CH_2_Cl_2_) to afford **22** (118 mg, 0.16 mmol, 68%) as a white solid. ^1^H NMR (400 MHz, Methanol-*d*_4_) δ 8.41 (s, 1H), 7.18 (dd, *J* = 13.0, 6.7 Hz, 1H), 6.98 (s, 1H), 6.89 (dd, *J* = 11.2, 7.3 Hz, 1H), 4.12 (s, 2H), 3.96 (t, *J* = 5.3 Hz, 2H), 3.73 (s, 2H), 3.66 – 3.62 (m, 4H), 3.61 – 3.55 (m, 2H), 3.55 (s, 2H), 3.56 – 3.52 (m, 4H), 3.41 (s, 2H), 2.52 – 2.44 (m, 4H), 2.42 (t, *J* = 6.4 Hz, 2H), 2.13 (s, 3H), 2.09 – 2.01 (m, 2H), 1.90 – 1.79 (m, 2H), 1.45 – 1.40 (m, 4H), 1.41 (s, 9H), 0.92 (s, 6H); ^13^C NMR (101 MHz, Methanol-*d*_4_) δ 173.02, 172.68, 170.26, 164.47, 163.20, 159.07, 158.99 (d, *J* = 241.7 Hz), 152.38 (d, *J* = 242.8 Hz), 139.84 (t, *J* = 9.9 Hz), 119.83 (dd, *J* = 23.2, 6.0 Hz), 118.75 (dd, *J* = 17.9, 7.1 Hz), 107.27 (d, *J* = 30.6 Hz), 100.63, 81.69, 71.37, 71.32, 69.90, 67.83, 56.22, 55.37, 54.52, 53.77, 50.49, 41.56, 39.20, 37.18, 35.98, 29.13, 28.38, 23.40. LRMS (ESI) m/z calculated for C_39_H_58_F_2_N_7_O_6_^+^ [M + H]^+^: 758.4 Found: 758.

*tert*-Butyl 3-(6-(4-(4-((4,4-dimethylpiperidin-1-yl)methyl)-2,5-difluorophenyl)-2-oxo-1,4,9-triazaspiro[5.5]undecan-9-yl)pyrimidin-4-yl)-2-oxo-6,9,12-trioxa-3-azapentadecan-15-oate (**23**).

Compound **23** was prepared by S_N_Ar reaction of compound **1** (100 mg, 0.23 mmol) with compound **11** (99 mg, 0.23 mmol) using general procedure D. The crude mixture was purified by flash column chromatography on silica gel (0% to 10% methanol/CH_2_Cl_2_) to afford **23** (129 mg, 0.16 mmol, 70%) as a white solid. ^1^H NMR (400 MHz, Methanol-*d*_4_) δ 8.42 (s, 1H), 7.18 (dd, *J* = 13.0, 6.7 Hz, 1H), 7.00 (s, 1H), 6.89 (dd, *J* = 11.2, 7.3 Hz, 1H), 4.13 (s, 2H), 3.97 (t, *J* = 5.4 Hz, 2H), 3.73 (s, 2H), 3.68 – 3.63 (m, 4H), 3.62 – 3.57 (m, 2H), 3.56 (s, 4H), 3.57 – 3.52 (m, 6H), 3.42 (s, 2H), 2.51 – 2.43 (m, 4H), 2.42 (t, *J* = 6.2 Hz, 2H), 2.13 (s, 3H), 2.09 – 2.00 (m, 2H), 1.90 – 1.80 (m, 2H), 1.43 (s, 9H), 1.46 – 1.39 (m, 4H), 0.93 (s, 6H); ^13^C NMR (101 MHz, Methanol-*d*_4_) δ 173.05, 172.71, 170.27, 164.48, 163.21, 159.08, 159.00 (d, *J* = 242.0 Hz), 152.39 (d, *J* = 242.5 Hz), 139.85 (t, *J* = 10.1 Hz), 119.84 (dd, *J* = 23.3, 5.9 Hz), 118.78 (dd, *J* = 17.8, 6.9 Hz), 107.27 (d, *J* = 28.0 Hz), 100.68, 81.69, 71.51, 71.42, 71.36, 69.82, 67.88, 56.22, 56.18, 55.38, 54.54, 53.77, 50.49, 41.59, 39.21, 37.18, 35.99, 29.13, 28.37, 23.40. LRMS (ESI) m/z calculated for C_41_H_62_F_2_N_7_O_7_^+^ [M + H]^+^: 802.5 Found: 802.

**General Procedure E for Basic Hydrolysis of the Ester Group and Acetyl Deprotection.**

To a solution of corresponding intermediate (1.0 equiv) in THF (0.2 M), aqueous 4 N LiOH solution (20 equiv) was added. The reaction mixture was stirred at 50 ^o^C for 16 h, cooled to 0 ^o^C and pH was adjusted to 4 by addition of 1 N aqueous HCl solution. The resulting mixture was extracted with isopropyl alcohol/chloroform (1:4). The combined organic layers were dried over Na_2_SO_4_, filtered, and concentrated under reduced pressure. The obtained carboxylic acids proceeded to the next step without further purification.

**General Procedure F for Acidic Hydrolysis of the Ester Group and Acetyl Deprotection.**

To a solution of corresponding intermediate (1.0 equiv) in MeOH (1.0 M), 4 M HCl in 1,4-dioxane solution (20 equiv) was added. The reaction mixture was stirred at 50 ^o^C for 16 h, cooled to 0 ^o^C and pH was adjusted to 4 by addition of 1 N aqueous NaOH solution. The resulting mixture was extracted with isopropyl alcohol/chloroform (1:4). The combined organic layers were dried over Na_2_SO_4_, filtered, and concentrated under reduced pressure. The obtained carboxylic acids proceeded to the next step without further purification.

**General Procedure G for the Synthesis of METTL3 PROTACs.**

To a solution of carboxylic acid (1.0 equiv) and amine (1.0 equiv) in DMF (0.2 M), HATU(1.0 equiv) and DIPEA (3.0 equiv) were added. The reaction mixture was stirred for 3 h at room temperature, diluted with H_2_O and extracted with isopropyl alcohol/chloroform (1:4). The combined organic layers were dried over Na_2_SO_4_, filtered, and concentrated under reduced pressure. The obtained residues were subjected to reverse phase column chromatography using C18 columns or flash column chromatography on silica gel.

(2*S*,4*R*)-1-((*S*)-2-(6-((6-(4-(4-((4,4-Dimethylpiperidin-1-yl)methyl)-2,5-difluorophenyl)-2-oxo-1,4,9-triazaspiro[5.5]undecan-9-yl)pyrimidin-4-yl)amino)hexanamido)-3,3-dimethylbutanoyl)-4-hydroxy-N-(4-(4-methylthiazol-5-yl)benzyl)pyrrolidine-2-carboxamide (**KH11**).

Compound **18** (50 mg, 0.07 mmol) was converted to **KH11** using general procedure E, followed by general procedure G with amine **24**. The product was purified with reverse phase column chromatography using RediSep Gold® C18 Reversed Phase Columns (20-40 *μ*m particle size; 0% to 60% methanol/H_2_O (0.1% formic acid)) to afford **KH11** (26 mg, 0.03 mmol, 34% over two steps.) as a white solid. ^1^H NMR (400 MHz, DMSO-*d*_6_) δ 8.98 (s, 1H), 8.62 (t, *J* = 6.1 Hz, 1H), 8.21 (s, 1H), 7.97 (s, 1H), 7.90 (d, *J* = 9.3 Hz, 1H), 7.40 (q, *J* = 8.2 Hz, 4H), 7.14 (dd, *J* = 13.1, 6.7 Hz, 1H), 6.93 (dd, *J* = 11.5, 7.3 Hz, 1H), 6.64 (t, *J* = 5.6 Hz, 1H), 5.60 (s, 1H), 5.17 (d, *J* = 3.4 Hz, 1H), 4.54 (d, *J* = 9.4 Hz, 1H), 4.48 – 4.39 (m, 2H), 4.35 (s, 1H), 4.25 – 4.17 (m, 1H), 3.93 – 3.81 (m, 2H), 3.66 (s, 2H), 3.59 (s, 2H), 3.41 (s, 2H), 3.40 – 3.27 (m, 2H), 3.25 (s, 2H), 3.20 – 3.10 (m, 2H), 2.44 (s, 3H), 2.35 – 2.29 (m, 4H), 2.29 – 2.21 (m, 1H), 2.18 – 1.85 (m, 3H), 1.81 – 1.73 (m, 2H), 1.69 – 1.60 (m, 2H), 1.55 – 1.41 (m, *J* = 7.1, 6.1 Hz, 4H), 1.33 – 1.26 (m, 6H), 0.93 (s, 9H), 0.86 (s, 6H);^13^C NMR (101 MHz, DMSO-*d*_6_) δ 172.13, 172.03, 169.77, 166.62, 163.29, 161.62, 157.39, 156.80 (d, *J* = 240.5 Hz), 151.53, 150.40 (d, *J* = 240.9 Hz), 147.75, 139.56, 138.09 (t, *J* = 10.1 Hz), 131.23, 129.67, 128.68, 127.45, 118.40 – 118.01 (m), 117.74 (dd, *J* = 24.0, 5.0 Hz), 106.45 (d, *J* = 28.3 Hz), 68.94, 58.75, 56.46, 56.34, 54.83, 54.09, 52.87, 52.72, 49.13, 41.68, 38.26, 38.02, 35.27, 34.91, 34.64, 28.80, 28.15, 26.43, 26.27, 25.55, 16.01. HRMS (ESI) m/z calculated for C_54_H_74_F_2_N_11_O_5_S^+^ [M + H]^+^: 1026.5558 Found: 1026.5558.

(2*S*,4*R*)-1-((*S*)-2-(8-((6-(4-(4-((4,4-Dimethylpiperidin-1-yl)methyl)-2,5-difluorophenyl)-2-oxo-1,4,9-triazaspiro[5.5]undecan-9-yl)pyrimidin-4-yl)amino)octanamido)-3,3-dimethylbutanoyl)-4-hydroxy-N-(4-(4-methylthiazol-5-yl)benzyl)pyrrolidine-2-carboxamide (**KH12**).

Compound **19** (50 mg, 0.07 mmol) was converted to **KH12** using general procedure E, followed by general procedure G with amine **24**. The product was purified with reverse phase column chromatography using RediSep Gold® C18 Reversed Phase Columns (20-40 *μ*m particle size; 0% to 60% methanol/H_2_O (0.1% formic acid)) to afford **KH12** (28 mg, 0.03 mmol, 36% over two steps.) as a white solid. ^1^H NMR (400 MHz, DMSO-*d*_6_) δ 8.98 (s, 1H), 8.56 (t, *J* = 6.1 Hz, 1H), 8.18 (s, 1H), 7.97 (s, 1H), 7.84 (d, *J* = 9.3 Hz, 1H), 7.40 (q, *J* = 8.4 Hz, 4H), 7.14 (dd, *J* = 13.1, 6.7 Hz, 1H), 6.92 (dd, *J* = 11.5, 7.4 Hz, 1H), 6.61 (t, *J* = 5.6 Hz, 1H), 5.60 (s, 1H), 5.13 (d, *J* = 3.6 Hz, 1H), 4.54 (d, *J* = 9.4 Hz, 1H), 4.47 – 4.39 (m, 2H), 4.35 (s, 1H), 4.25 – 4.18 (m, 1H), 3.91 – 3.79 (m, 2H), 3.69 – 3.62 (m, 2H), 3.59 (s, 2H), 3.42 (s, 2H), 3.39 – 3.28 (m, 2H), 3.26 (s, 2H), 3.20 – 3.11 (m, 2H), 2.44 (s, 3H), 2.37 – 2.30 (m, 4H), 2.29 – 2.21 (m, 1H), 2.16 – 1.86 (m, 3H), 1.81 – 1.73 (m, 2H), 1.70 – 1.60 (m, 2H), 1.51 – 1.43 (m, 4H), 1.32 – 1.23 (m, 10H), 0.93 (s, 9H), 0.87 (s, 6H); ^13^C NMR (101 MHz, DMSO-*d*_6_) δ 172.07, 171.92, 169.71, 166.53, 163.28, 161.58, 157.31, 156.73 (d, *J* = 241.4 Hz), 151.41, 150.34 (d, *J* = 240.1 Hz), 147.70, 139.49, 138.01 (t, *J* = 10.1 Hz), 131.15, 129.63, 128.62, 127.41, 118.22 (dd, *J* = 16.9, 7.0 Hz), 117.66 (dd, *J* = 23.1, 6.2 Hz), 106.37 (d, *J* = 27.6 Hz), 68.86, 58.68, 56.34, 56.26, 54.78, 54.03, 52.83, 52.65, 49.07, 41.64, 38.23, 37.94, 35.19, 34.85, 34.59, 29.02, 28.68, 28.56, 28.08, 26.45, 26.37, 25.39, 15.93. HRMS (ESI) m/z calculated for C_56_H_78_F_2_N_11_O_5_S^+^ [M + H]^+^: 1054.5871 Found: 1054.5881

(2*S*,4*S*)-1-((*S*)-2-(8-((6-(4-(4-((4,4-dimethylpiperidin-1-yl)methyl)-2,5-difluorophenyl)-2-oxo-1,4,9-triazaspiro[5.5]undecan-9-yl)pyrimidin-4-yl)amino)octanamido)-3,3-dimethylbutanoyl)-4-hydroxy-N-(4-(4-methylthiazol-5-yl)benzyl)pyrrolidine-2-carboxamide (**KH12-NC**).

Compound **19** (50 mg, 0.07 mmol) was converted to **KH12** using general procedure E, followed by general procedure G with amine **25**. The product was purified with reverse phase column chromatography using RediSep Gold® C18 Reversed Phase Columns (20-40 *μ*m particle size; 0% to 60% methanol/H_2_O (0.1% formic acid)) to afford **KH12** (17 mg, 0.02 mmol, 23% over two steps.) as a white solid. ^1^H NMR (400 MHz, DMSO-*d*_6_) δ 8.98 (s, 1H), 8.64 (t, *J* = 6.1 Hz, 1H), 8.18 (s, 1H), 7.97 (s, 1H), 7.84 (d, *J* = 8.8 Hz, 1H), 7.39 (q, *J* = 8.1 Hz, 4H), 7.21 (dd, *J* = 13.1, 6.7 Hz, 1H), 6.94 (dd, *J* = 11.5, 7.3 Hz, 1H), 6.62 (t, *J* = 5.7 Hz, 1H), 5.60 (s, 1H), 5.44 (s, 1H), 4.48 – 4.40 (m, 2H), 4.36 (dd, *J* = 8.5, 6.1 Hz, 1H), 4.29 – 4.16 (m, 2H), 3.93 (dd, *J* = 10.1, 5.7 Hz, 1H), 3.90 – 3.80 (m, 2H), 3.60 (s, 2H), 3.54 (s, 2H), 3.43 (dd, *J* = 10.0, 5.3 Hz, 1H), 3.27 (s, 3H), 3.19 – 3.11 (m, 2H), 2.44 (s, 3H), 2.47 – 2.41 (m, 4H), 2.36 – 2.27 (m, 1H), 2.27 – 2.19 (m, 1H), 2.15 – 2.05 (m, 1H), 2.00 – 1.90 (m, 1H), 1.81 – 1.72 (m, 3H), 1.71 – 1.60 (m, 2H), 1.53 – 1.41 (m, 4H), 1.37 – 1.20 (m, 10H), 0.94 (s, 9H), 0.88 (s, 6H); ^13^C NMR (101 MHz, DMSO-*d*_6_) δ 172.45, 172.40, 169.97, 166.49, 163.27, 161.58, 157.30, 156.88 (d, *J* = 240.9 Hz), 151.45, 150.23 (d, *J* = 240.5 Hz), 147.73, 139.20, 138.39 (t, *J* = 9.9 Hz), 131.12, 129.72, 128.65, 127.43, 118.04 (dd, *J* = 22.4, 6.2 Hz), 116.82 (dd, *J* = 22.3, 4.6 Hz), 106.35 (d, *J* = 27.5 Hz), 69.08, 58.49, 56.59, 56.50, 55.58, 54.70, 52.77, 52.64, 48.88, 41.77, 37.71, 36.93, 35.77, 34.73, 34.64, 34.57, 28.99, 28.66, 28.54, 28.00, 26.45, 26.37, 25.35, 15.93. HRMS (ESI) m/z calculated for C_56_H_78_F_2_N_11_O_5_S^+^ [M + H]^+^: 1054.5871 Found: 1054.5888.

(2*S*,4*R*)-1-((*S*)-2-(11-((6-(4-(4-((4,4-dimethylpiperidin-1-yl)methyl)-2,5-difluorophenyl)-2-oxo-1,4,9-triazaspiro[5.5]undecan-9-yl)pyrimidin-4-yl)amino)undecanamido)-3,3-dimethylbutanoyl)-4-hydroxy-N-(4-(4-methylthiazol-5-yl)benzyl)pyrrolidine-2-carboxamide (**KH13**).

Compound **20** (50 mg, 0.07 mmol) was converted to **KH13** using general procedure E, followed by general procedure G with amine **24**. The product was purified with reverse phase column chromatography using RediSep Gold® C18 Reversed Phase Columns (20-40 *μ*m particle size; 0% to 60% methanol/H_2_O (0.1% formic acid)) to afford **KH13** (21 mg, 0.02 mmol, 28% over two steps.) as a white solid. ^1^H NMR (400 MHz, DMSO-*d*_6_) δ 8.97 (s, 1H), 8.55 (t, *J* = 6.0 Hz, 1H), 8.18 (s, 1H), 7.97 (s, 1H), 7.83 (d, *J* = 9.3 Hz, 1H), 7.40 (q, *J* = 8.1 Hz, 4H), 7.13 (dd, *J* = 13.1, 6.7 Hz, 1H), 6.92 (dd, *J* = 11.5, 7.3 Hz, 1H), 6.60 (t, *J* = 5.5 Hz, 1H), 5.60 (s, 1H), 5.12 (d, *J* = 3.5 Hz, 1H), 4.54 (d, *J* = 9.3 Hz, 1H), 4.47 – 4.39 (m, 2H), 4.35 (s, 1H), 4.26 – 4.18 (m, 1H), 3.92 – 3.78 (m, 2H), 3.70 – 3.62 (m, 2H), 3.59 (s, 2H), 3.42 (s, 2H), 3.38 – 3.28 (m, 2H), 3.25 (s, 2H), 3.19 – 3.11 (m, 2H), 2.44 (s, 3H), 2.38 – 2.30 (m, 4H), 2.28 – 2.20 (m, 1H), 2.14 – 1.86 (m, 3H), 1.81 – 1.73 (m, 2H), 1.71 – 1.60 (m, 2H), 1.53 – 1.42 (m, 4H), 1.33 – 1.18 (m, 16H), 0.93 (s, 9H), 0.86 (s, 6H); ^13^C NMR (101 MHz, DMSO-*d*_6_) δ 172.09, 171.94, 169.72, 166.54, 163.29, 161.60, 157.32, 156.76 (d, *J* = 240.3 Hz), 151.41, 150.35 (d, *J* = 240.4 Hz), 147.70, 139.50, 138.03 (t, *J* = 9.6 Hz), 131.16, 129.64, 128.63, 127.42, 118.45 – 118.02 (m), 117.62 (dd, *J* = 22.9, 5.5 Hz), 106.38 (d, *J* = 28.7 Hz), 68.87, 58.69, 56.34, 56.27, 54.80, 54.02, 52.83, 52.65, 49.07, 41.65, 38.22, 37.95, 35.21, 34.88, 34.59, 29.04, 28.95, 28.85, 28.77, 28.67, 28.08, 26.54, 26.37, 25.44, 15.94. HRMS (ESI) m/z calculated for C_59_H_84_F_2_N_11_O_5_S^+^ [M + H]^+^: 1096.6340 Found: 1096.6361.

(2*S*,4*R*)-1-((*S*)-2-(3-(2-((6-(4-(4-((4,4-dimethylpiperidin-1-yl)methyl)-2,5-difluorophenyl)-2-oxo-1,4,9-triazaspiro[5.5]undecan-9-yl)pyrimidin-4-yl)amino)ethoxy)propanamido)-3,3-dimethylbutanoyl)-4-hydroxy-N-(4-(4-methylthiazol-5-yl)benzyl)pyrrolidine-2-carboxamide (**KH14**).

Compound **21** (50 mg, 0.07 mmol) was converted to **KH14** using general procedure F, followed by general procedure G with amine **24**. The product was purified with reverse phase column chromatography using RediSep Gold® C18 Reversed Phase Columns (20-40 *μ*m particle size; 0% to 60% methanol/H_2_O (0.1% formic acid)) to afford **KH14** (13 mg, 0.01 mmol, 18% over two steps.) as a white solid. ^1^H NMR (400 MHz, DMSO-*d*_6_) δ 8.98 (s, 1H), 8.62 (t, *J* = 6.1 Hz, 1H), 8.20 (s, 1H), 8.02 – 7.94 (m, 2H), 7.39 (q, *J* = 8.2 Hz, 4H), 7.13 (dd, *J* = 13.1, 6.7 Hz, 1H), 6.93 (dd, *J* = 11.5, 7.4 Hz, 1H), 6.64 (s, 1H), 5.67 (s, 1H), 5.19 (d, *J* = 3.5 Hz, 1H), 4.56 (d, *J* = 9.4 Hz, 1H), 4.47 – 4.38 (m, 2H), 4.35 (s, 1H), 4.25 – 4.17 (m, 1H), 3.91 – 3.77 (m, 2H), 3.70 – 3.63 (m, 2H), 3.63 – 3.54 (m, 2H), 3.59 (s, 2H), 3.51 – 3.43 (m, 2H), 3.40 (s, 2H), 3.39 – 3.28 (m, 4H), 3.25 (s, 2H), 2.43 (s, 3H), 2.42 – 2.35 (m, 1H), 2.35 – 2.28 (m, 4H), 2.08 – 1.85 (m, 3H), 1.81 – 1.71 (m, 2H), 1.69 – 1.58 (m, 2H), 1.29 (t, *J* = 5.2 Hz, 4H), 0.93 (s, 9H), 0.86 (s, 6H); ^13^C NMR (101 MHz, DMSO-*d*_6_) δ 171.86, 169.99, 169.58, 166.53, 163.17, 162.30, 161.54, 157.52 (d, *J* = 149.4 Hz), 157.31, 151.42, 150.34 (d, *J* = 237.2 Hz), 147.70, 139.47, 138.00 (t, *J* = 6.3 Hz), 131.14, 129.63, 128.62, 127.41, 118.47 – 118.17 (m), 117.85 – 117.43 (m), 106.39 (d, *J* = 28.6 Hz), 68.95, 68.86, 66.71, 58.72, 56.42, 56.32, 54.79, 54.06, 52.84, 52.64, 49.08, 41.64, 38.26, 37.94, 35.62, 35.37, 35.10, 34.58, 28.09, 26.33, 15.93. HRMS (ESI) m/z calculated for C_53_H_72_F_2_N_11_O_6_S^+^ [M + H]^+^: 1028.5350 Found: 1028.5362.

(2*S*,4*R*)-1-((*S*)-2-(3-(2-(2-((6-(4-(4-((4,4-dimethylpiperidin-1-yl)methyl)-2,5-difluorophenyl)-2-oxo-1,4,9-triazaspiro[5.5]undecan-9-yl)pyrimidin-4-yl)amino)ethoxy)ethoxy)propanamido)-3,3-dimethylbutanoyl)-4-hydroxy-N-(4-(4-methylthiazol-5-yl)benzyl)pyrrolidine-2-carboxamide (**KH15**).

Compound **22** (50 mg, 0.07 mmol) was converted to **KH15** using general procedure F, followed by general procedure G with amine **24**. The product was purified with reverse phase column chromatography using RediSep Gold® C18 Reversed Phase Columns (20-40 *μ*m particle size; 0% to 60% methanol/H_2_O (0.1% formic acid)) to afford **KH15** (28 mg, 0.03 mmol, 40% over two steps.) as a white solid. ^1^H NMR (400 MHz, DMSO-*d*_6_) δ 8.98 (s, 1H), 8.61 (t, *J* = 6.1 Hz, 1H), 8.20 (s, 1H), 8.00 – 7.94 (m, 2H), 7.40 (q, *J* = 8.3 Hz, 4H), 7.14 (dd, *J* = 13.1, 6.8 Hz, 1H), 6.93 (dd, *J* = 11.5, 7.3 Hz, 1H), 6.64 (s, 1H), 5.68 (s, 1H), 5.16 (d, *J* = 2.2 Hz, 1H), 4.56 (d, *J* = 9.4 Hz, 1H), 4.47 – 4.39 (m, 2H), 4.35 (s, 1H), 4.25 – 4.17 (m, 1H), 3.91 – 3.78 (m, 2H), 3.68 – 3.62 (m, 2H), 3.62 – 3.55 (m, 2H), 3.59 (s, 2H), 3.54 – 3.45 (m, 6H), 3.42 (s, 2H), 3.40 – 3.27 (m, 4H), 3.25 (s, 2H), 2.44 (s, 3H), 2.40 – 2.34 (m, 1H), 2.35 – 2.30 (m, 4H), 2.08 – 1.85 (m, 3H), 1.83 – 1.71 (m, 2H), 1.70 – 1.58 (m, 2H), 1.29 (t, *J* = 5.5 Hz, 4H), 0.93 (s, 9H), 0.86 (s, 6H); ^13^C NMR (101 MHz, DMSO-*d*_6_) δ 171.99, 170.01, 169.57, 166.60, 163.24, 162.36, 161.58, 157.39, 156.80 (d, *J* = 241.1 Hz), 151.53, 150.39 (d, *J* = 241.0 Hz), 147.75, 139.54, 138.12 (t, *J* = 9.9 Hz), 131.21, 129.67, 128.68, 127.45, 118.43 – 117.90 (m), 117.71 (dd, *J* = 23.6, 6.5 Hz), 106.59 (d, *J* = 28.6 Hz), 69.62, 69.50, 69.30, 68.92, 66.99, 58.75, 56.46, 56.32, 54.81, 54.06, 52.86, 52.71, 49.11, 41.67, 38.22, 38.01, 35.84, 35.66, 35.44, 34.61, 28.15, 26.36, 16.00. HRMS (ESI) m/z calculated for C_55_H_76_F_2_N_11_O_7_S^+^ [M + H]^+^: 1072.5613 Found: 1072.5633.

(2*S*,4*R*)-1-((*S*)-14-(tert-butyl)-1-((6-(4-(4-((4,4-dimethylpiperidin-1-yl)methyl)-2,5-difluorophenyl)-2-oxo-1,4,9-triazaspiro[5.5]undecan-9-yl)pyrimidin-4-yl)amino)-12-oxo-3,6,9-trioxa-13-azapentadecan-15-oyl)-4-hydroxy-N-(4-(4-methylthiazol-5-yl)benzyl)pyrrolidine-2-carboxamide (**KH16**).

Compound **23** (50 mg, 0.06 mmol) was converted to **KH16** using general procedure F, followed by general procedure G with amine **24**. The product was purified with reverse phase column chromatography using RediSep Gold® C18 Reversed Phase Columns (20-40 *μ*m particle size; 0% to 60% methanol/H_2_O (0.1% formic acid)) to afford **KH16** (23 mg, 0.02 mmol, 33% over two steps.) as a white solid. ^1^H NMR (400 MHz, DMSO-*d*_6_) δ 8.99 (s, 1H), 8.61 (t, *J* = 6.1 Hz, 1H), 8.21 (s, 1H), 8.01 – 7.94 (m, 2H), 7.40 (q, *J* = 8.3 Hz, 4H), 7.15 (dd, *J* = 13.1, 6.7 Hz, 1H), 6.94 (dd, *J* = 11.6, 7.4 Hz, 1H), 6.66 (s, 1H), 5.69 (s, 1H), 5.17 (d, *J* = 3.4 Hz, 1H), 4.56 (d, *J* = 9.4 Hz, 1H), 4.48 – 4.40 (m, 2H), 4.35 (s, 1H), 4.26 – 4.18 (m, 1H), 3.92 – 3.77 (m, 2H), 3.71 – 3.63 (m, 2H), 3.59 (s, 2H), 3.62 – 3.55 (m, 2H), 3.53 – 3.45 (m, 10H), 3.42 (s, 2H), 3.40 – 3.28 (m, 4H), 3.26 (s, 2H), 2.44 (s, 3H), 2.40 – 2.35 (m, 1H), 2.35 – 2.29 (m, 4H), 2.08 – 1.85 (m, 3H), 1.82 – 1.72 (m, 2H), 1.70 – 1.60 (m, 2H), 1.30 (t, *J* = 5.5 Hz, 4H), 0.93 (s, 9H), 0.87 (s, 6H); ^13^C NMR (101 MHz, DMSO-*d*_6_) δ 171.91, 169.94, 169.53, 166.53, 163.22, 162.30, 161.56, 157.32, 156.74 (d, *J* = 240.7 Hz), 151.48 (d, *J* = 11.0 Hz), 150.34 (d, *J* = 240.5 Hz), 147.71, 139.49, 138.02 (d, *J* = 9.2 Hz), 131.15, 129.64, 128.63, 127.42, 118.43 – 118.00 (m), 117.68 (dd, *J* = 24.4, 4.4 Hz), 106.38 (d, *J* = 28.3 Hz), 69.77, 69.72, 69.63, 69.48, 69.26, 68.86, 66.94, 58.71, 56.36, 56.29, 54.79, 54.03, 52.83, 52.65, 49.07, 41.65, 38.23, 37.94, 35.77, 35.65, 35.35, 34.58, 28.09, 26.32, 15.93. HRMS (ESI) m/z calculated for C_57_H_80_F_2_N_11_O_8_S^+^ [M + H]^+^: 1116.5875 Found: 1116.5892.

5-(4-(6-((6-(4-(4-((4,4-dimethylpiperidin-1-yl)methyl)-2,5-difluorophenyl)-2-oxo-1,4,9-triazaspiro[5.5]undecan-9-yl)pyrimidin-4-yl)amino)hexanoyl)piperazin-1-yl)-2-(2,6-dioxopiperidin-3-yl)isoindoline-1,3-dione (**KH21**).

Compound **18** (50 mg, 0.07 mmol) was converted to **KH21** using general procedure E, followed by general procedure G with amine **17**. The product was purified by flash column chromatography on silica gel (5% to 10% methanol/CH_2_Cl_2_) to afford **KH21** (17 mg, 0.02 mmol, 24% over two steps.) as a yellow solid. ^1^H NMR (400 MHz, DMSO-*d*_6_) δ 11.08 (s, 1H), 8.19 (s, 1H), 7.97 (s, 1H), 7.69 (d, *J* = 8.5 Hz, 1H), 7.33 (d, *J* = 2.3 Hz, 1H), 7.35 – 7.20 (m, 1H), 7.23 (dd, *J* = 8.7, 2.4 Hz, 1H), 6.95 (dd, *J* = 11.6, 7.3 Hz, 1H), 6.65 (t, *J* = 5.5 Hz, 1H), 5.61 (s, 1H), 5.07 (dd, *J* = 12.9, 5.4 Hz, 1H), 3.91 – 3.80 (m, 2H), 3.65 – 3.57 (m, 8H), 3.53 – 3.48 (m, 2H), 3.47 – 3.42 (m, 2H), 3.41 – 3.29 (m, 2H), 3.27 (s, 2H), 3.21 – 3.14 (m, 2H), 2.94 – 2.83 (m, 1H), 2.63 – 2.46 (m, 6H), 2.35 (t, *J* = 7.4 Hz, 2H), 2.05 – 1.97 (m, 1H), 1.80 – 1.71 (m, 2H), 1.70 – 1.62 (m, 2H), 1.57 – 1.47 (m, 4H), 1.35 (d, *J* = 15.8 Hz, 6H), 0.89 (s, 6H); ^13^C NMR (101 MHz, DMSO-*d*_6_) δ 172.89, 170.89, 170.15, 167.58, 167.02, 166.52, 163.29, 161.59, 157.37, 157.05 (d, *J* = 241.5 Hz), 154.90, 150.18 (d, *J* = 240.3 Hz), 138.99 – 138.45 (m), 133.88, 124.99, 118.47, 117.80, 107.97, 106.38 (d, *J* = 26.2 Hz), 54.66, 53.25, 52.75, 52.70, 48.80, 48.75, 46.85, 46.58, 44.13, 40.40, 37.29, 34.59, 32.25, 31.02, 28.97, 28.00, 26.30, 24.53, 22.21. LRMS (ESI) m/z calculated for C_49_H_62_F_2_N_11_O_6_^+^ [M + H]^+^: 938.5 Found: 938.

5-(4-(8-((6-(4-(4-((4,4-dimethylpiperidin-1-yl)methyl)-2,5-difluorophenyl)-2-oxo-1,4,9-triazaspiro[5.5]undecan-9-yl)pyrimidin-4-yl)amino)octanoyl)piperazin-1-yl)-2-(2,6-dioxopiperidin-3-yl)isoindoline-1,3-dione (**KH22**).

Compound **19** (50 mg, 0.07 mmol) was converted to **KH22** using general procedure E, followed by general procedure G with amine **17**. The product was purified by flash column chromatography on silica gel (5% to 10% methanol/CH_2_Cl_2_) to afford **KH22** (16 mg, 0.02 mmol, 23% over two steps.) as a yellow solid. ^1^H NMR (400 MHz, DMSO-*d*_6_) δ 11.08 (s, 1H), 8.19 (s, 1H), 7.97 (s, 1H), 7.69 (d, *J* = 8.5 Hz, 1H), 7.33 (d, *J* = 2.3 Hz, 1H), 7.35 – 7.19 (m, 1H), 7.23 (dd, *J* = 8.6, 2.3 Hz, 1H), 6.94 (dd, *J* = 11.6, 7.3 Hz, 1H), 6.64 (t, *J* = 5.7 Hz, 1H), 5.61 (s, 1H), 5.07 (dd, *J* = 12.9, 5.4 Hz, 1H), 3.90 – 3.80 (m, 2H), 3.64 – 3.56 (m, 8H), 3.53 – 3.48 (m, 2H), 3.46 – 3.41 (m, 2H), 3.41 – 3.30 (m, 2H), 3.27 (s, 2H), 3.19 – 3.13 (m, 2H), 2.93 – 2.83 (m, 1H), 2.62 – 2.52 (m, 2H), 2.50 – 2.36 (m, 4H), 2.34 (t, *J* = 7.4 Hz, 2H), 2.05 – 1.97 (m, 1H), 1.82 – 1.73 (m, 2H), 1.69 – 1.61 (m, 2H), 1.54 – 1.44 (m, 4H), 1.40 – 1.29 (m, 10H), 0.88 (s, 6H); ^13^C NMR (101 MHz, DMSO-*d*_6_) δ 172.77, 170.88, 170.04, 167.50, 166.95, 166.46, 163.27, 161.57, 157.28, 156.97 (d, *J* = 251.1 Hz), 154.85, 150.17 (d, *J* = 244.3 Hz), 133.84, 124.91, 118.45, 117.74, 107.89, 106.31 (d, *J* = 28.4 Hz), 54.67, 53.26, 52.75, 52.64, 48.82, 48.78, 46.81, 46.54, 44.07, 40.34, 37.78, 34.56, 32.21, 30.97, 28.98, 28.78, 28.67, 27.96, 26.42, 24.64, 22.17. LRMS (ESI) m/z calculated for C_51_H_66_F_2_N_11_O_6_^+^ [M + H]^+^: 966.5 Found: 967.

5-(4-(11-((6-(4-(4-((4,4-dimethylpiperidin-1-yl)methyl)-2,5-difluorophenyl)-2-oxo-1,4,9-triazaspiro[5.5]undecan-9-yl)pyrimidin-4-yl)amino)undecanoyl)piperazin-1-yl)-2-(2,6-dioxopiperidin-3-yl)isoindoline-1,3-dione (**KH23**).

Compound **20** (50 mg, 0.07 mmol) was converted to **KH23** using general procedure E, followed by general procedure G with amine **17**. The product was purified by flash column chromatography on silica gel (5% to 10% methanol/CH_2_Cl_2_) to afford **KH23** (18 mg, 0.02 mmol, 26% over two steps.) as a yellow solid. ^1^H NMR (400 MHz, DMSO-*d*_6_) δ 11.08 (s, 1H), 8.19 (s, 1H), 7.97 (s, 1H), 7.70 (d, *J* = 8.5 Hz, 1H), 7.34 (d, *J* = 2.3 Hz, 1H), 7.27 – 7.17 (m, 2H), 6.95 (dd, *J* = 11.6, 7.3 Hz, 1H), 6.62 (t, *J* = 5.7 Hz, 1H), 5.60 (s, 1H), 5.07 (dd, *J* = 12.9, 5.4 Hz, 1H), 3.90 – 3.79 (m, 2H), 3.64 – 3.55 (m, 8H), 3.53 – 3.49 (m, 2H), 3.47 – 3.43 (m, 2H), 3.42 – 3.30 (m, 2H), 3.27 (s, 2H), 3.19 – 3.11 (m, 2H), 2.94 – 2.82 (m, 1H), 2.62 – 2.53 (m, 2H), 2.49 – 2.40 (m, 4H), 2.33 (t, *J* = 7.4 Hz, 2H), 2.07 – 1.96 (m, 1H), 1.81 – 1.71 (m, 2H), 1.70 – 1.60 (m, 2H), 1.53 – 1.43 (m, 4H), 1.37 – 1.32 (m, 4H), 1.29 – 1.23 (m, 12H), 0.88 (s, 6H); ^13^C NMR (101 MHz, DMSO-*d*_6_) δ 172.88, 170.92, 170.14, 167.57, 167.02, 166.55, 163.29, 161.60, 157.36, 156.54 (d, *J* = 238.9 Hz), 154.91, 150.25 (d, *J* = 239.4 Hz), 133.88, 124.98, 118.48, 117.79, 107.98, 106.42 (d, *J* = 30.6 Hz), 54.73, 53.60, 52.69, 48.92, 48.80, 46.85, 46.58, 44.11, 40.37, 37.70, 34.60, 32.26, 31.02, 29.08, 29.00, 28.97, 28.88, 28.85, 28.05, 26.56, 24.75, 22.21. LRMS (ESI) m/z calculated for C_54_H_72_F_2_N_11_O_6_^+^ [M + H]^+^: 1008.6 Found: 1009.

5-(4-(3-(2-((6-(4-(4-((4,4-dimethylpiperidin-1-yl)methyl)-2,5-difluorophenyl)-2-oxo-1,4,9-triazaspiro[5.5]undecan-9-yl)pyrimidin-4-yl)amino)ethoxy)propanoyl)piperazin-1-yl)-2-(2,6-dioxopiperidin-3-yl)isoindoline-1,3-dione (**KH24**).

Compound **21** (50 mg, 0.07 mmol) was converted to **KH24** using general procedure F, followed by general procedure G with amine **17**. The product was purified by flash column chromatography on silica gel (5% to 15% methanol/CH_2_Cl_2_) to afford **KH24** (14 mg, 0.01 mmol, 21% over two steps.) as a yellow solid. ^1^H NMR (400 MHz, DMSO-*d*_6_) δ 11.08 (s, 1H), 8.19 (s, 1H), 7.98 (s, 1H), 7.69 (d, *J* = 8.5 Hz, 1H), 7.33 (d, *J* = 2.3 Hz, 1H), 7.38 – 7.15 (m, 1H), 7.22 (dd, *J* = 8.6, 2.3 Hz, 1H), 7.02 – 6.90 (m, 1H), 6.65 – 6.57 (m, 1H), 5.68 (s, 1H), 5.07 (dd, *J* = 12.9, 5.4 Hz, 1H), 3.84 (d, *J* = 13.3 Hz, 2H), 3.71 – 3.58 (m, 10H), 3.53 – 3.44 (m, 8H), 3.40 – 3.28 (m, 2H), 3.27 (s, 2H), 2.93 – 2.83 (m, 1H), 2.63 (t, *J* = 6.7 Hz, 2H), 2.62 – 2.32 (m, 6H), 2.06 – 1.93 (m, 1H), 1.80 – 1.72 (m, 2H), 1.69 – 1.59 (m, 2H), 1.41 – 1.31 (m, 4H), 0.90 (s, 6H); ^13^C NMR (101 MHz, DMSO-*d*_6_) δ 172.77, 170.04, 169.12, 167.50, 166.95, 166.43, 163.21, 161.54, 157.30, 154.81, 133.84, 124.90, 118.43, 117.69, 107.86, 106.34 (d, *J* = 27.8 Hz), 69.05, 66.50, 54.60, 52.72, 52.62, 48.83, 48.77, 46.75, 46.45, 44.15, 40.38, 35.77, 34.52, 32.83, 30.96, 27.94, 22.16. LRMS (ESI) m/z calculated for C_48_H_60_F_2_N_11_O_7_^+^ [M + H]^+^: 940.5 Found: 940.

5-(4-(3-(2-(2-((6-(4-(4-((4,4-dimethylpiperidin-1-yl)methyl)-2,5-difluorophenyl)-2-oxo-1,4,9-triazaspiro[5.5]undecan-9-yl)pyrimidin-4-yl)amino)ethoxy)ethoxy)propanoyl)piperazin-1-yl)-2-(2,6-dioxopiperidin-3-yl)isoindoline-1,3-dione (**KH25**).

Compound **22** (50 mg, 0.07 mmol) was converted to **KH25** using general procedure F, followed by general procedure G with amine **17**. The product was purified by flash column chromatography on silica gel (5% to 15% methanol/CH_2_Cl_2_) to afford **KH25** (21 mg, 0.02 mmol, 32% over two steps.) as a yellow solid. ^1^H NMR (400 MHz, DMSO-*d*_6_) δ 11.08 (s, 1H), 8.17 (s, 1H), 7.99 (s, 1H), 7.68 (d, *J* = 8.5 Hz, 1H), 7.33 (d, *J* = 2.3 Hz, 1H), 7.22 (dd, *J* = 8.7, 2.4 Hz, 1H), 7.15 (dd, *J* = 13.0, 6.9 Hz, 1H), 6.93 (dd, *J* = 11.5, 7.4 Hz, 1H), 6.59 (s, 1H), 5.69 (s, 1H), 5.07 (dd, *J* = 12.9, 5.4 Hz, 1H), 3.85 (d, *J* = 13.1 Hz, 2H), 3.69 – 3.56 (m, 10H), 3.54 – 3.41 (m, 12H), 3.37 – 3.28 (m, 2H), 3.26 (s, 2H), 2.95 – 2.81 (m, 1H), 2.62 (t, *J* = 6.6 Hz, 2H), 2.60 – 2.52 (m, 2H), 2.42 – 2.28 (m, 4H), 2.06 – 1.96 (m, 1H), 1.80 – 1.72 (m, 2H), 1.68 – 1.58 (m, 2H), 1.30 (t, *J* = 5.5 Hz, 4H), 0.87 (s, 6H); ^13^C NMR (101 MHz, DMSO-*d*_6_) δ 172.77, 170.04, 169.09, 167.50, 166.95, 166.51, 163.22, 161.56, 157.31, 156.77 (d, *J* = 238.5 Hz), 154.82, 150.30 (d, *J* = 242.0 Hz), 133.84, 124.90, 118.44, 117.70, 107.88, 106.36 (d, *J* = 28.5 Hz), 69.67, 69.60, 69.26, 66.73, 54.77, 53.96, 52.81, 52.64, 49.03, 48.77, 46.76, 46.47, 44.16, 40.36, 38.13, 35.77, 34.56, 32.81, 30.97, 28.06, 22.16. LRMS (ESI) m/z calculated for C_50_H_64_F_2_N_11_O_8_^+^ [M + H]^+^: 984.5 Found: 984.

5-(4-(3-(2-(2-(2-((6-(4-(4-((4,4-dimethylpiperidin-1-yl)methyl)-2,5-difluorophenyl)-2-oxo-1,4,9-triazaspiro[5.5]undecan-9-yl)pyrimidin-4-yl)amino)ethoxy)ethoxy)ethoxy)propanoyl)piperazin-1-yl)-2-(2,6-dioxopiperidin-3-yl)isoindoline-1,3-dione (**KH26**)

Compound **23** (50 mg, 0.06 mmol) was converted to **KH26** using general procedure F, followed by general procedure G with amine **17**. The product was purified by flash column chromatography on silica gel (5% to 15% methanol/CH_2_Cl_2_) to afford **KH26** (23 mg, 0.02 mmol, 36% over two steps.) as a yellow solid. ^1^H NMR (400 MHz, DMSO-*d*_6_) δ 11.08 (s, 1H), 8.19 (s, 1H), 7.99 (s, 1H), 7.69 (d, *J* = 8.5 Hz, 1H), 7.34 (d, *J* = 2.3 Hz, 1H), 7.37 – 7.16 (m, 1H), 7.23 (dd, *J* = 8.6, 2.4 Hz, 1H), 7.00 – 6.91 (m, 1H), 6.60 (s, 1H), 5.68 (s, 1H), 5.07 (dd, *J* = 12.9, 5.4 Hz, 1H), 3.91 – 3.78 (m, 2H), 3.67 – 3.56 (m, 10H), 3.55 – 3.40 (m, 16H), 3.39 – 3.29 (m, 2H), 3.27 (s, 2H), 2.97 – 2.81 (m, 1H), 2.62 (t, *J* = 6.6 Hz, 2H), 2.60 – 2.53 (m, 2H), 2.50 – 2.20 (m, 4H), 2.04 – 1.93 (m, 1H), 1.81 – 1.71 (m, 2H), 1.70 – 1.57 (m, 2H), 1.46 – 1.29 (m, 4H), 0.89 (s, 6H); ^13^C NMR (101 MHz, DMSO-*d*_6_) δ 172.88, 170.14, 169.13, 167.57, 167.01, 166.52, 163.23, 161.54, 157.36, 154.88, 133.88, 124.97, 118.46, 117.76, 107.95, 69.80, 69.73, 69.68, 69.29, 66.79, 56.06, 54.74, 52.80, 52.69, 48.91, 48.79, 46.80, 46.50, 44.21, 40.15, 35.83, 34.57, 32.82, 31.01, 28.01, 22.20. LRMS (ESI) m/z calculated for C_52_H_68_F_2_N_11_O_9_^+^ [M + H]^+^: 1028.5 Found: 1029.

**Biology and Biochemistry Methods**

**Cell Culture**

MOLM-14, HL-60, H1299, HCT116, MDA-MB-231, AGS and HeLa cells were obtained from ATCC or KCLB, and MOLM-13 cells were generously provided by Dr. Yong-Chul Kim (GIST, Gwangju, Korea). MOLM-13, MOLM-14, HL-60, H1299, HCT116, MDA-MB-231 and AGS cells were cultured at RPMI-1640. HeLa cells were cultured at DMEM. All media were supplemented with 10% FBS (fetal bovine serum) and 1% P/S (penicillin/streptomycin). Cells were incubated at 37 °C with 5% CO_2._ DPBS, RPMI-1640, DMEM, FBS and P/S were purchased from WELGENE. Cell culture dishes, flasks and multi-well plates were purchased from SPL Life Sciences.

Human gastric cancer cell lines Hs746T, SNU484, SNU668, MKN1, and SNU216 were grown in RPMI supplemented with 10% fetal bovine serum, 100X sodium pyruvate, and 100X antibiotics. and gastric cancer cell line YCC7 was grown in DMEM supplemented with 10% fetal bovine serum with 100X antibiotics.

**Western Blot**

Indicated concentration of compounds and vehicle (DMSO) were treated for indicated times. Cells were harvested, washed once with cold DPBS, and resuspended in cell lysis buffer (50 mM Tris pH 7.4, 2 mM EDTA pH 8, 1% NP40, 150 mM NaCl) supplemented with protease inhibitor cocktail (Roche, 11873580001) and phosphatase inhibitor cocktail (Roche, 06906837001). Protein concentration was assessed by Bradford assay (Bio-Rad, BR5000006) and equal amounts of protein were loaded on 10% SDS-PAGE gel. Separated proteins were transferred to nitrocellulose membranes and blocked with 5% skim milk in TBS/T buffer. After blocking, membranes were incubated overnight at 4 °C with the following primary antibodies: METTL3 (Abcam, ab195352), METTL14 (Sigma, HPA038002), c-MYC (CST, #5605), GAPDH (CST, #5174). All primary antibodies were diluted at 1:1000 in TBS/T. HRP-conjugated secondary antibodies (Gendepot, SA002-500) were diluted at 1:10,000 in TBS/T and incubated with membranes for 1 h. ECL solution (Geneplex, G-E3-0250-1) was dropped on the membrane and the signal was detected by ImageQuant^TM^ LAS 4000.

**CETSA**

MOLM-13 cells were seeded in 100 μl of RPMI-1640 at 2 × 10^7^ cells/ml in a microcentrifuge tube (Axygen, MCT-175-C) and added to a compound dilution series with a final DMSO concentration of 0.5% (v/v). Cells were incubated at 37 °C for 1 h and resuspended in 100 μl of DPBS. Samples were then heated up to 54 °C for 10 m and lysed by three freeze-thaw cycles. METTL3 and GAPDH protein in the supernatant was detected by Western Blot as previously described.

**HCS imaging**

For VHL target engagement assay, 1 × 10^4^ BRD4-eGFP_mCherry expressing HeLa cells/well were seeded on 96 well black plates (PerkinElmer, 6055300). Compounds and vehicle (DMSO) were treated for 4 h after cell adhesion. The plate was scanned by PerkinElmer Operetta CLS^TM.^

**Cell Viability Assay**

For MOLM-13 and AGS, Cell viability assay was performed in duplicate and at least twice. 3000 cells/well in 100 μl of cell culture media were seeded on 96 well plates (Nest, 701003) and serial dilution compounds (maximum 50 μM, 1/3 serial diluted in DMSO, 10 points) were treated with a final DMSO concentration of 0.5% (v/v). Suspension cells were treated after cell seeding and adherent cells were treated after 24 h of cell seeding. After 72 h of treatment, cell viability was determined using CellTiter-Glo luminescent cell viability assay (Promega, G7572) according to manufacturer’s instructions. Luminescence was measured by PerkinElmer EnVision^®^ 2105 and normalized to DMSO. Cell viability curves and IC_50_ values were fitted with nonlinear regression in GraphPad Prism 8.0.2 software.

For Hs746T, SNU484, SNU668, MKN1 and SNU216, cells were seeded in a 96-well white microplate at 20,000 cells per well in fully supplemented media and incubated with compounds (final DMSO concentration at 0.1%). Relative cell viability was measured 72h after the addition of drug using CellTiter-Glo (Promega) according to the manufacturer’s protocol. Each analysis was performed in two independent replicates with triplicate.

**Flow cytometry**

All fluorescent signals were detected by BD Accuri^TM^ C6 Plus.

For VHL target engagement assay, 0.5 × 10^6^ BRD4-eGFP_mCherry expressing HeLa cells/well were seeded on 12 well plates. Compounds and vehicles (DMSO) were treated for 4 h after cell adhesion. Cells were harvested and washed once with cold DPBS. GFP signal was detected from 100 μl of resuspended cells in DPBS and normalized to DMSO control.

For apoptosis assay, 0.5 × 10^6^ MOLM-13 cells/well were seeded on 12 well plates, and compounds and vehicle (DMSO) were treated for 48 h. Cells were harvested and washed once with cold DPBS with Ca^2+^ and Mg^2+^. Samples were stained with Annexin V (Invitrogen, A13201) and PI (BD, 556463), and apoptotic cells were analyzed.

For differentiation assay, 0.5 × 10^6^ MOLM-13 cells/well were seeded on 12 well plates, and compounds and vehicle (DMSO) were treated for 48 h. Cells were harvested, and washed once with cold DPBS. After blocking with human Fc block reagent (BD, 564220), samples were stained with CD11b antibody (Invitrogen, #25-0112-82), and differentiated cells were analyzed.

**Isolation and establishment of human gastric cancer organoid**

This study was approved by Yonsei University Hospital Institutional Review Board (Seoul, Korea). All patients provided written informed consent. Tissue pieces obtained from gastric cancer patients were immediately put into ice-cold DMEM/F12 (Gibco-Invitrogen, CA, USA) along with penicillin, primocin. Dissociated cells were transferred to a new tube through a 100-μm cell strainer. The dissociated cells were seeded in Matrigel (Corning Inc., Corning, NY, USA) along with organoid culture media (DMEM/F12 media supplemented with antibiotics, Glutamine, HEPES, conditioned media for Wnts, R-spondins, as well as Noggin, B27, N-acetyl cysteine, nicotinamide, FGF10, Primocin, EGF, Gastrin, Y-27632, and A83-01)

**Organoid passing and freezing**

Medium for PDOs were replaced every 2 days, and passage was performed mainly by mechanical dissociation using 29-31G syringe every 10-14 days, at a ratio of 1:3-5. The stocks of PDOs were prepared in freezing media and stored in liquid nitrogen after retaining for 2 days at -80°C.

**Organoid viability test**

The growth curve of PDOs was examined by the Cell Titer-Glo assay (Promega, USA) according to the manufacturer’s instructions. On the indicated day between day 0-12, the cell viability assay was performed and the luminescence was quantified by the luminometer (Molecular devices, USA). For drug treatment, 1000 PDOs/well were seeded. After cultivating for 48 h, the PDOs were treated 8 drug concentrations. After 5 days, cell viability assay according to the manufacturer’s protocol. IC50 values were calculated by GraphPadPrism Ver.5.

**Hotspot RNA methyltransferase assay**

METTL3/14 methyltransferase activity was measured by Reaction Biology according to RBC Hotspot RNA methyltransferase assay protocol.

**^1^H and ^13^C NMR spectra of intermediates.**

Methyl 6-((6-chloropyrimidin-4-yl)amino)hexanoate (**3**).

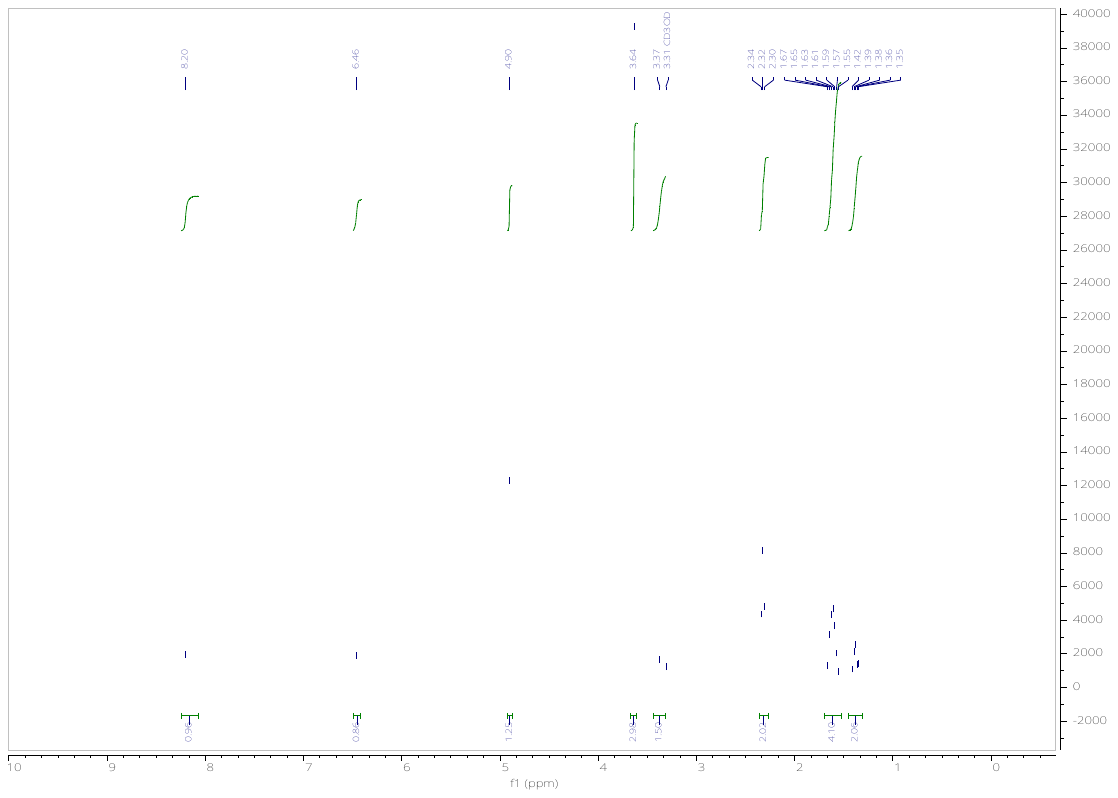

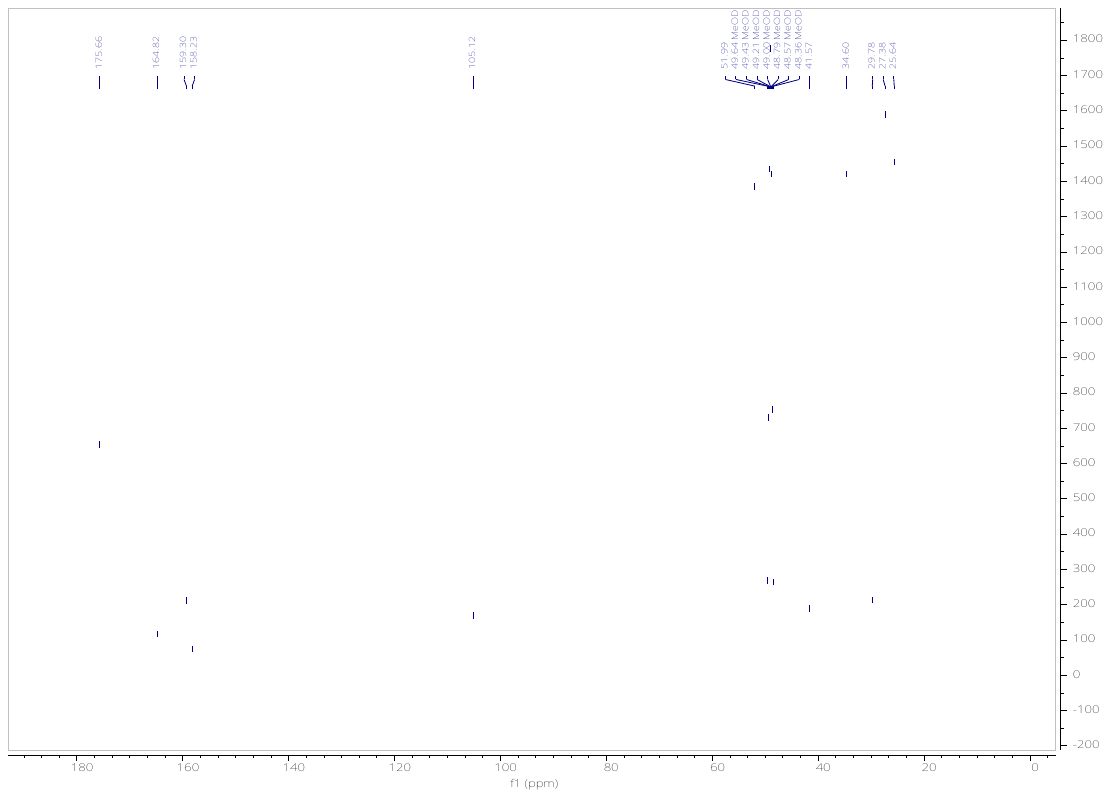

Methyl 8-((6-chloropyrimidin-4-yl)amino)octanoate (**4**).

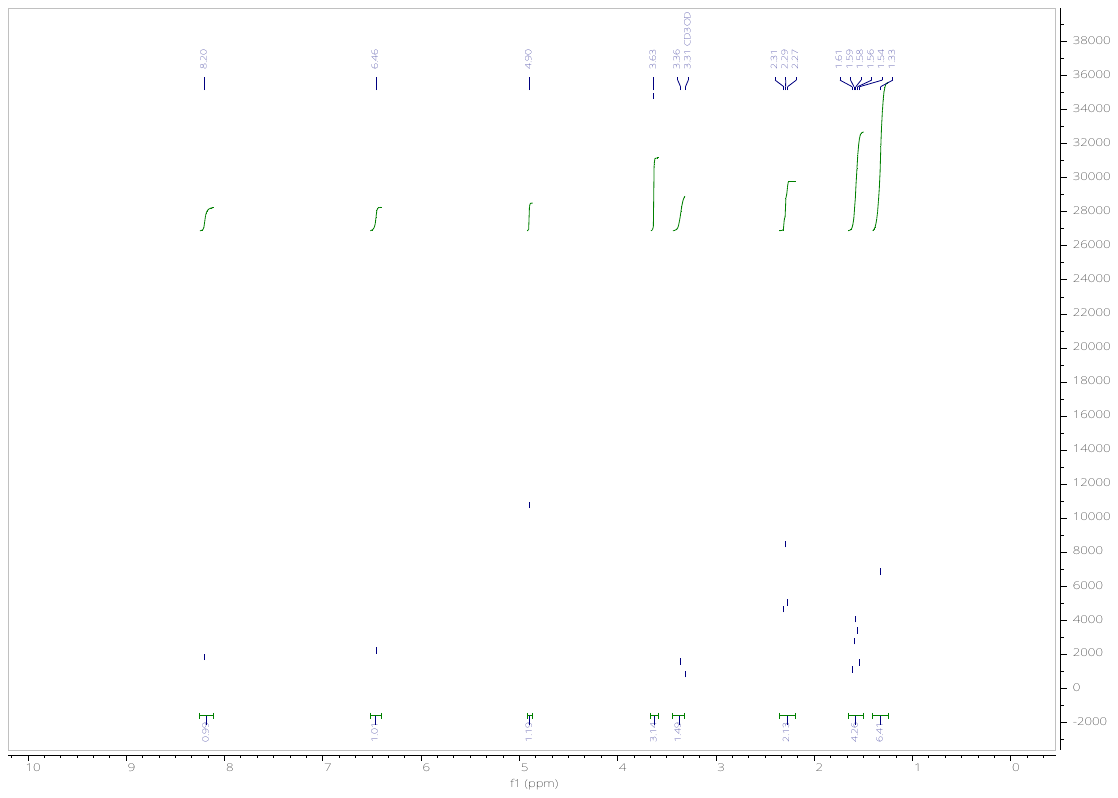

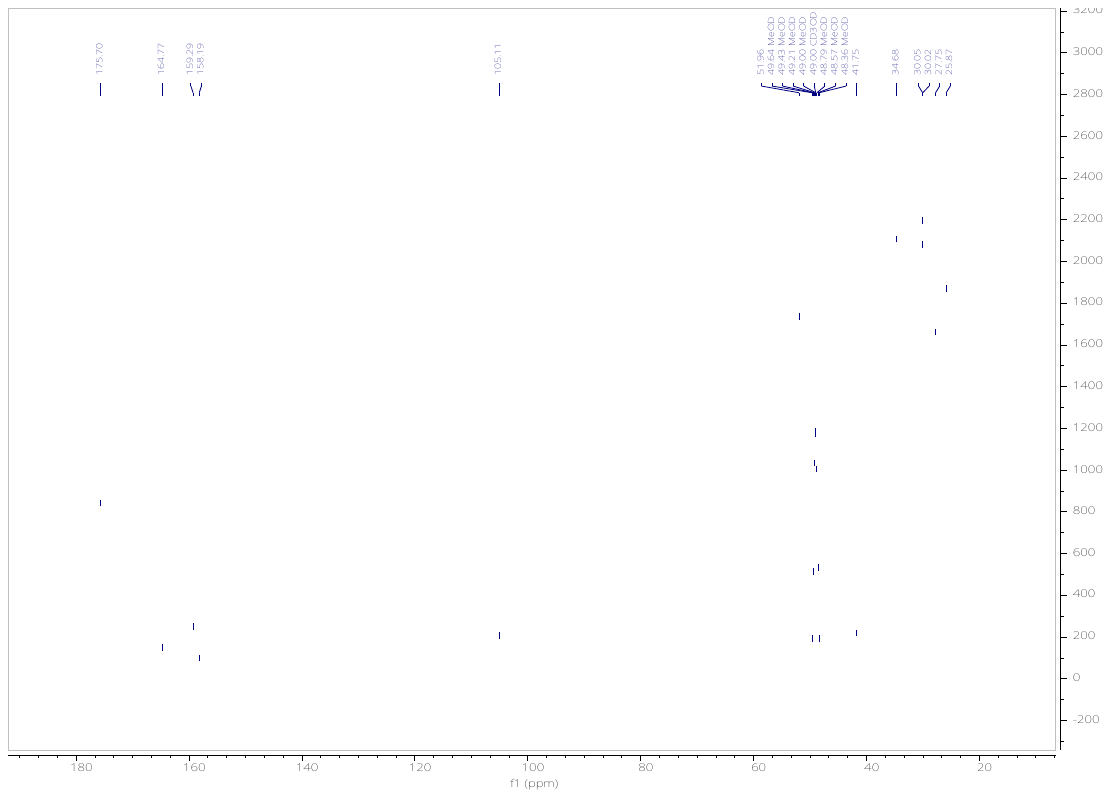

Methyl 11-((6-chloropyrimidin-4-yl)amino)undecanoate (**5**).

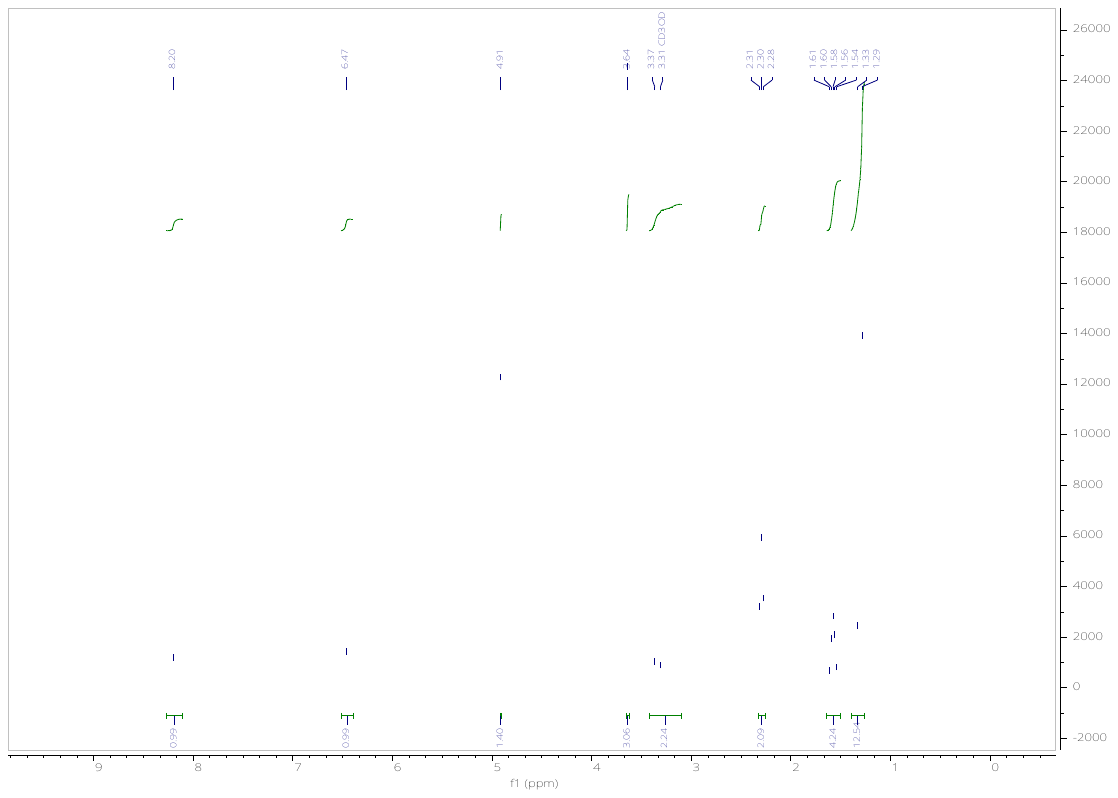

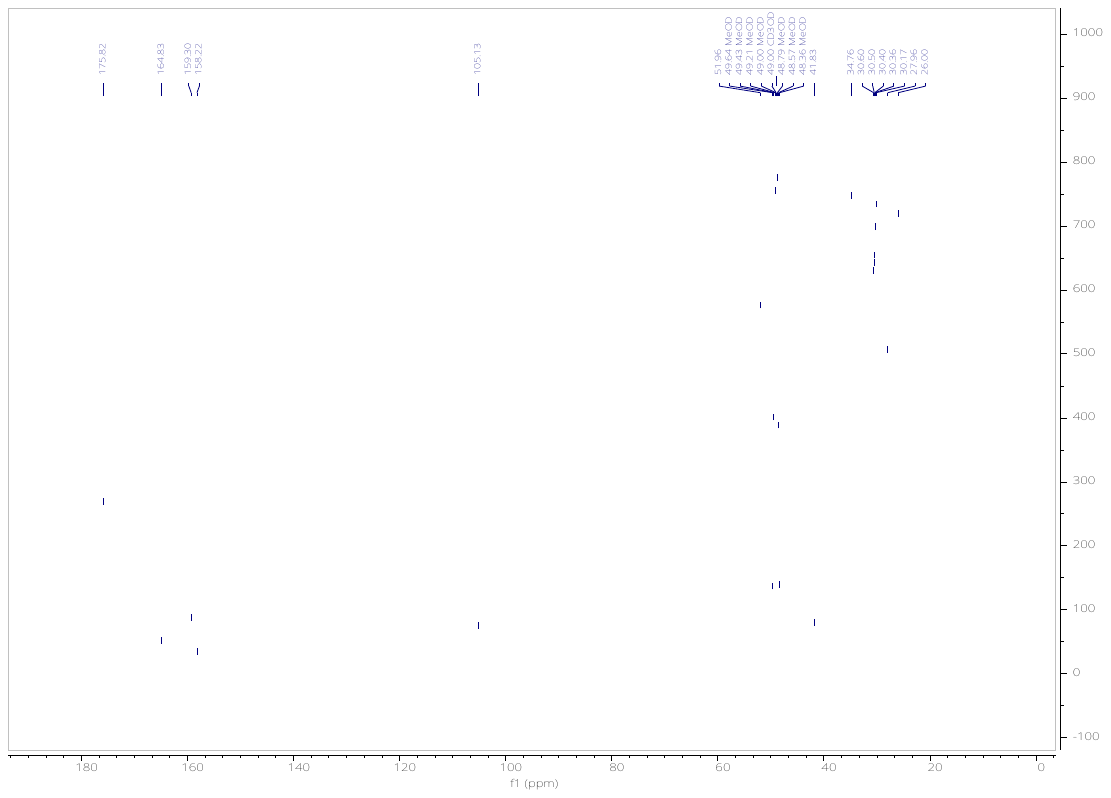

Methyl 6-(N-(6-chloropyrimidin-4-yl)acetamido)hexanoate (**6**).

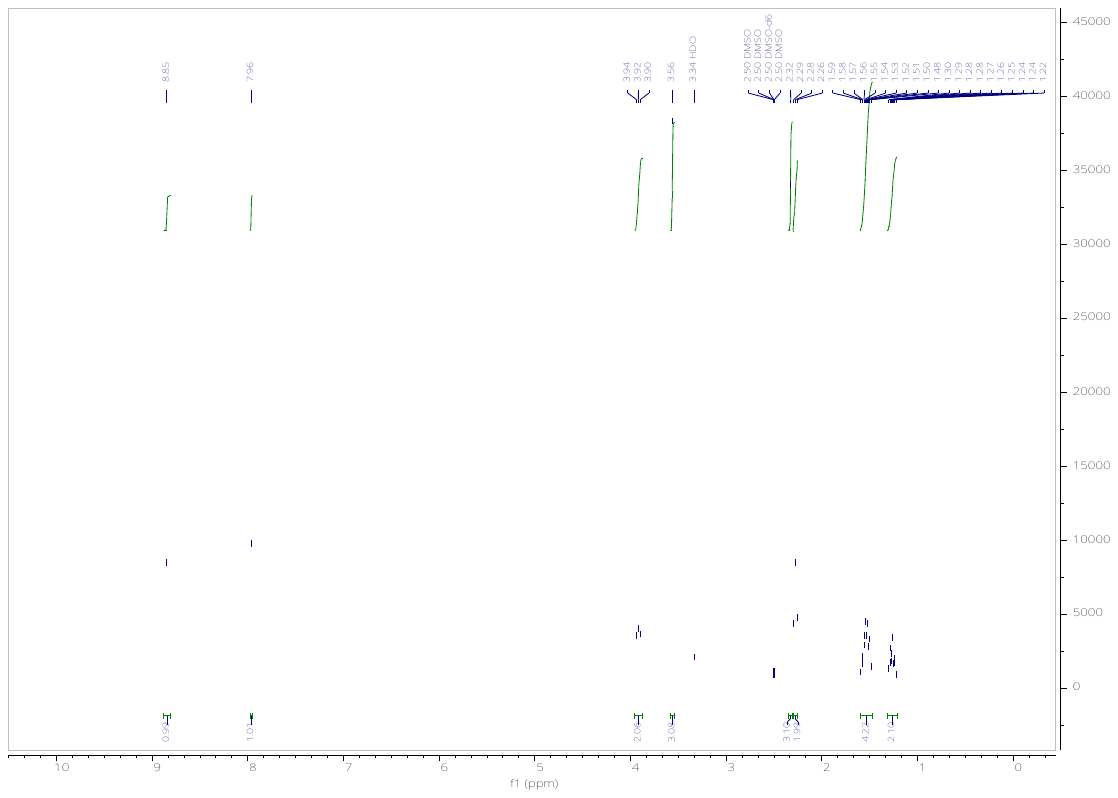

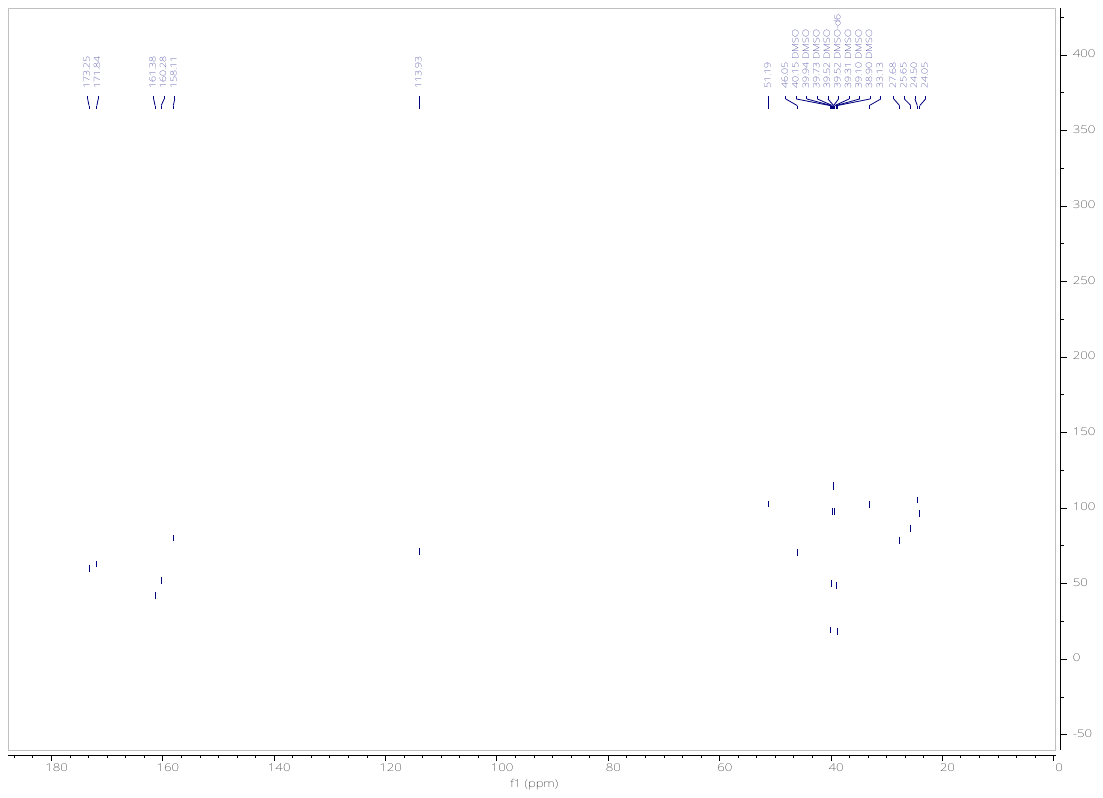

Methyl 8-(N-(6-chloropyrimidin-4-yl)acetamido)octanoate (**7**).

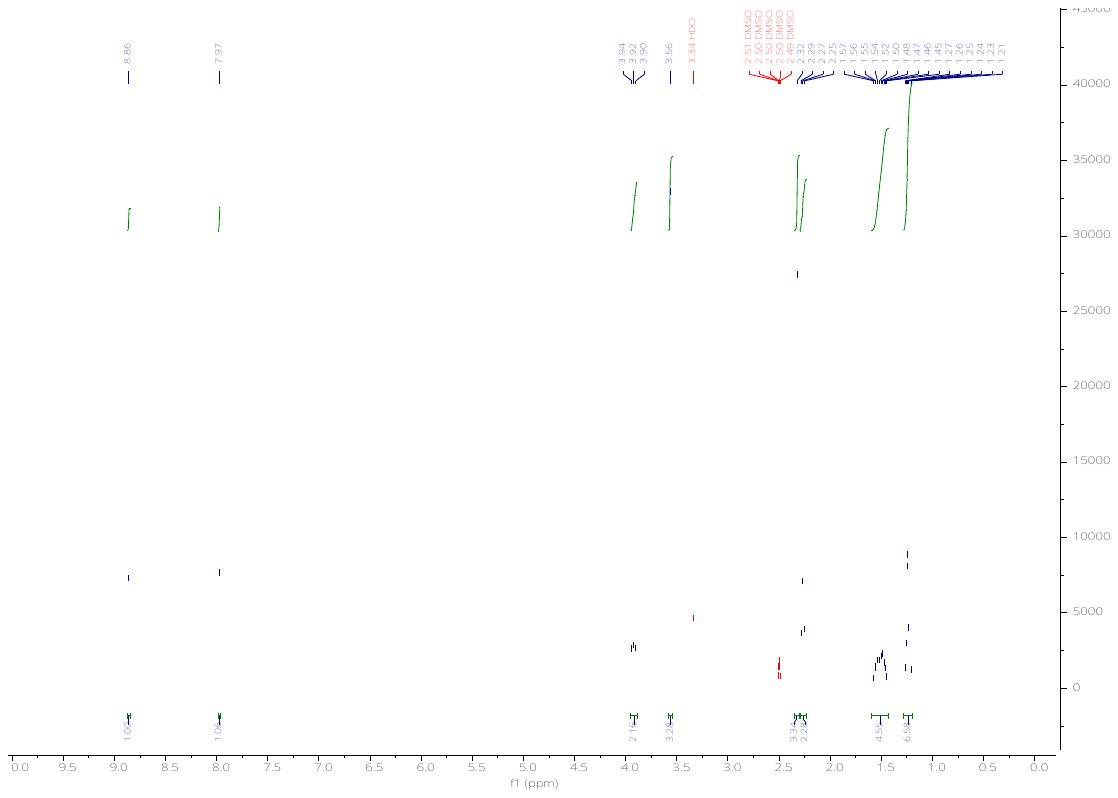

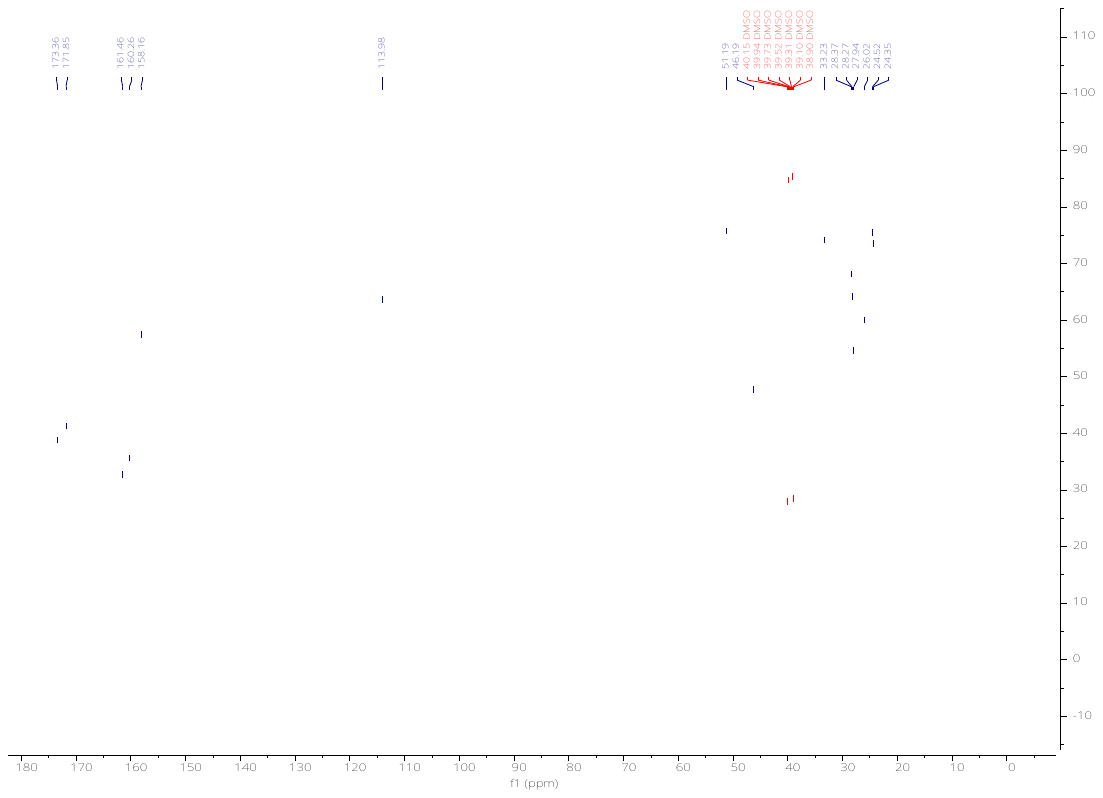

Methyl 11-(N-(6-chloropyrimidin-4-yl)acetamido)undecanoate (**8**).

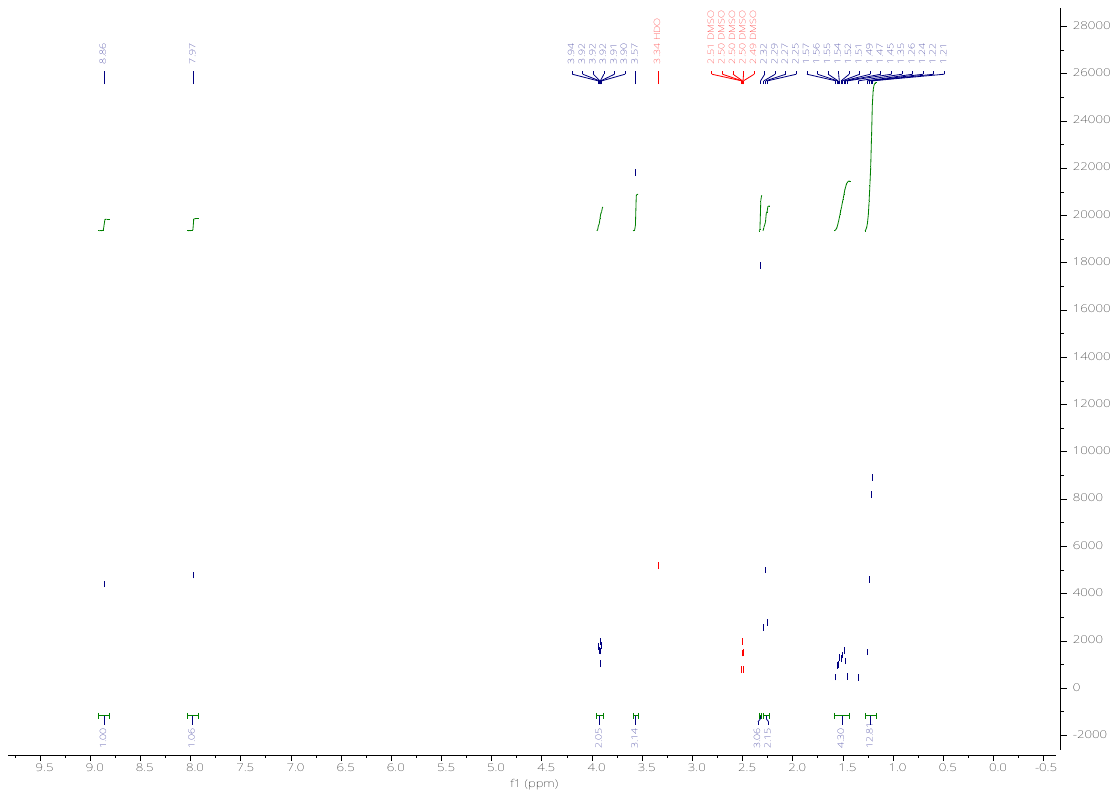

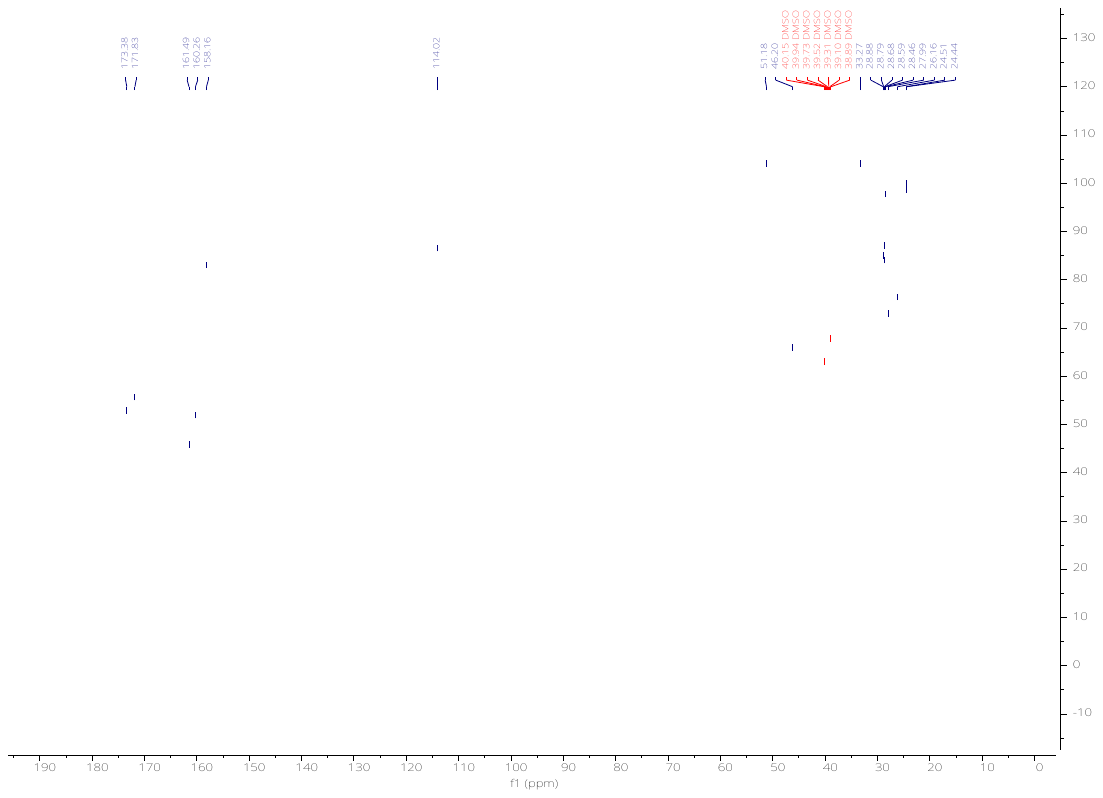

*tert*-Butyl 3-(2-((6-chloropyrimidin-4-yl)amino)ethoxy)propanoate (**9**).

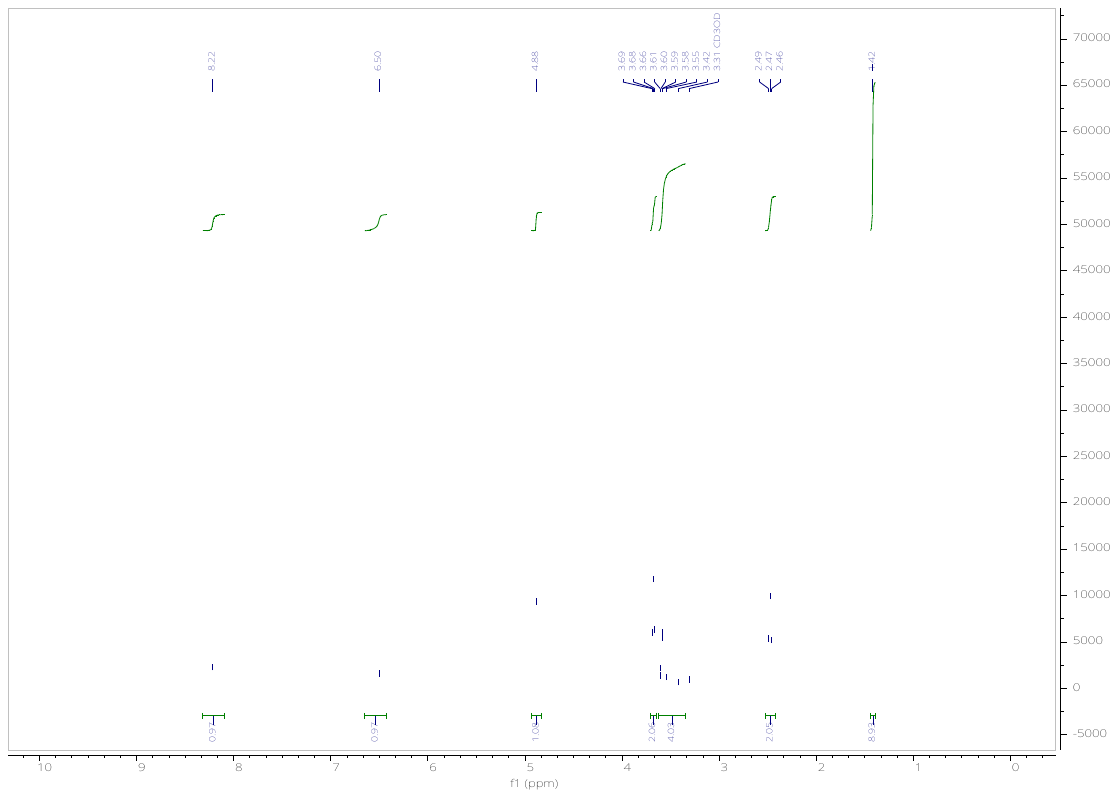

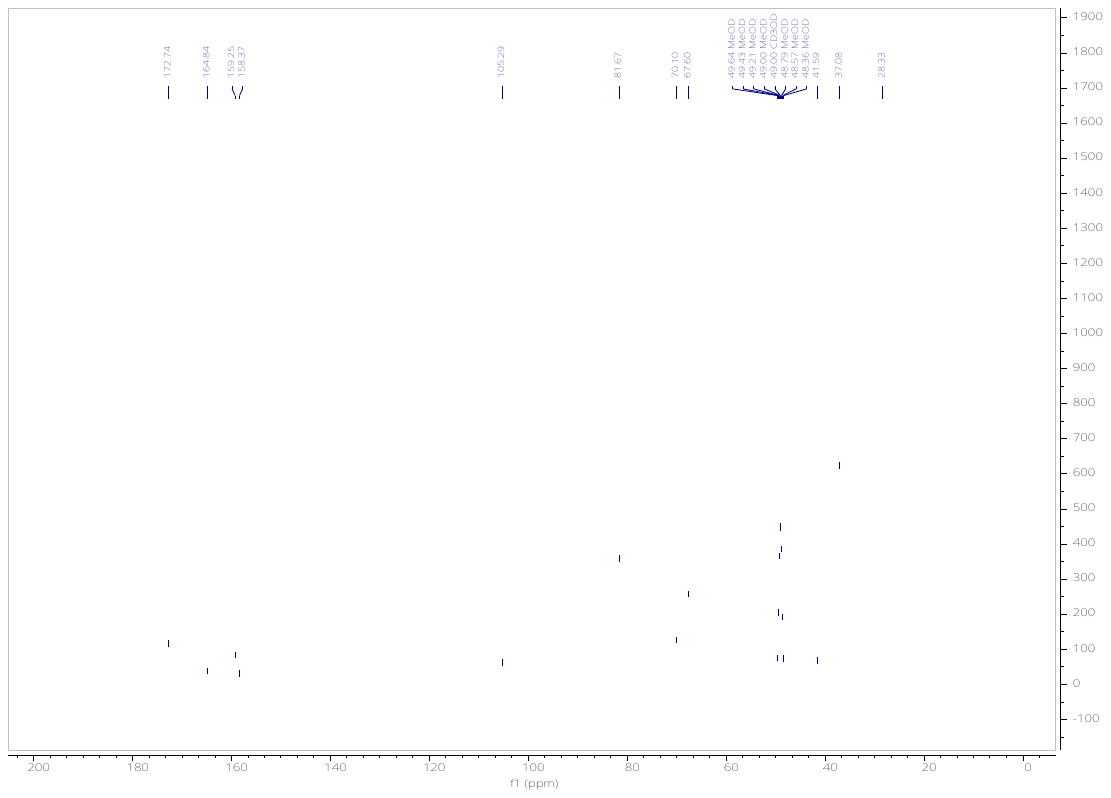

*tert*-Butyl 3-(2-(2-((6-chloropyrimidin-4-yl)amino)ethoxy)ethoxy)propanoate (**10**).

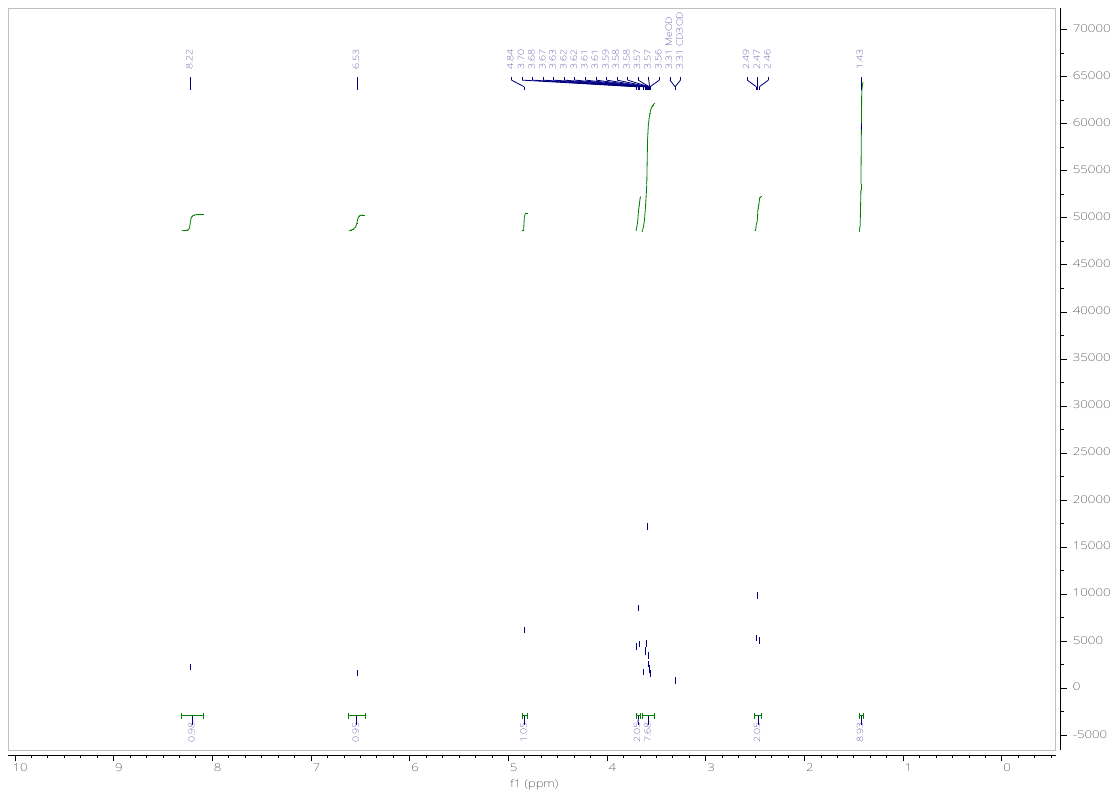

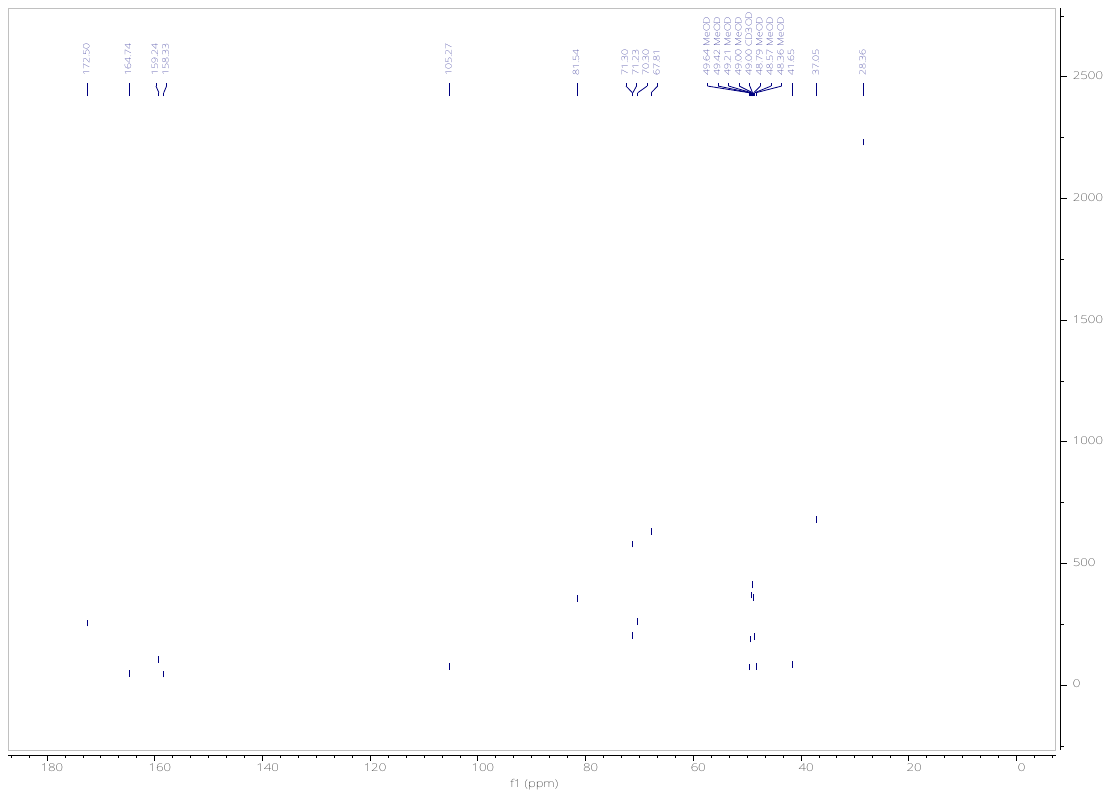

*tert*-Butyl 3-(2-(2-(2-((6-chloropyrimidin-4-yl)amino)ethoxy)ethoxy)ethoxy)propanoate (**11**).

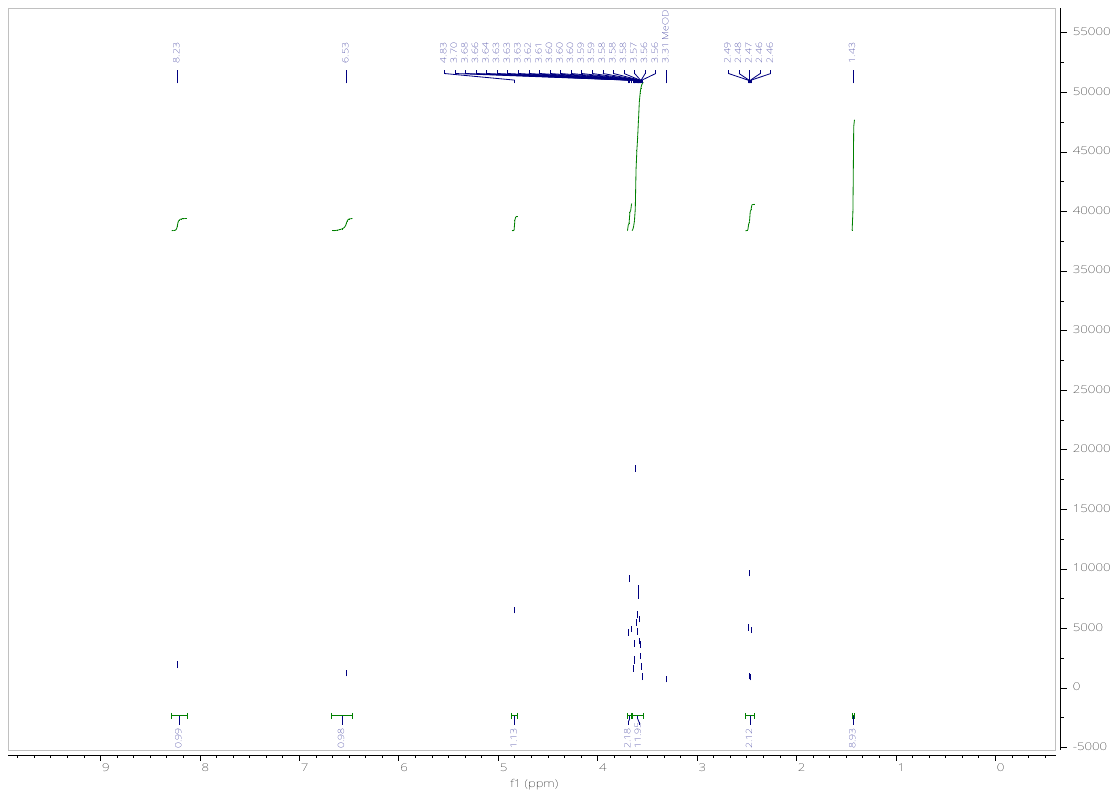

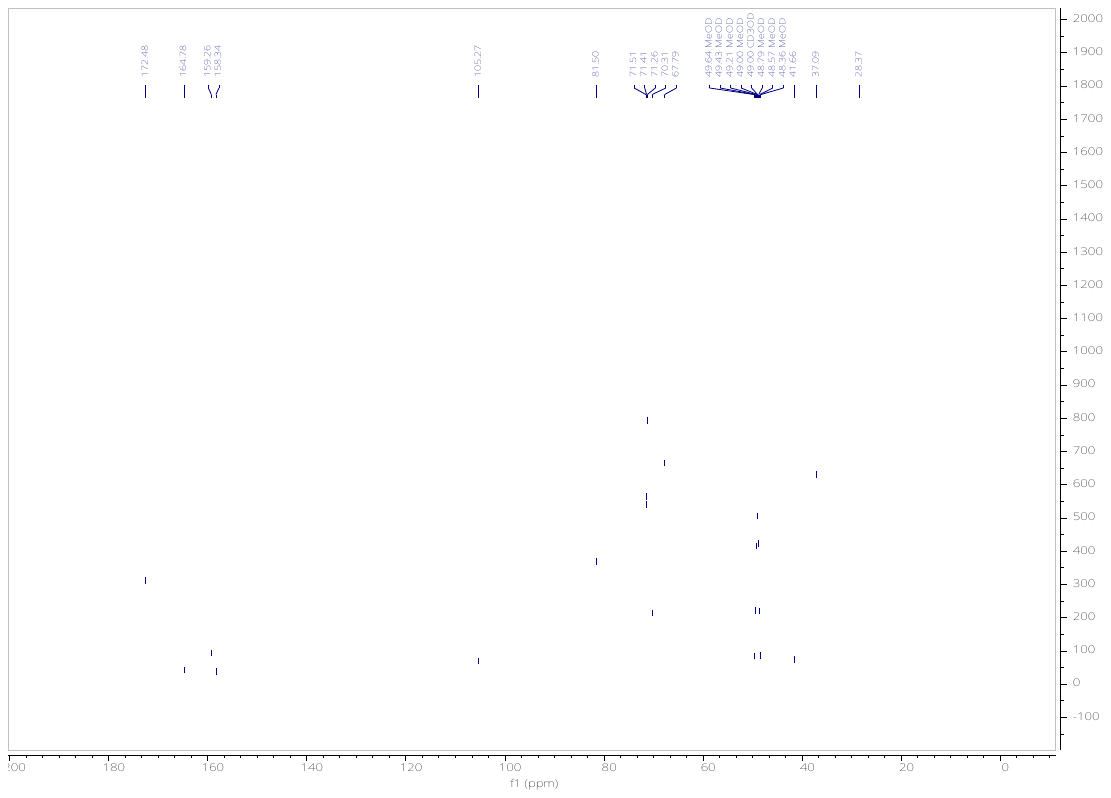

*tert*-Butyl 3-(2-(N-(6-chloropyrimidin-4-yl)acetamido)ethoxy)propanoate (**12**).

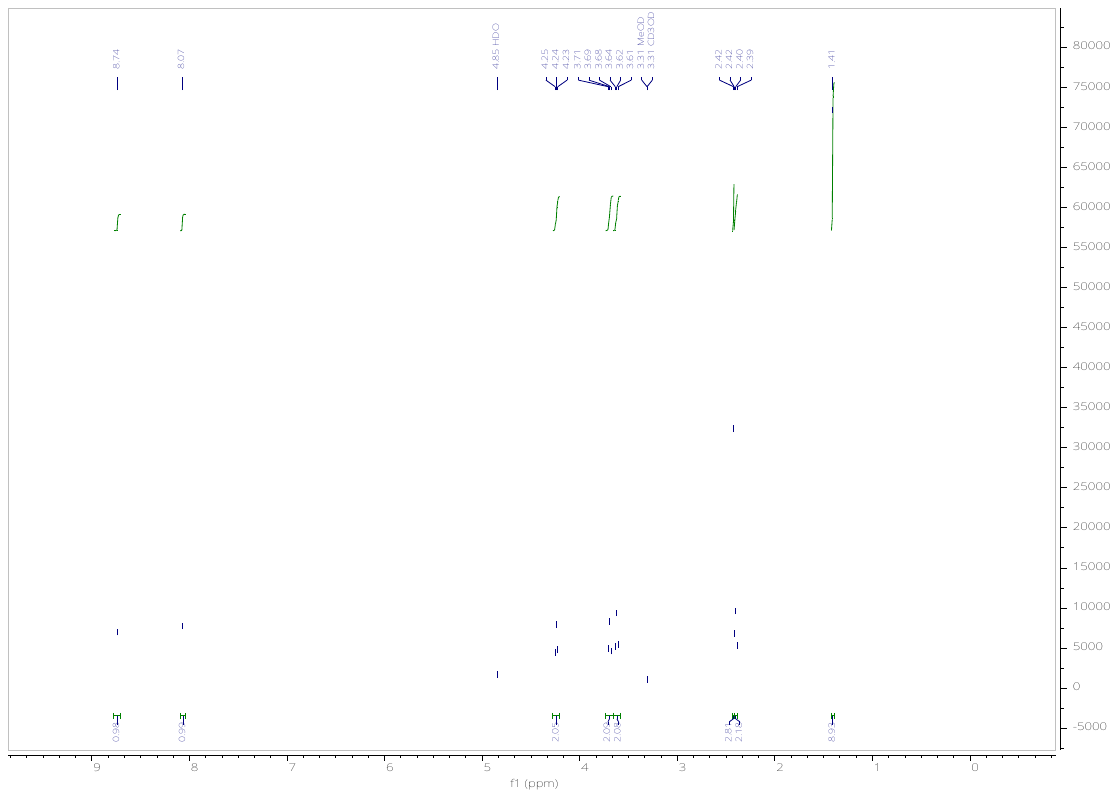

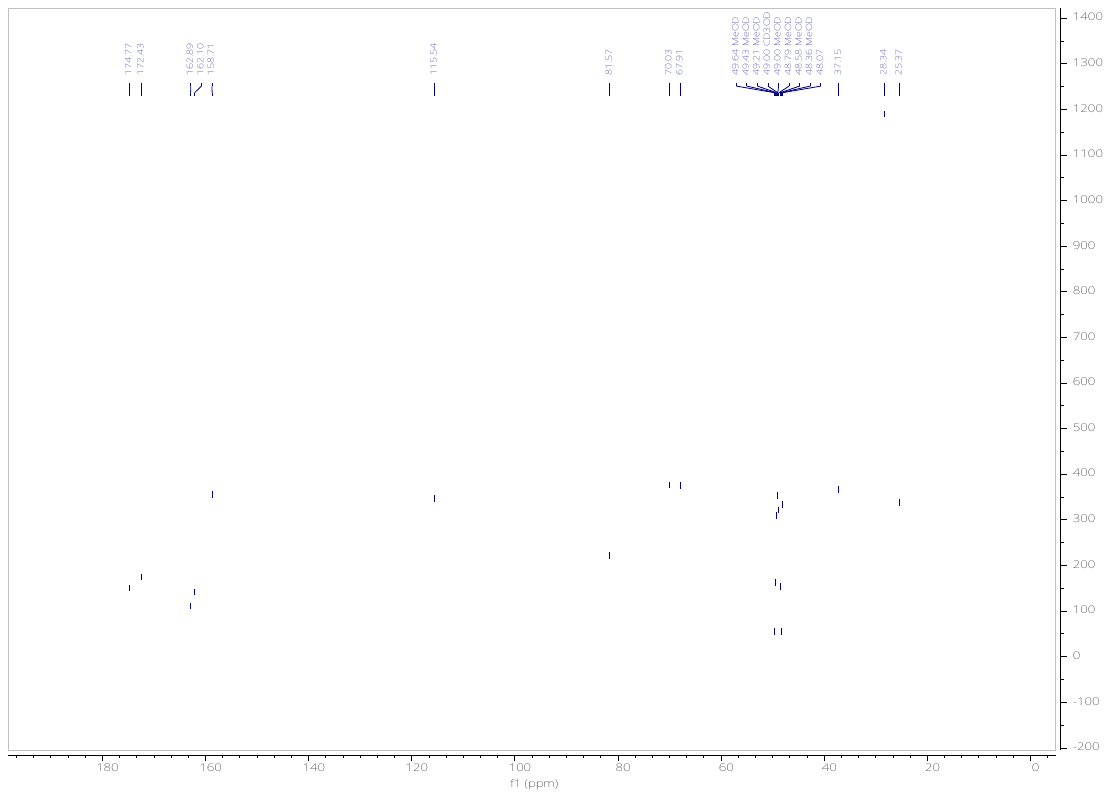

*tert-*Butyl 3-(2-(2-(N-(6-chloropyrimidin-4-yl)acetamido)ethoxy)ethoxy)propanoate (**13**).

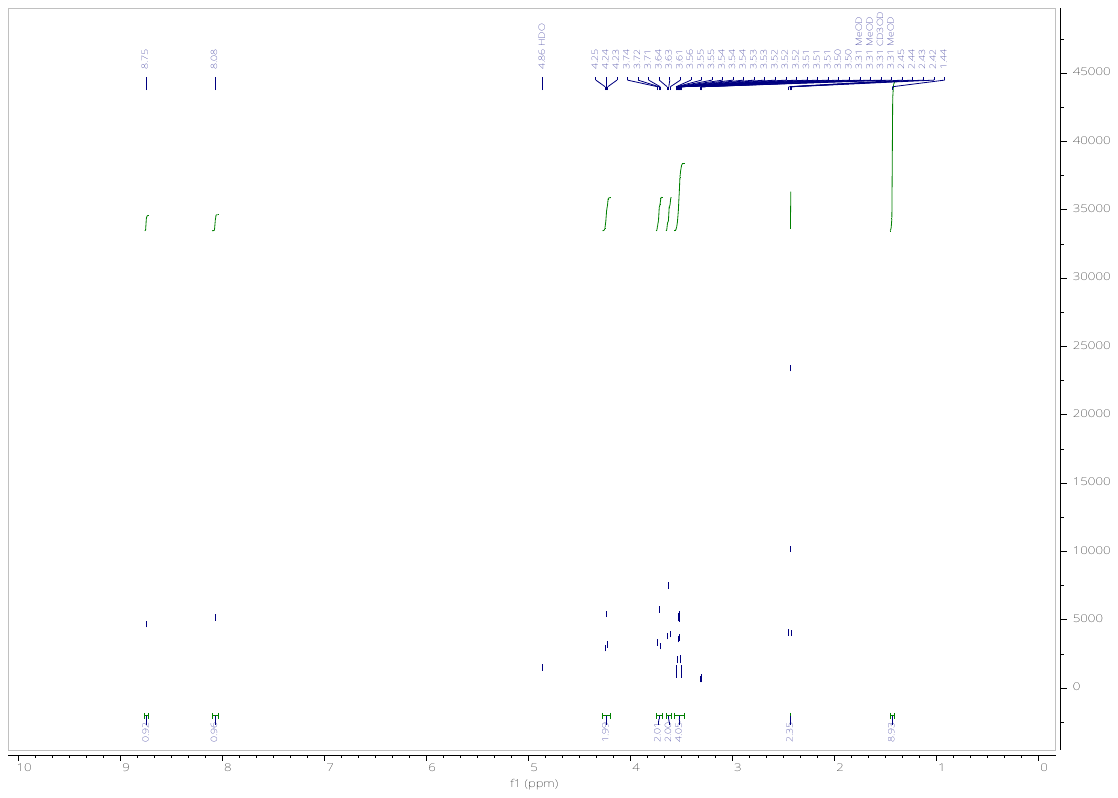

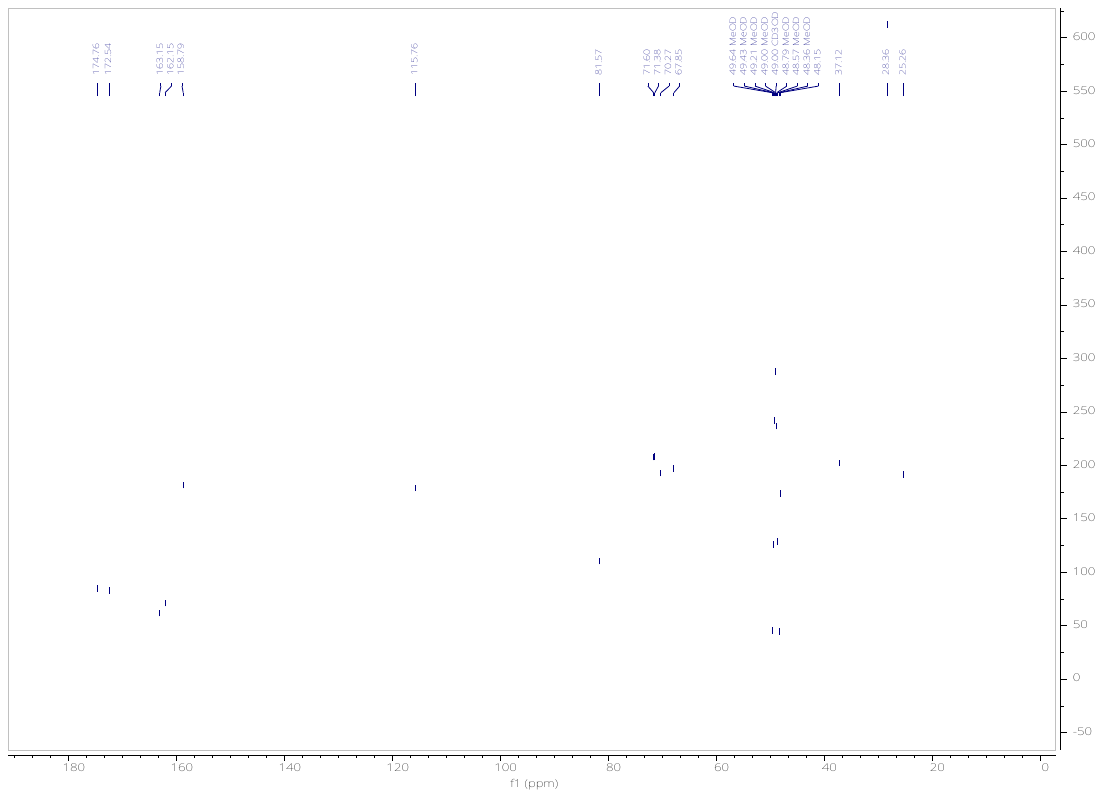

*tert*-Butyl 3-(6-chloropyrimidin-4-yl)-2-oxo-6,9,12-trioxa-3-azapentadecan-15-oate (**14**).

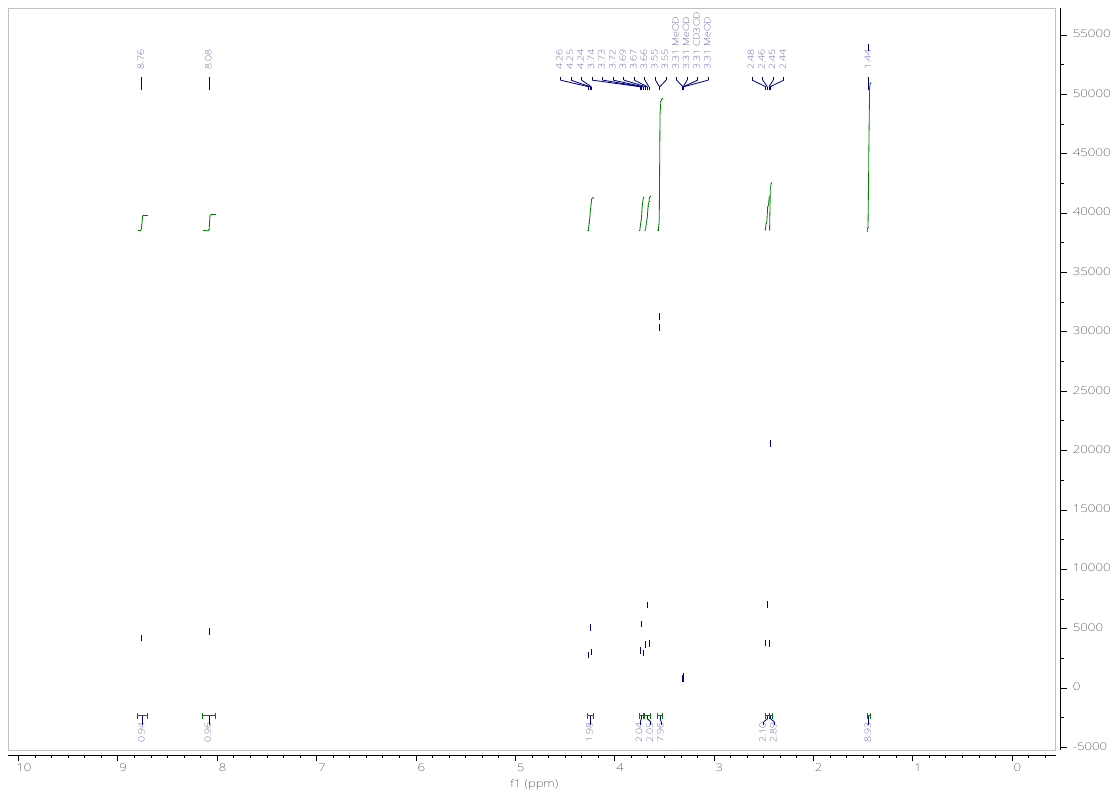

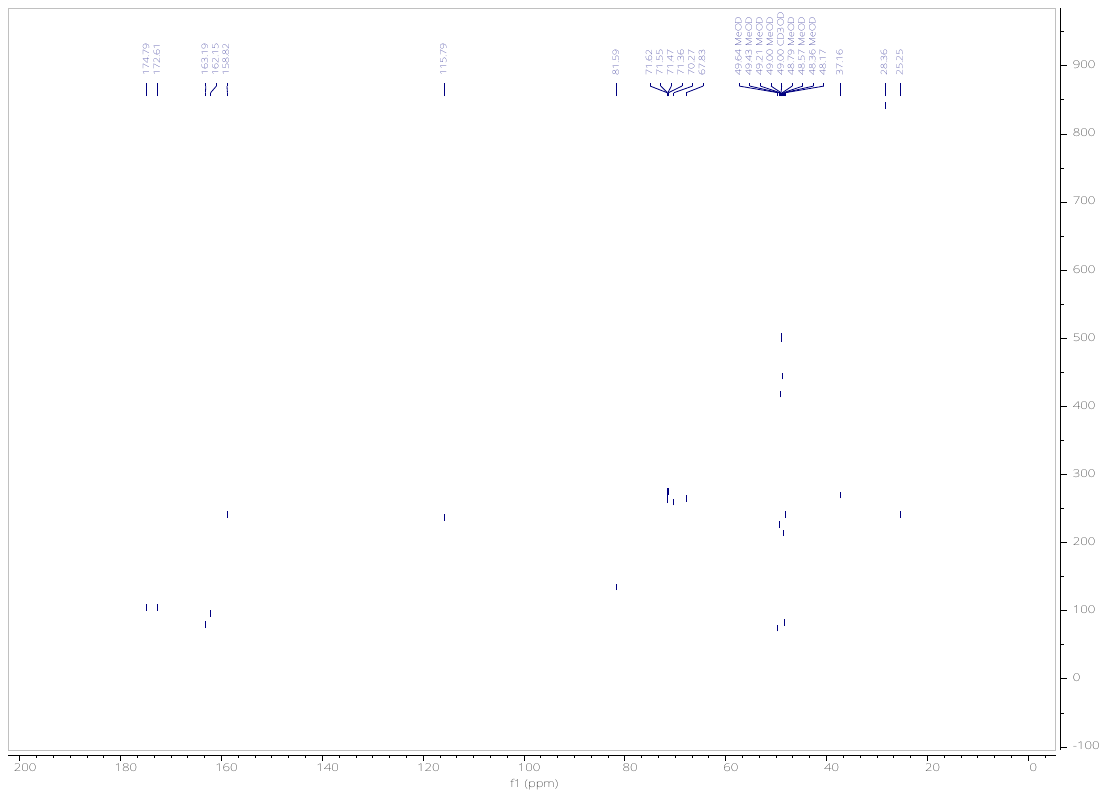

*tert*-butyl 4-(2-(2,6-dioxopiperidin-3-yl)-1,3-dioxoisoindolin-5-yl)piperazine-1-carboxylate (**16**).

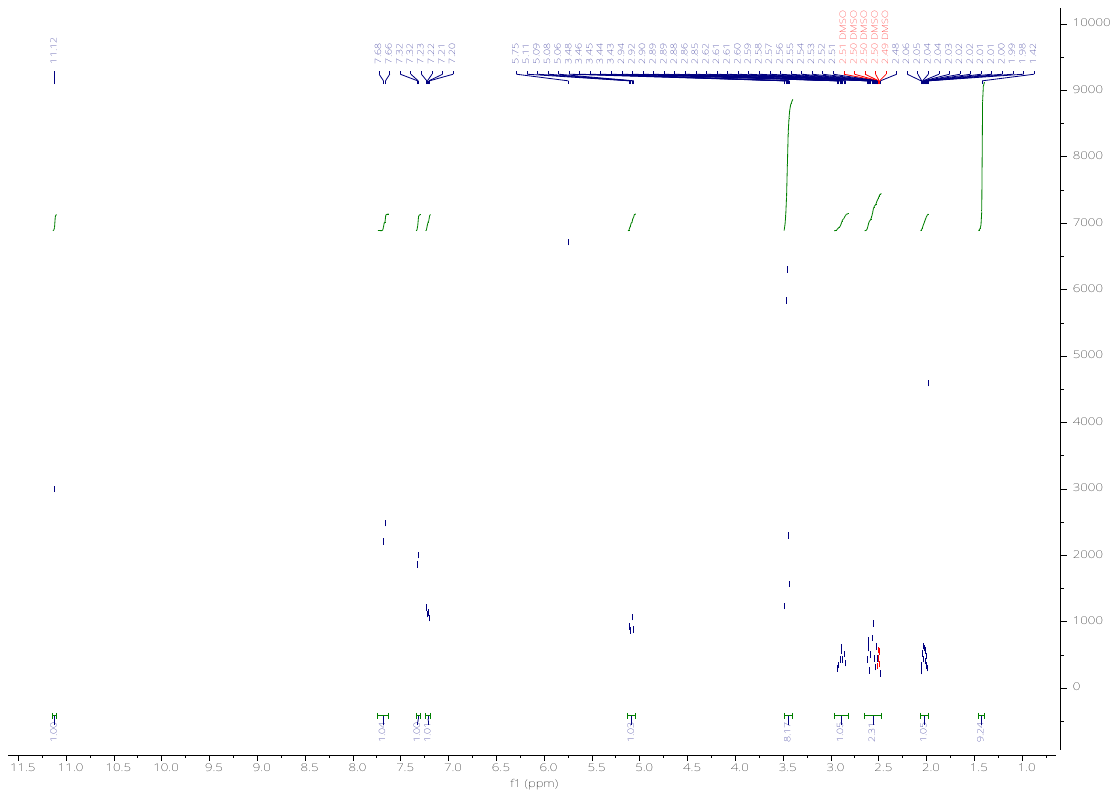

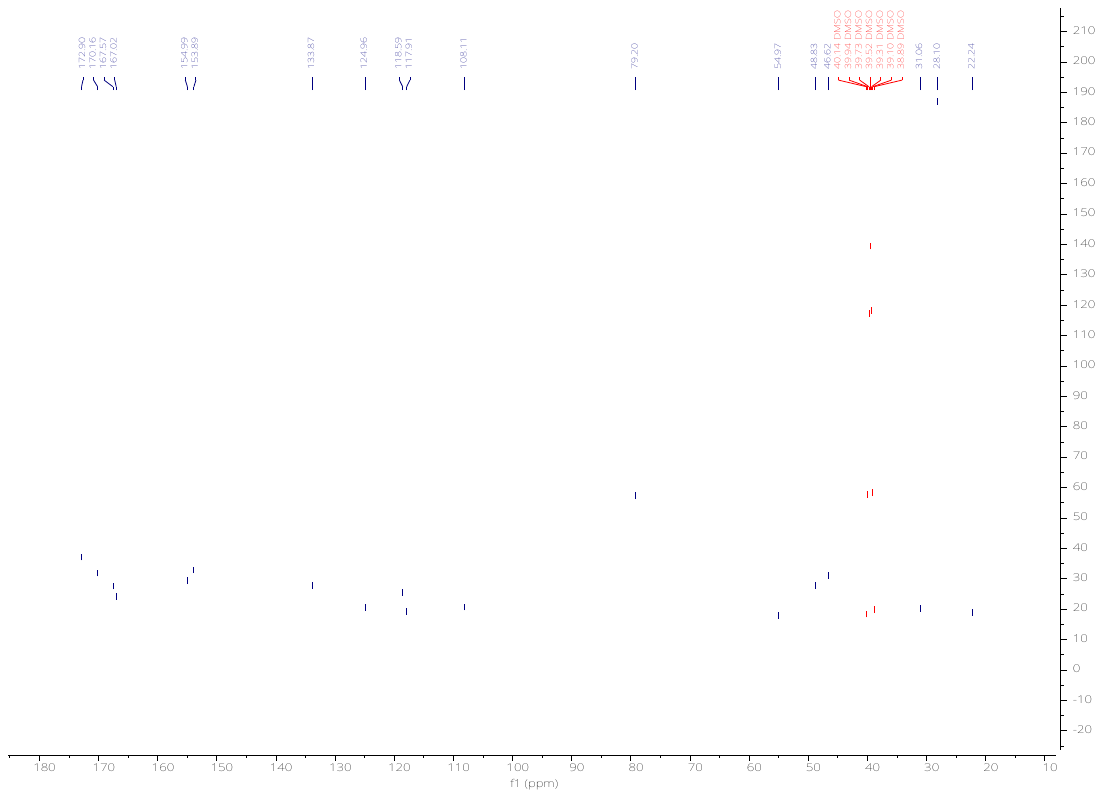

Methyl 6-(N-(6-(4-(4-((4,4-dimethylpiperidin-1-yl)methyl)-2,5-difluorophenyl)-2-oxo-1,4,9-triazaspiro[5.5]undecan-9-yl)pyrimidin-4-yl)acetamido)hexanoate (**18**).

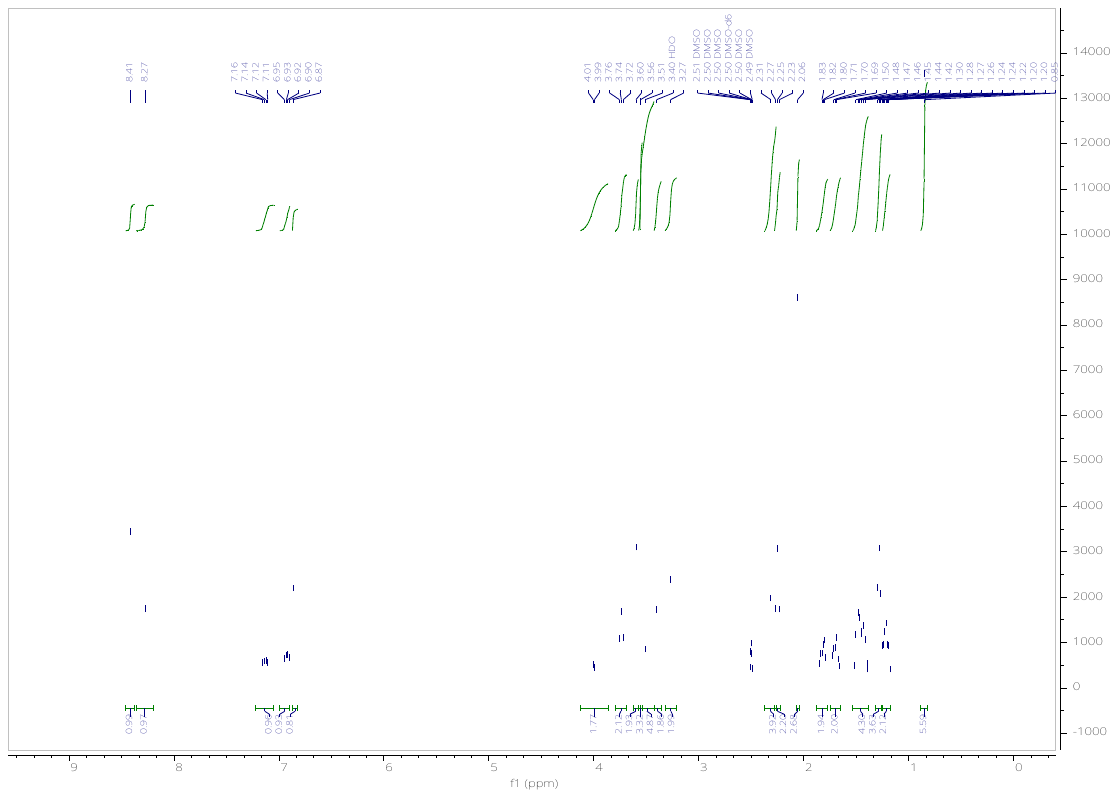

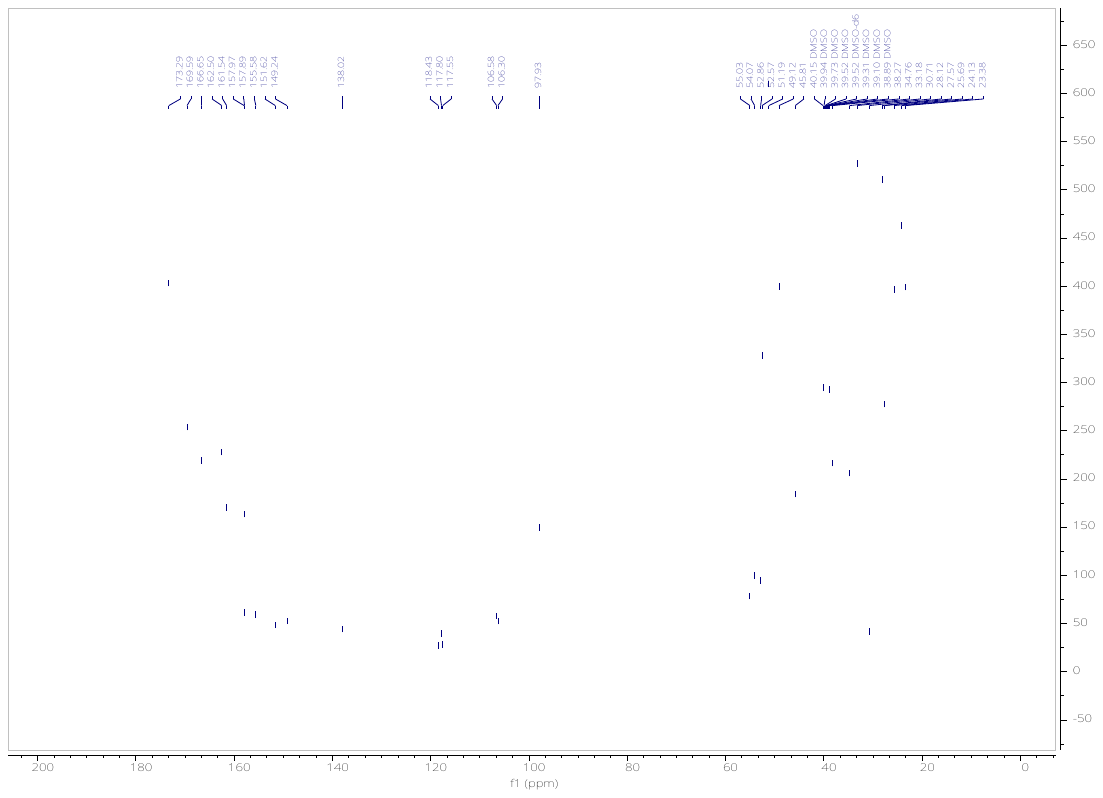

Methyl 8-(N-(6-(4-(4-((4,4-dimethylpiperidin-1-yl)methyl)-2,5-difluorophenyl)-2-oxo-1,4,9-triazaspiro[5.5]undecan-9-yl)pyrimidin-4-yl)acetamido)octanoate (**19**).

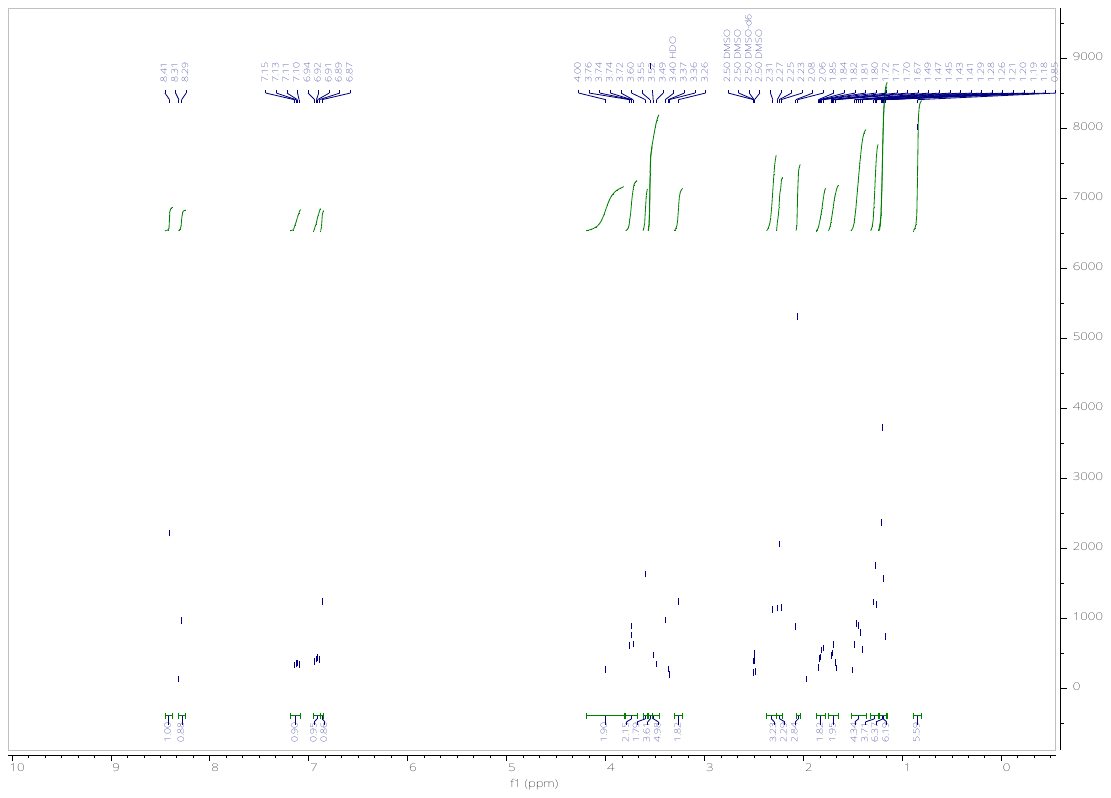

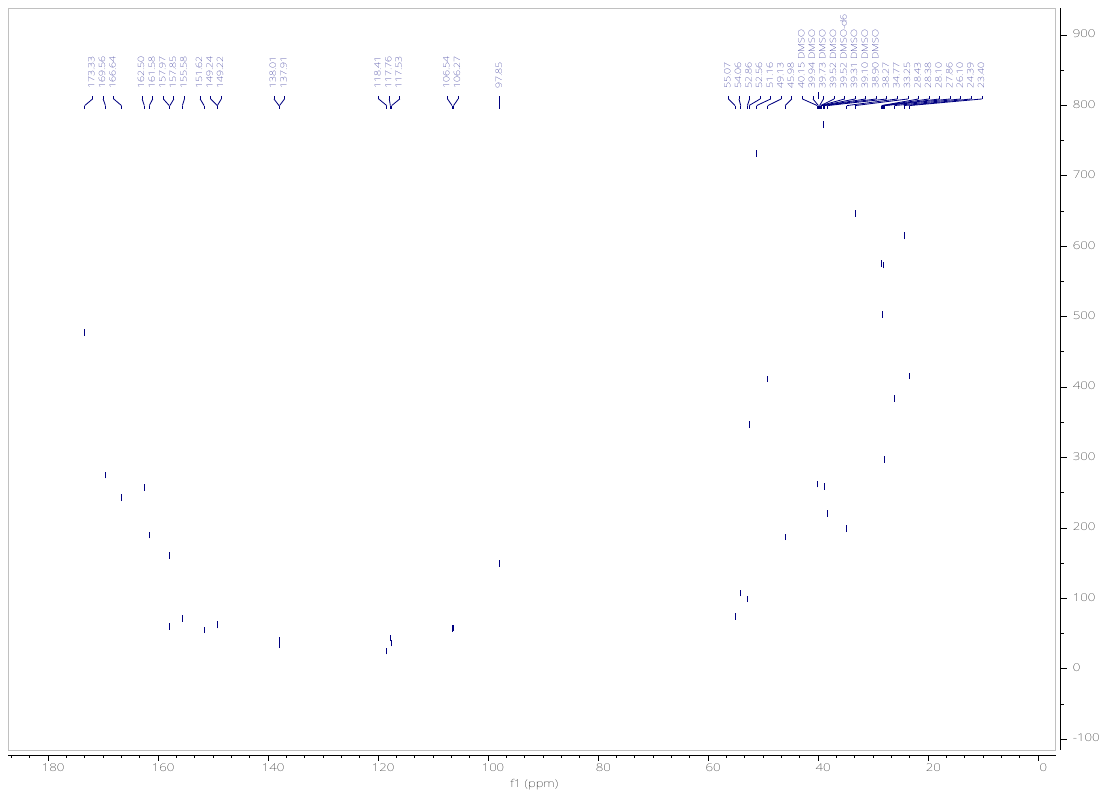

Methyl 11-(N-(6-(4-(4-((4,4-dimethylpiperidin-1-yl)methyl)-2,5-difluorophenyl)-2-oxo-1,4,9-triazaspiro[5.5]undecan-9-yl)pyrimidin-4-yl)acetamido)undecanoate (**20**).

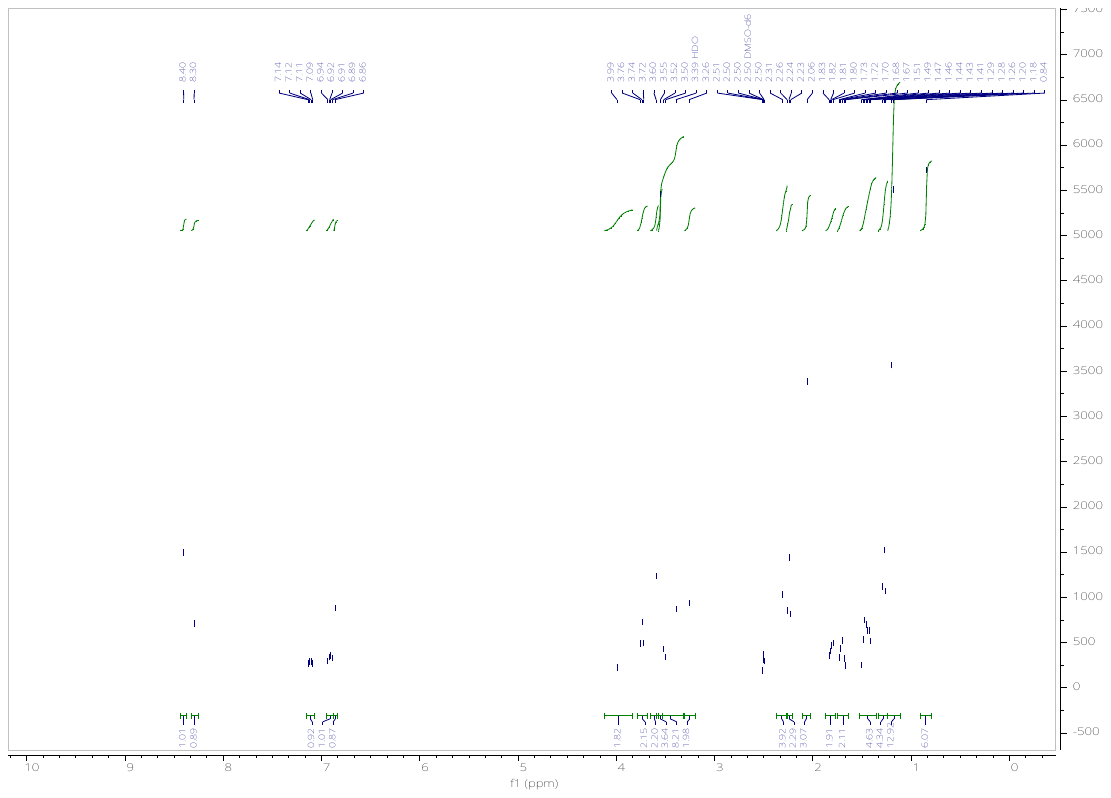

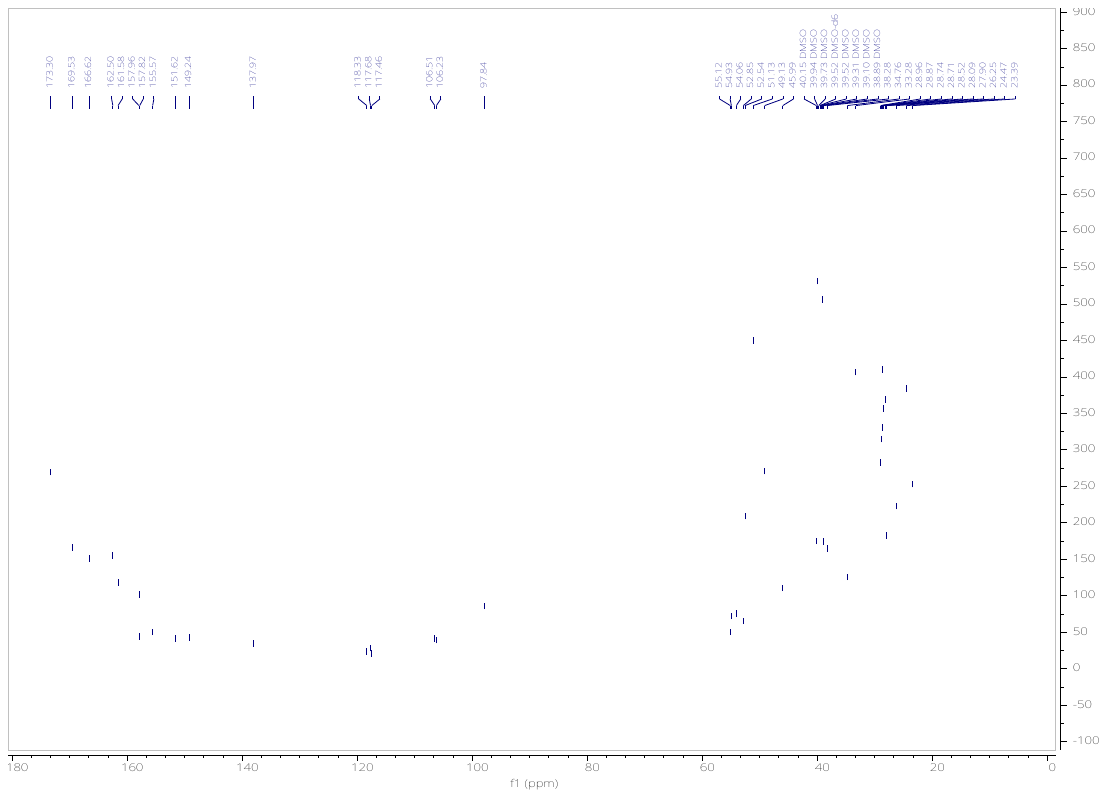

*tert*-Butyl 3-(2-(N-(6-(4-(4-((4,4-dimethylpiperidin-1-yl)methyl)-2,5-difluorophenyl)-2-oxo-1,4,9-triazaspiro[5.5]undecan-9-yl)pyrimidin-4-yl)acetamido)ethoxy)propanoate (**21**).

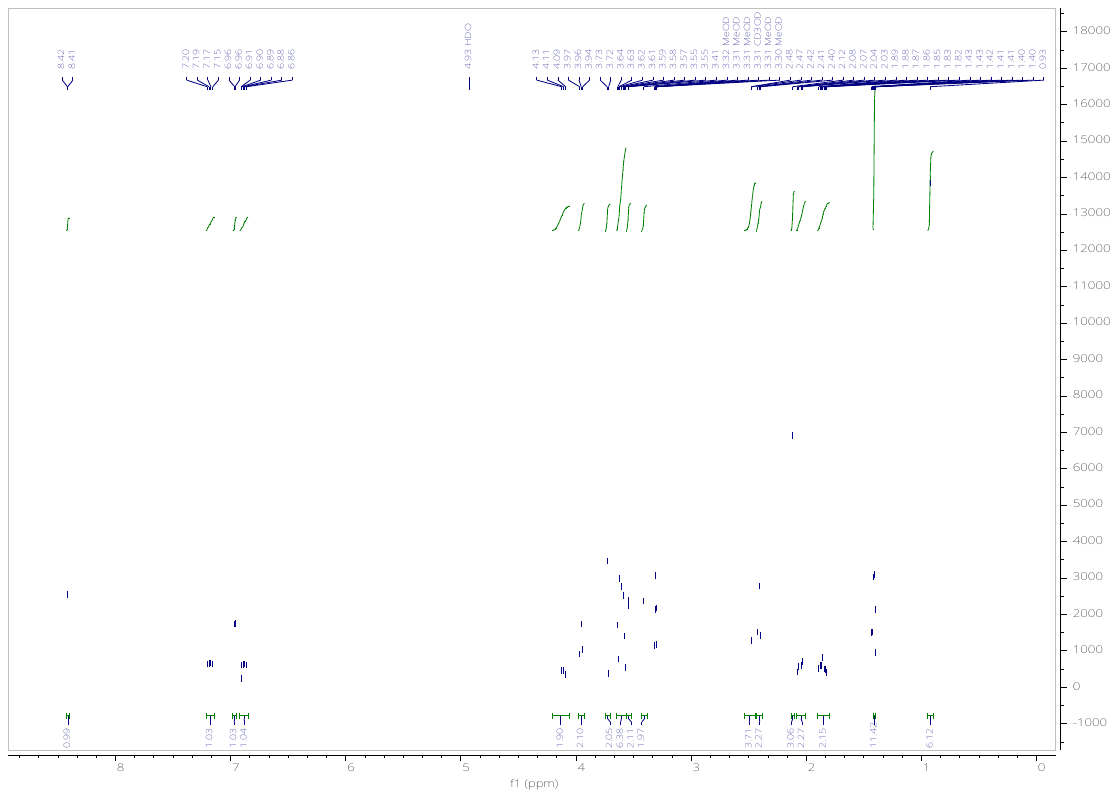

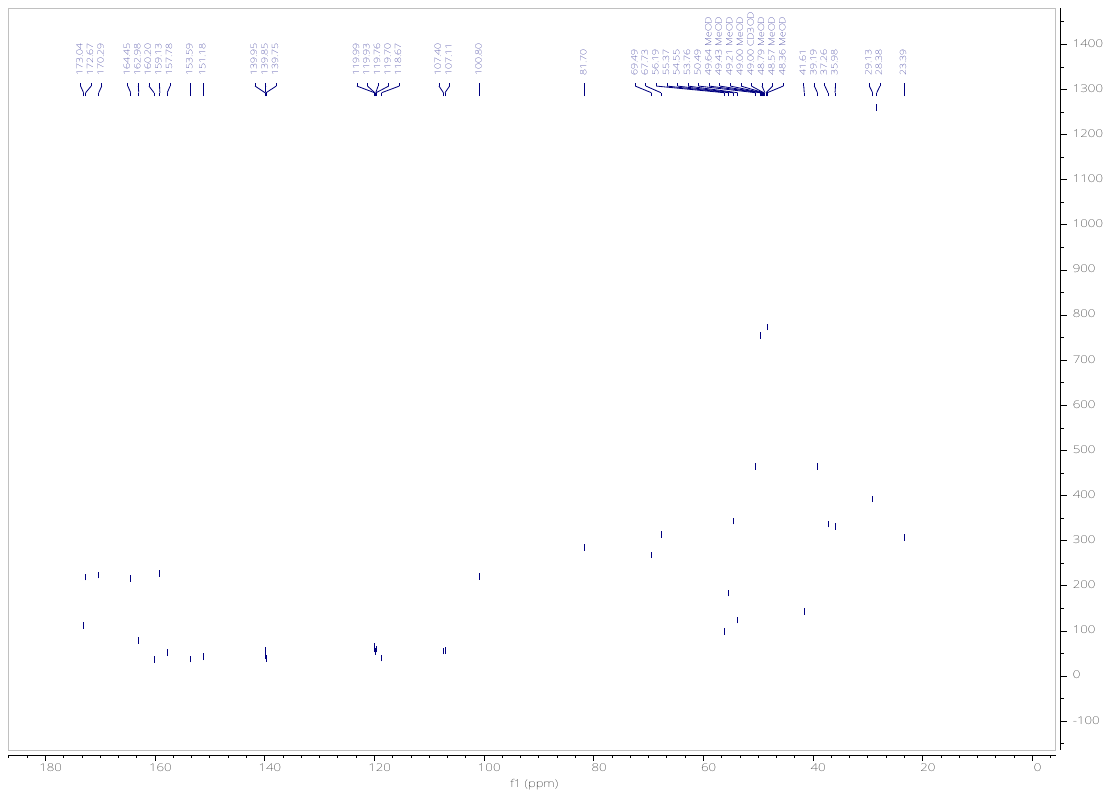

*tert*-Butyl 3-(2-(2-(N-(6-(4-(4-((4,4-dimethylpiperidin-1-yl)methyl)-2,5-difluorophenyl)-2-oxo-1,4,9-triazaspiro[5.5]undecan-9-yl)pyrimidin-4-yl)acetamido)ethoxy)ethoxy)propanoate (**22**).

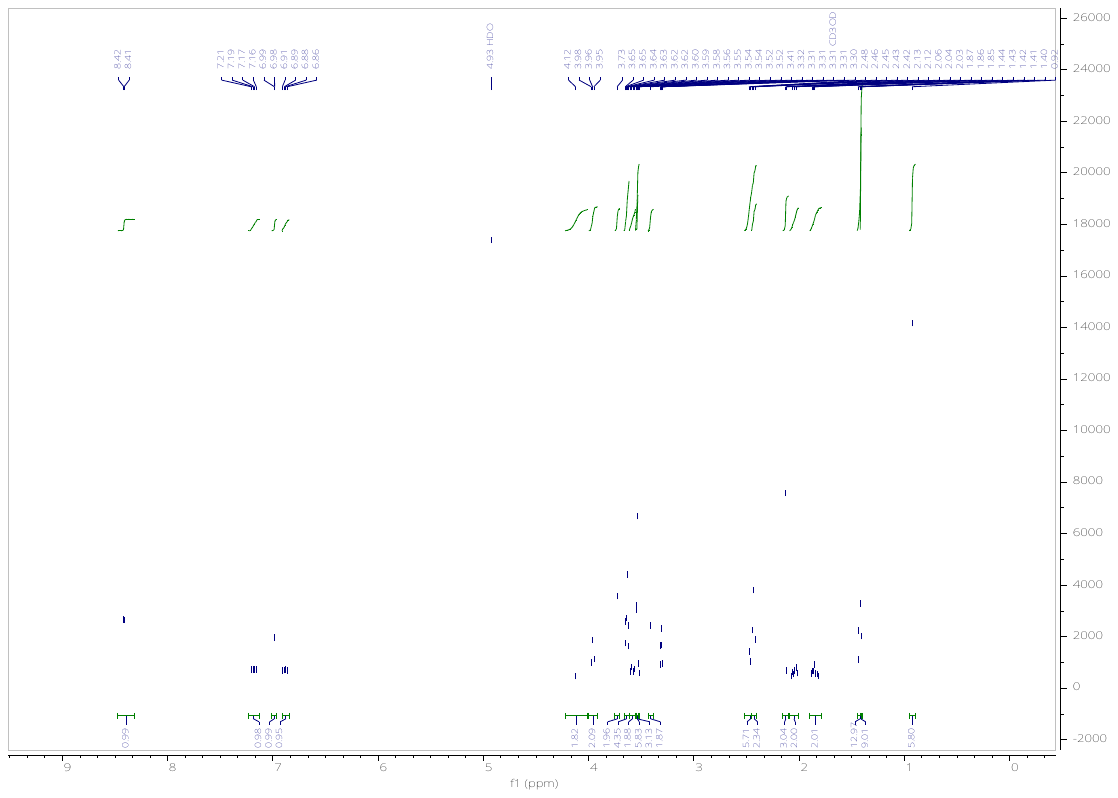

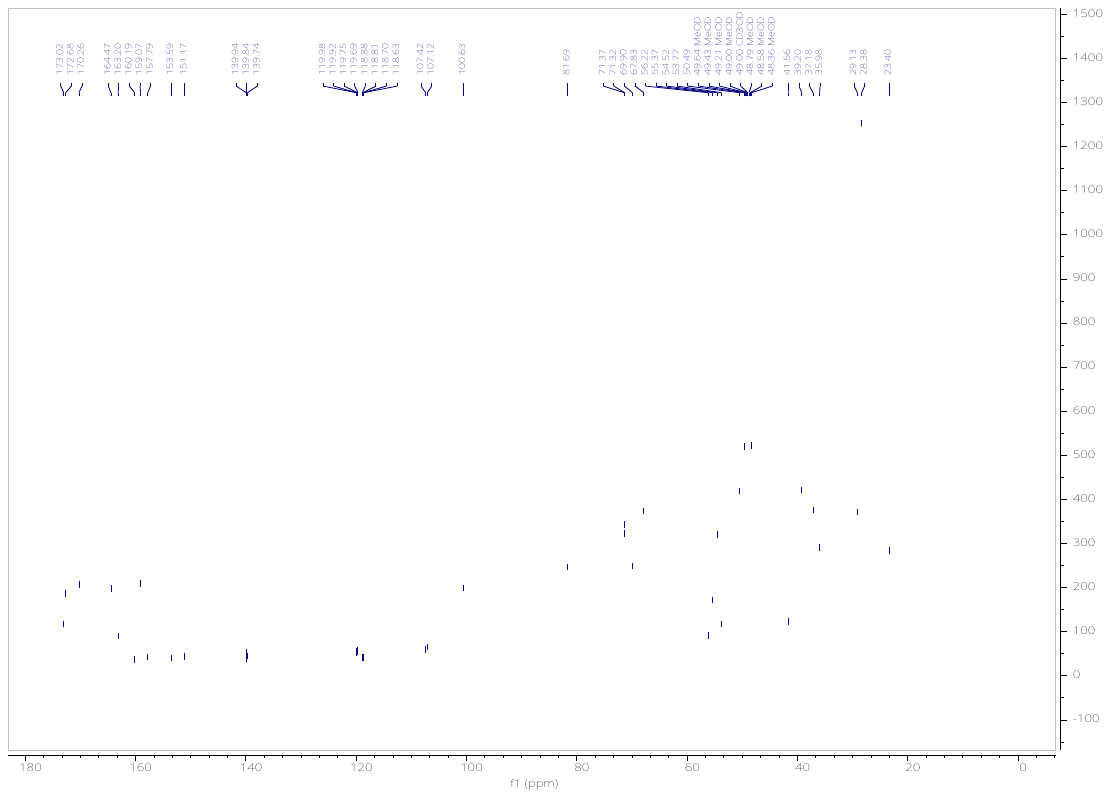

*tert*-Butyl 3-(6-(4-(4-((4,4-dimethylpiperidin-1-yl)methyl)-2,5-difluorophenyl)-2-oxo-1,4,9-triazaspiro[5.5]undecan-9-yl)pyrimidin-4-yl)-2-oxo-6,9,12-trioxa-3-azapentadecan-15-oate (**23**).

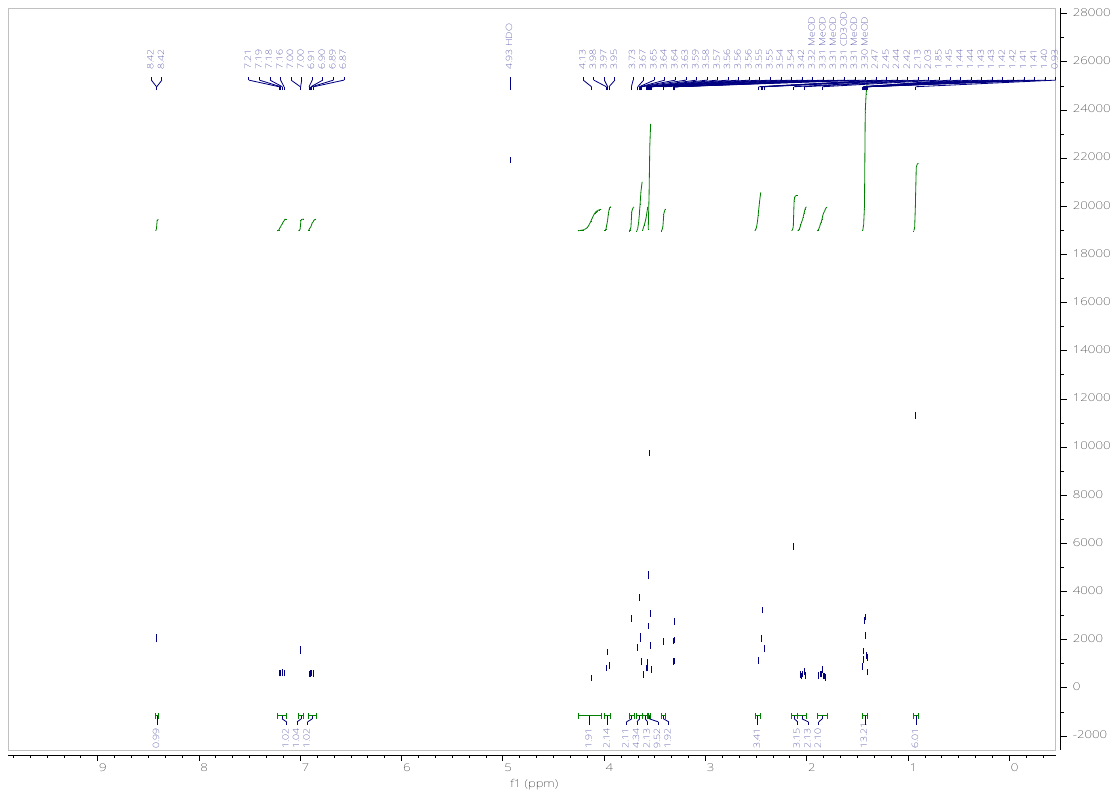

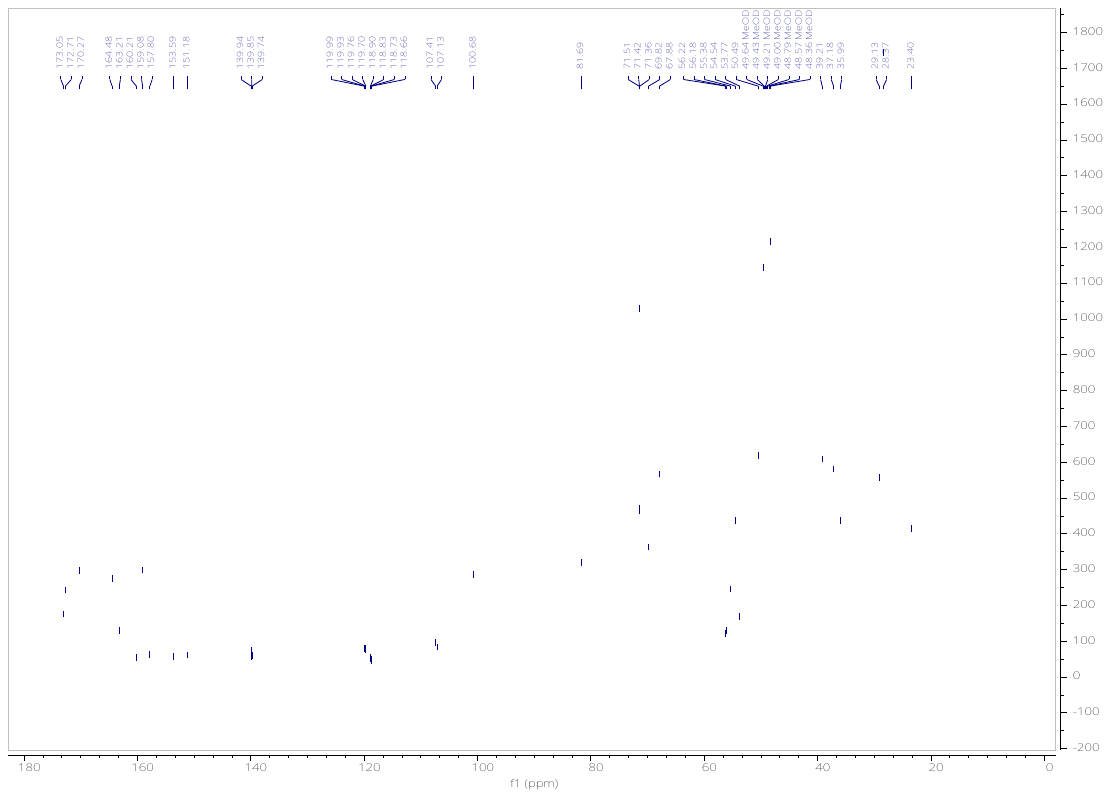

**^1^H and ^13^C NMR spectra of METTL3 PROTACs.**

(2*S*,4*R*)-1-((*S*)-2-(6-((6-(4-(4-((4,4-Dimethylpiperidin-1-yl)methyl)-2,5-difluorophenyl)-2-oxo-1,4,9-triazaspiro[5.5]undecan-9-yl)pyrimidin-4-yl)amino)hexanamido)-3,3-dimethylbutanoyl)-4-hydroxy-N-(4-(4-methylthiazol-5-yl)benzyl)pyrrolidine-2-carboxamide (**KH11**).

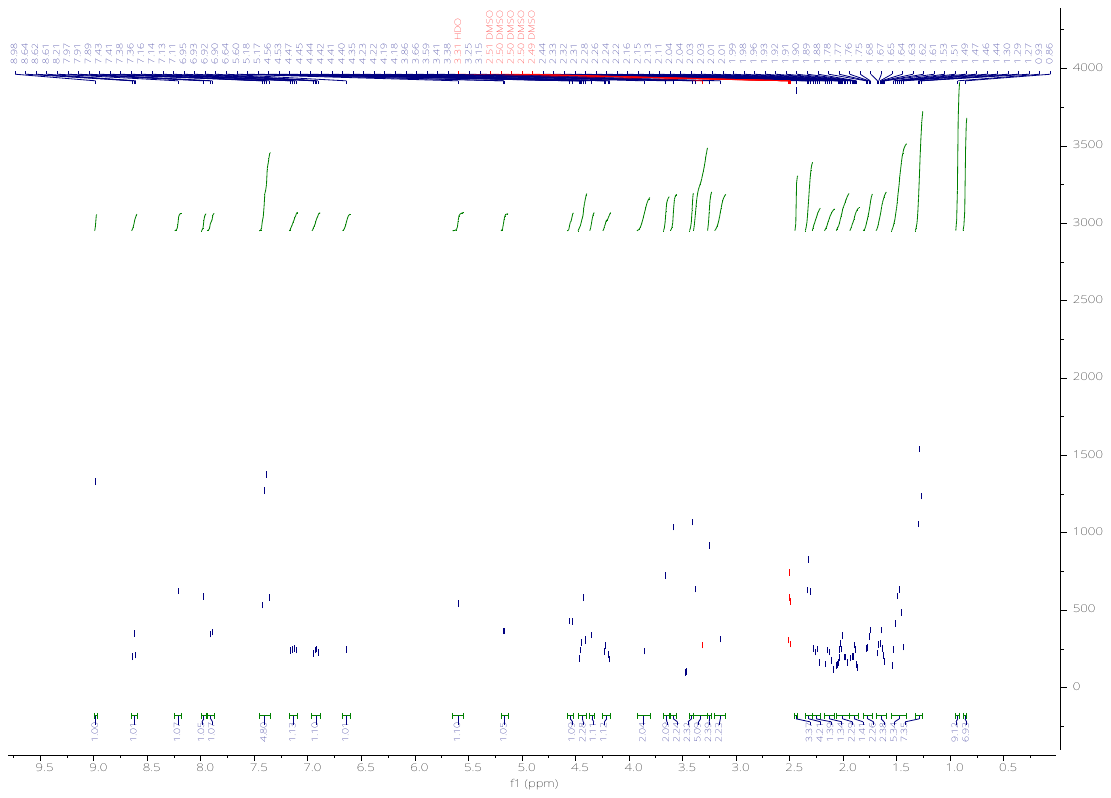

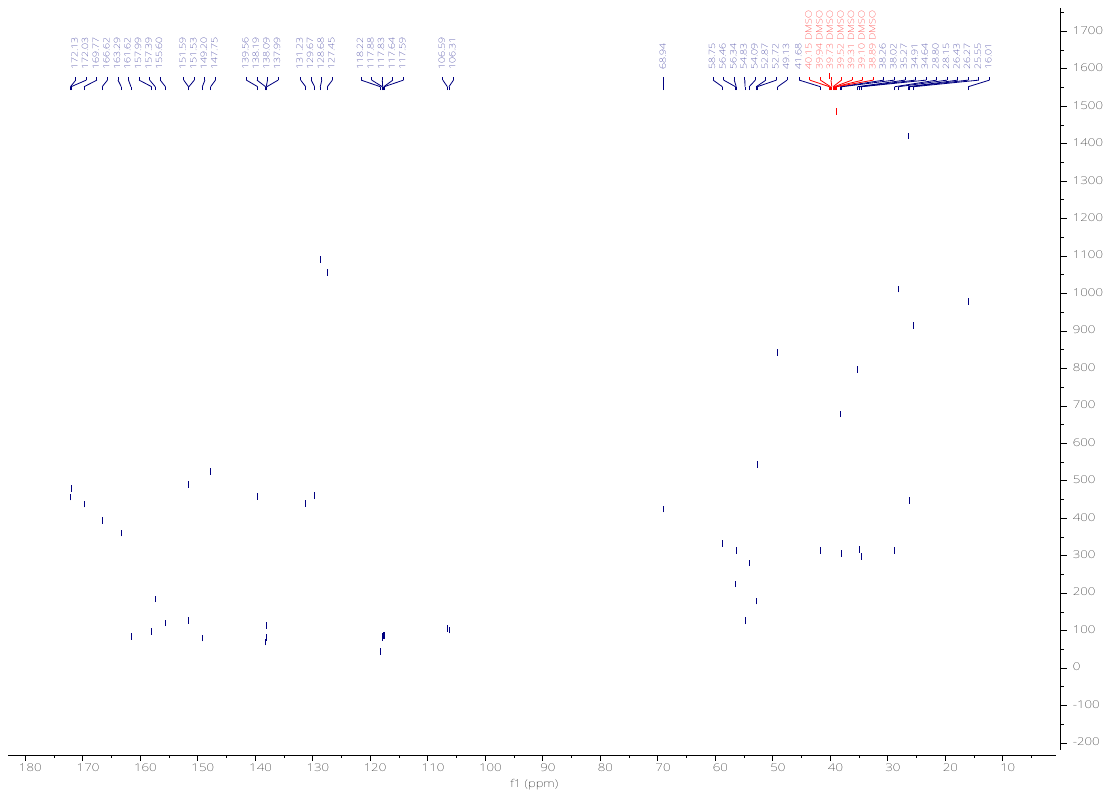

(2*S*,4*R*)-1-((*S*)-2-(8-((6-(4-(4-((4,4-Dimethylpiperidin-1-yl)methyl)-2,5-difluorophenyl)-2-oxo-1,4,9-triazaspiro[5.5]undecan-9-yl)pyrimidin-4-yl)amino)octanamido)-3,3-dimethylbutanoyl)-4-hydroxy-N-(4-(4-methylthiazol-5-yl)benzyl)pyrrolidine-2-carboxamide (**KH12**).

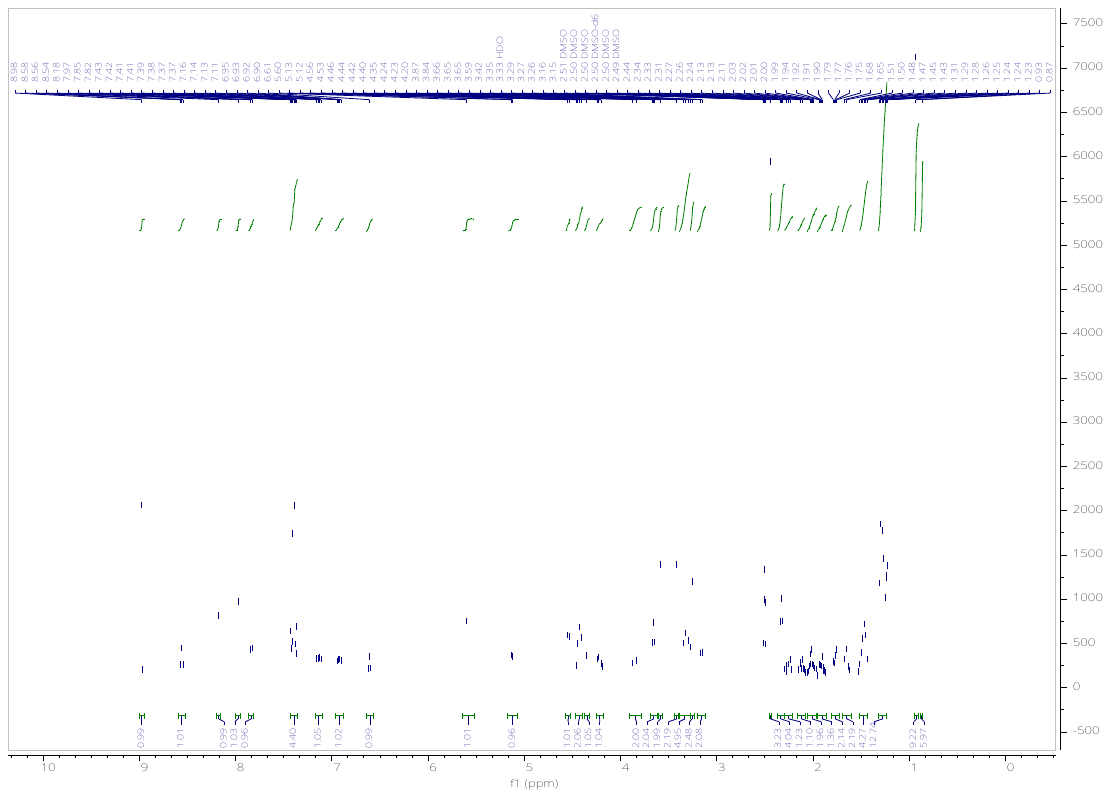

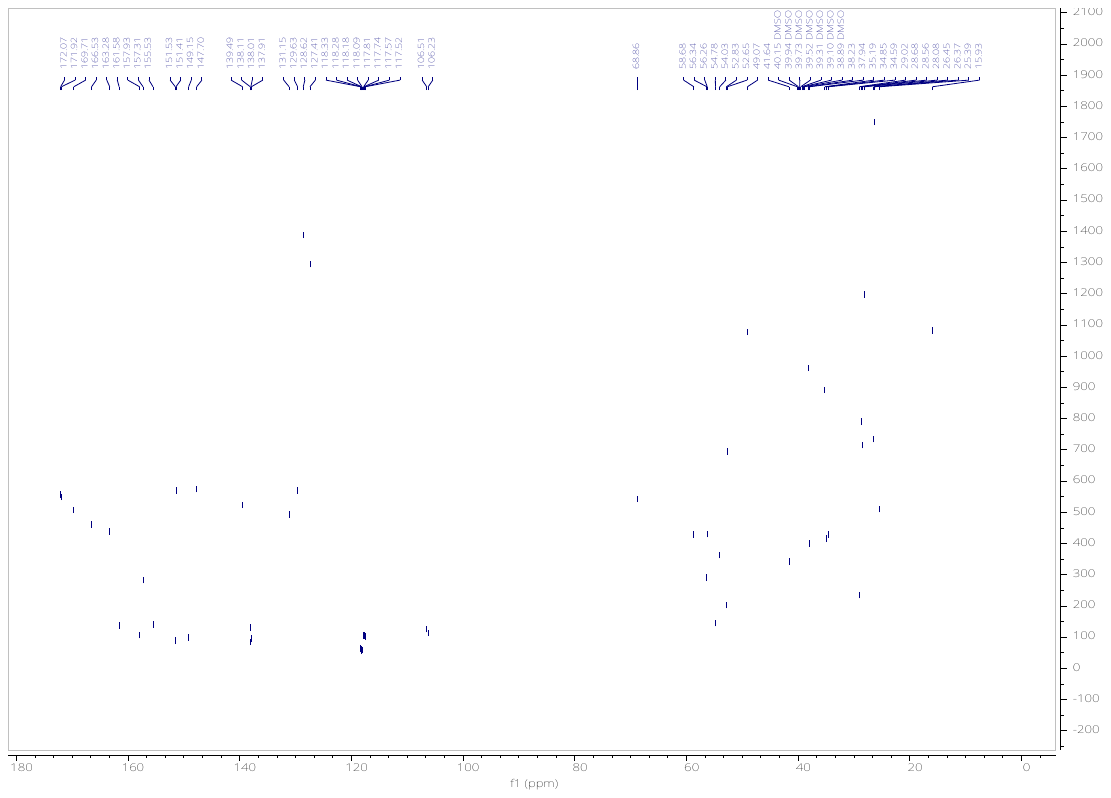

(2*S*,4*S*)-1-((*S*)-2-(8-((6-(4-(4-((4,4-dimethylpiperidin-1-yl)methyl)-2,5-difluorophenyl)-2-oxo-1,4,9-triazaspiro[5.5]undecan-9-yl)pyrimidin-4-yl)amino)octanamido)-3,3-dimethylbutanoyl)-4-hydroxy-N-(4-(4-methylthiazol-5-yl)benzyl)pyrrolidine-2-carboxamide (**KH12-NC**).

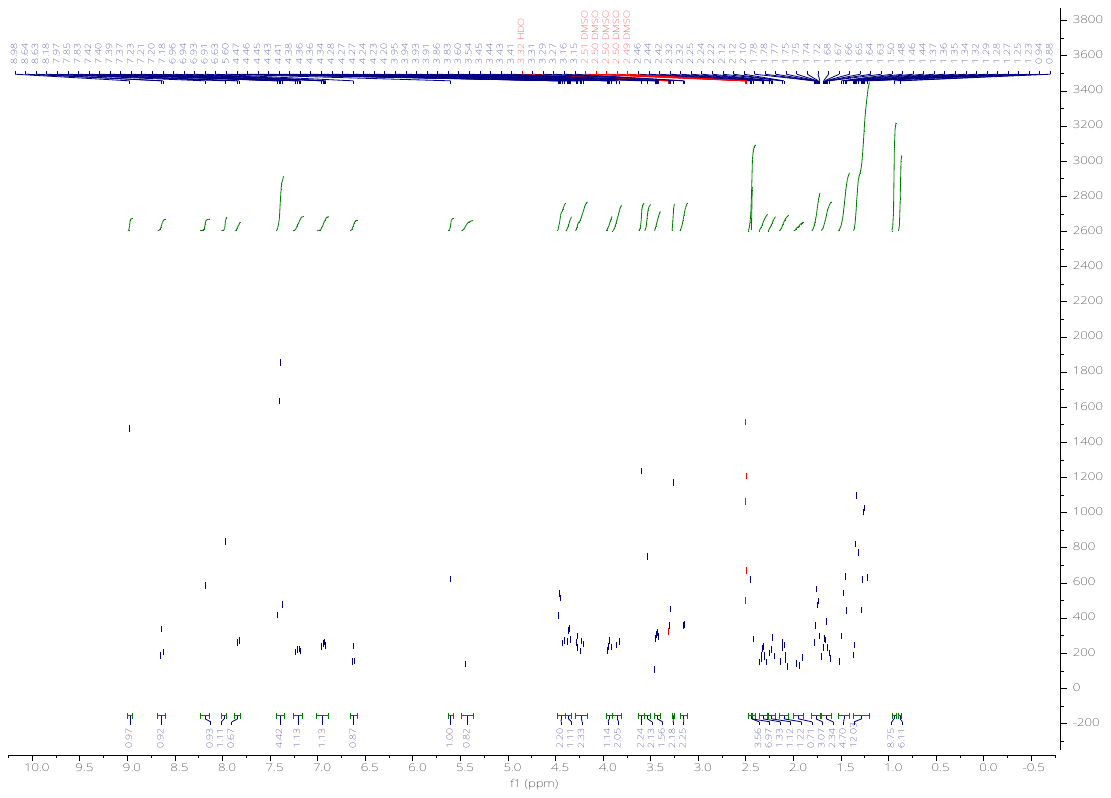

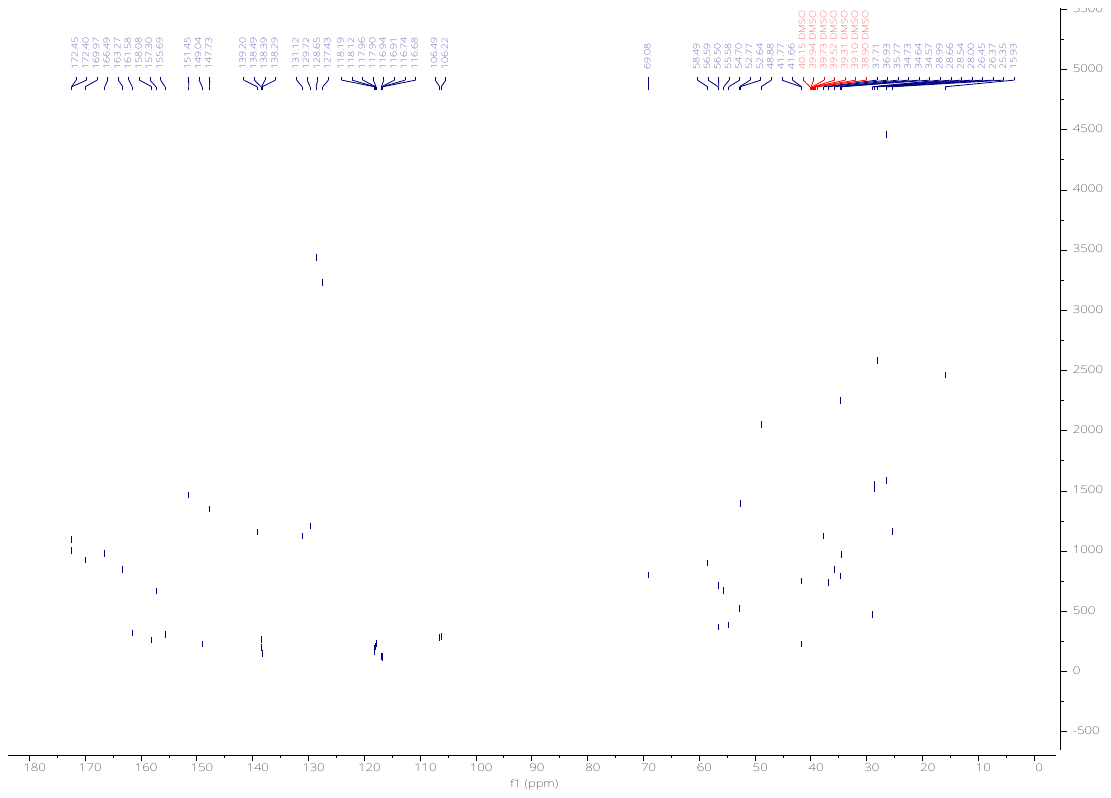

(2*S*,4*R*)-1-((*S*)-2-(11-((6-(4-(4-((4,4-dimethylpiperidin-1-yl)methyl)-2,5-difluorophenyl)-2-oxo-1,4,9-triazaspiro[5.5]undecan-9-yl)pyrimidin-4-yl)amino)undecanamido)-3,3-dimethylbutanoyl)-4-hydroxy-N-(4-(4-methylthiazol-5-yl)benzyl)pyrrolidine-2-carboxamide (**KH13**).

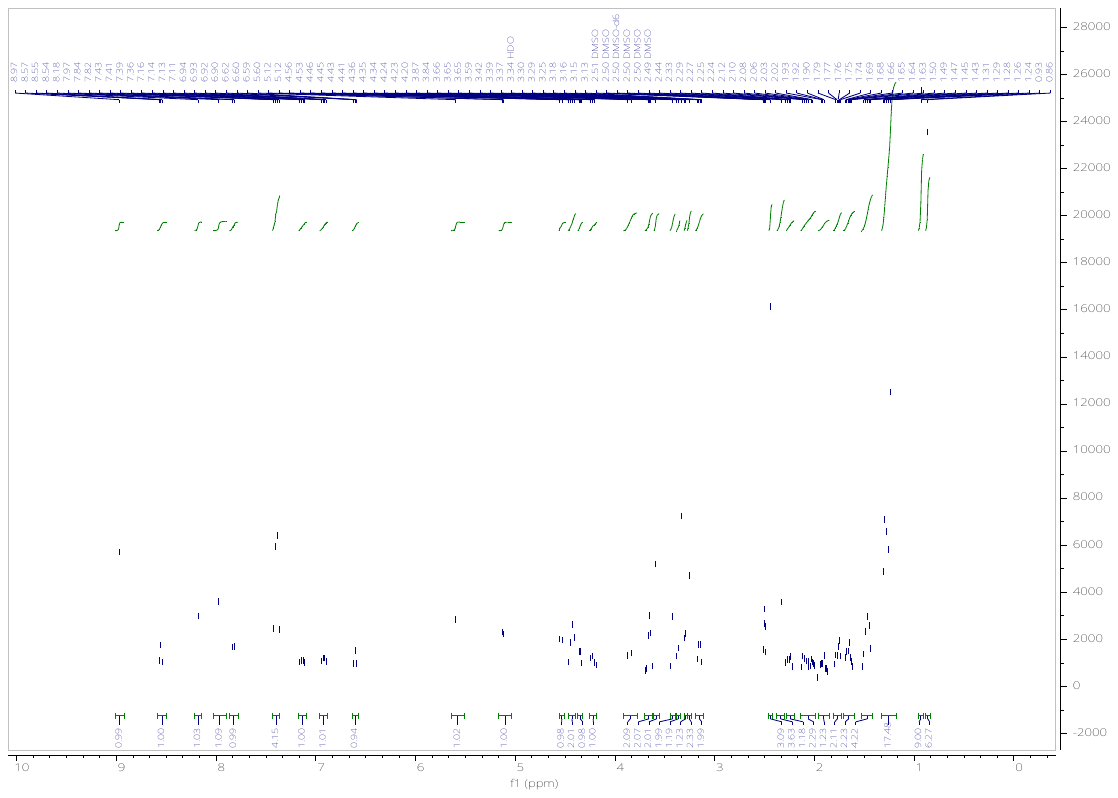

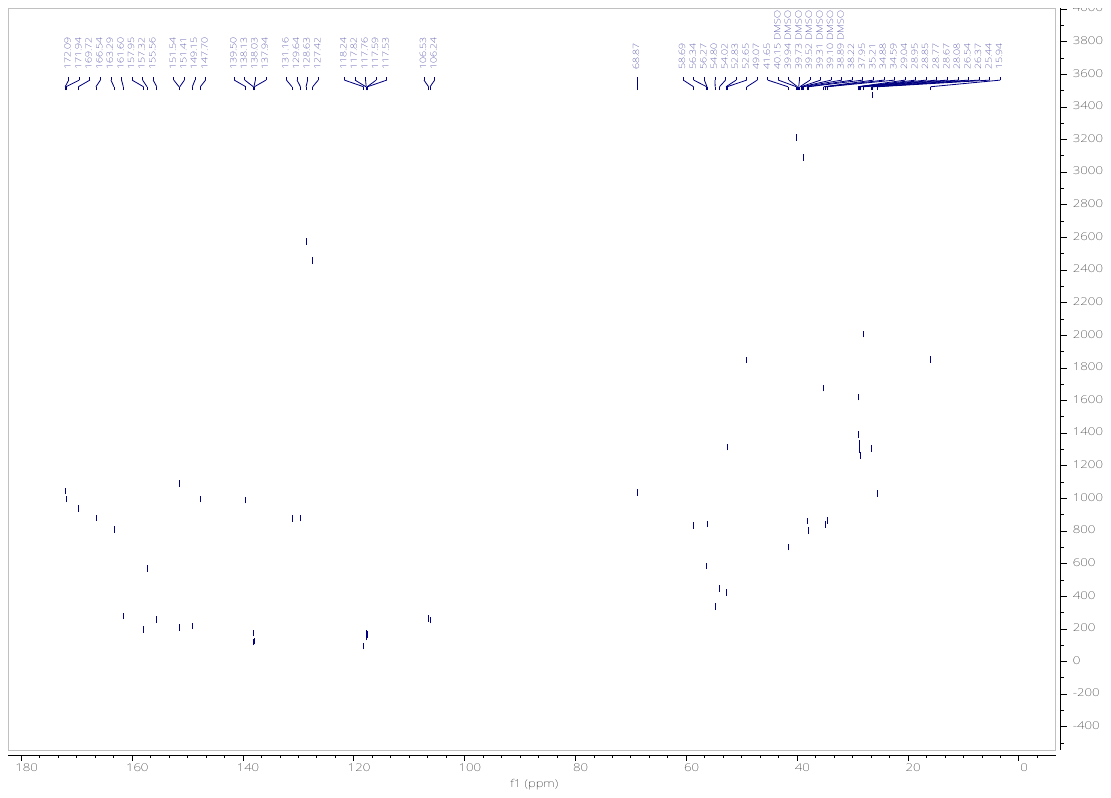

(2*S*,4*R*)-1-((*S*)-2-(3-(2-((6-(4-(4-((4,4-dimethylpiperidin-1-yl)methyl)-2,5-difluorophenyl)-2-oxo-1,4,9-triazaspiro[5.5]undecan-9-yl)pyrimidin-4-yl)amino)ethoxy)propanamido)-3,3-dimethylbutanoyl)-4-hydroxy-N-(4-(4-methylthiazol-5-yl)benzyl)pyrrolidine-2-carboxamide (**KH14**).

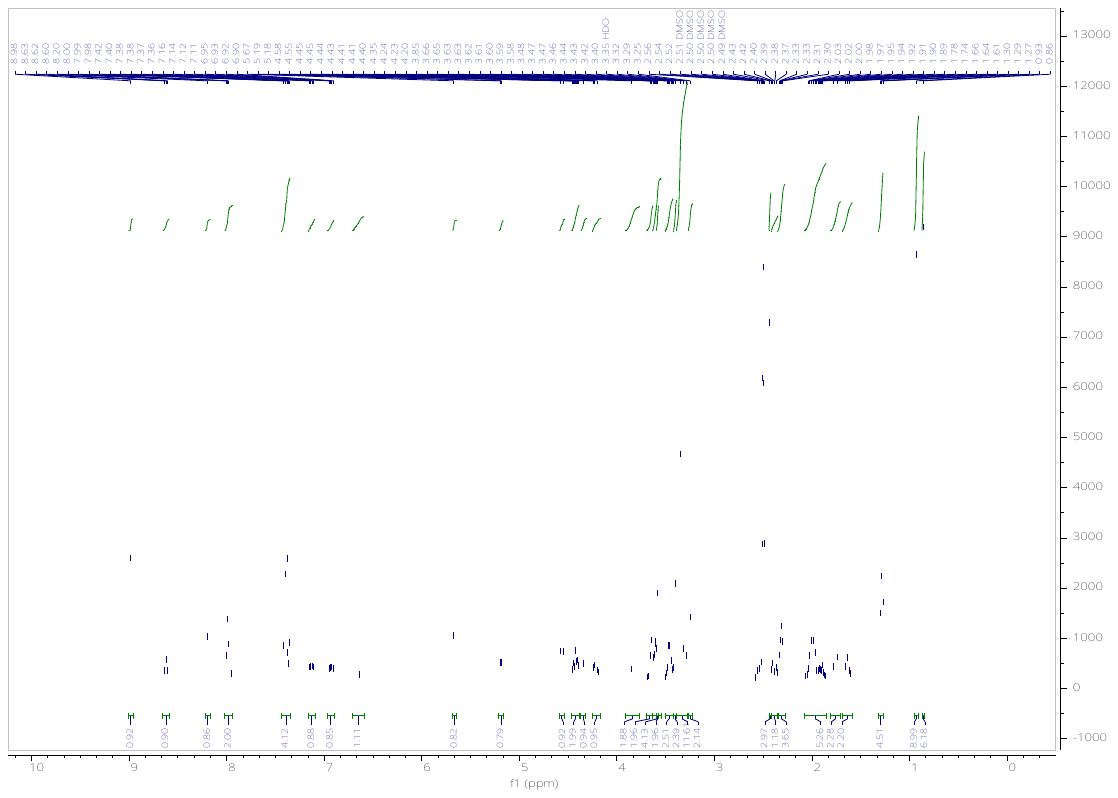

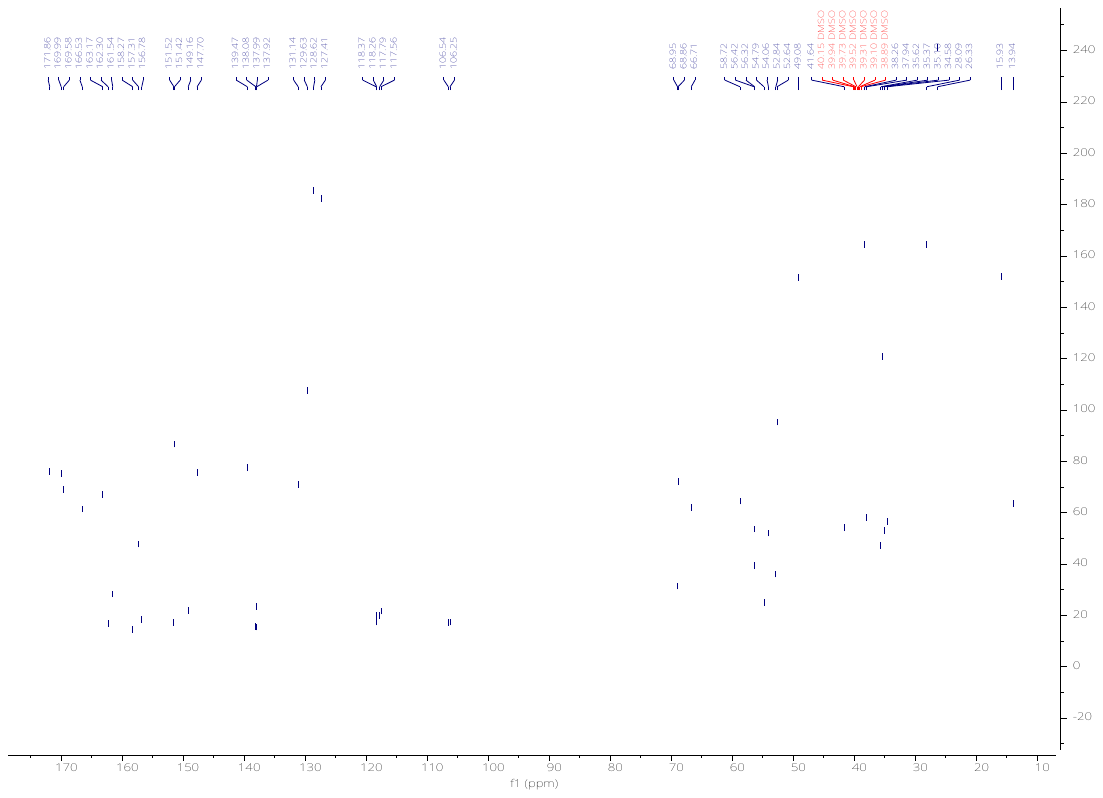

(2*S*,4*R*)-1-((*S*)-2-(3-(2-(2-((6-(4-(4-((4,4-dimethylpiperidin-1-yl)methyl)-2,5-difluorophenyl)-2-oxo-1,4,9-triazaspiro[5.5]undecan-9-yl)pyrimidin-4-yl)amino)ethoxy)ethoxy)propanamido)-3,3-dimethylbutanoyl)-4-hydroxy-N-(4-(4-methylthiazol-5-yl)benzyl)pyrrolidine-2-carboxamide (**KH15**).

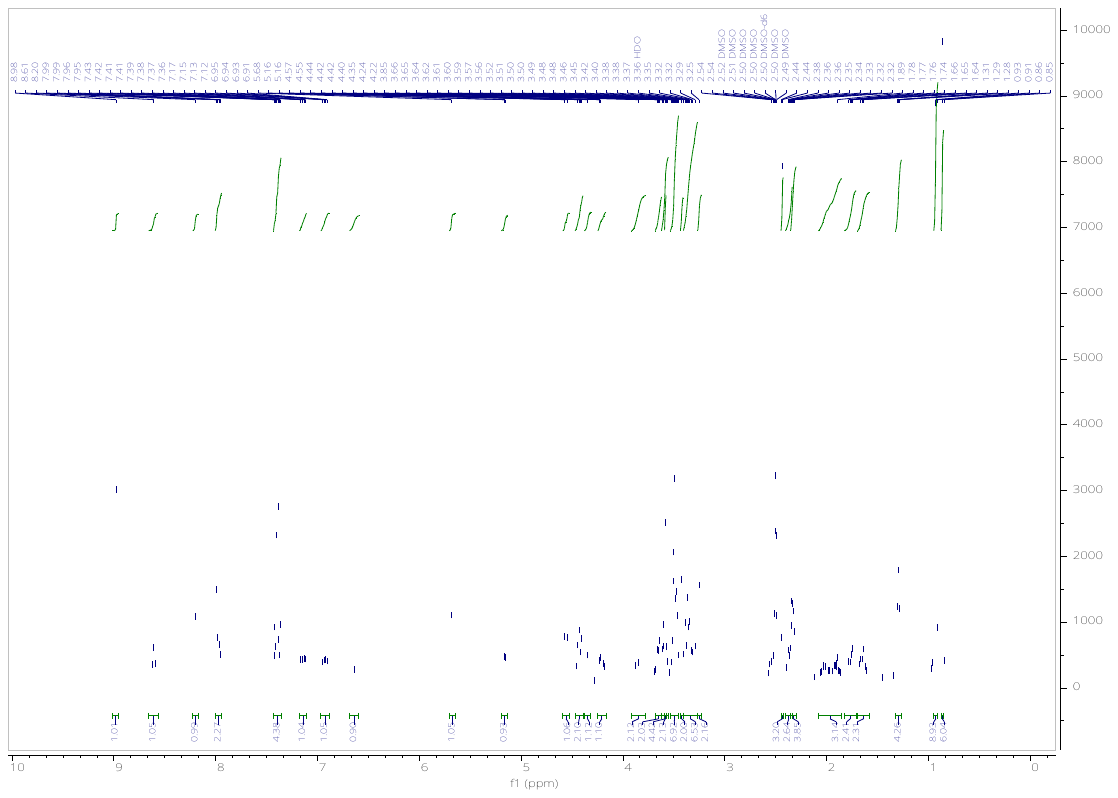

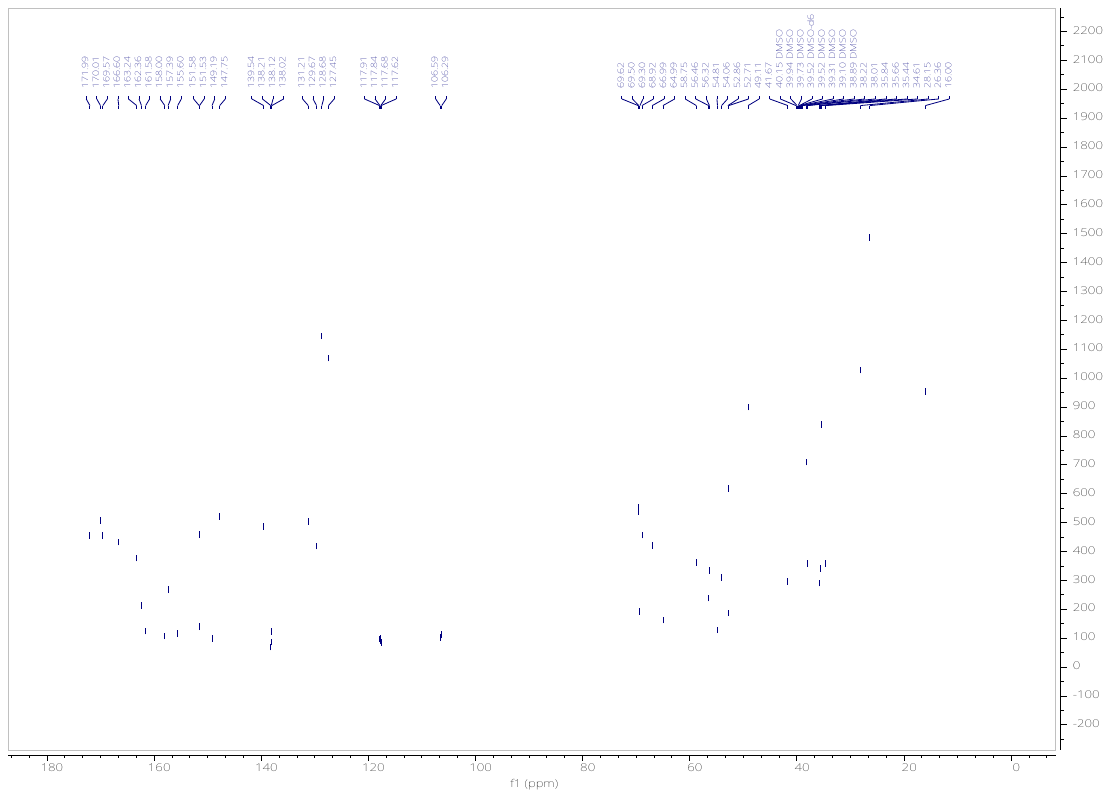

(2*S*,4*R*)-1-((*S*)-14-(tert-butyl)-1-((6-(4-(4-((4,4-dimethylpiperidin-1-yl)methyl)-2,5-difluorophenyl)-2-oxo-1,4,9-triazaspiro[5.5]undecan-9-yl)pyrimidin-4-yl)amino)-12-oxo-3,6,9-trioxa-13-azapentadecan-15-oyl)-4-hydroxy-N-(4-(4-methylthiazol-5-yl)benzyl)pyrrolidine-2-carboxamide (**KH16**).

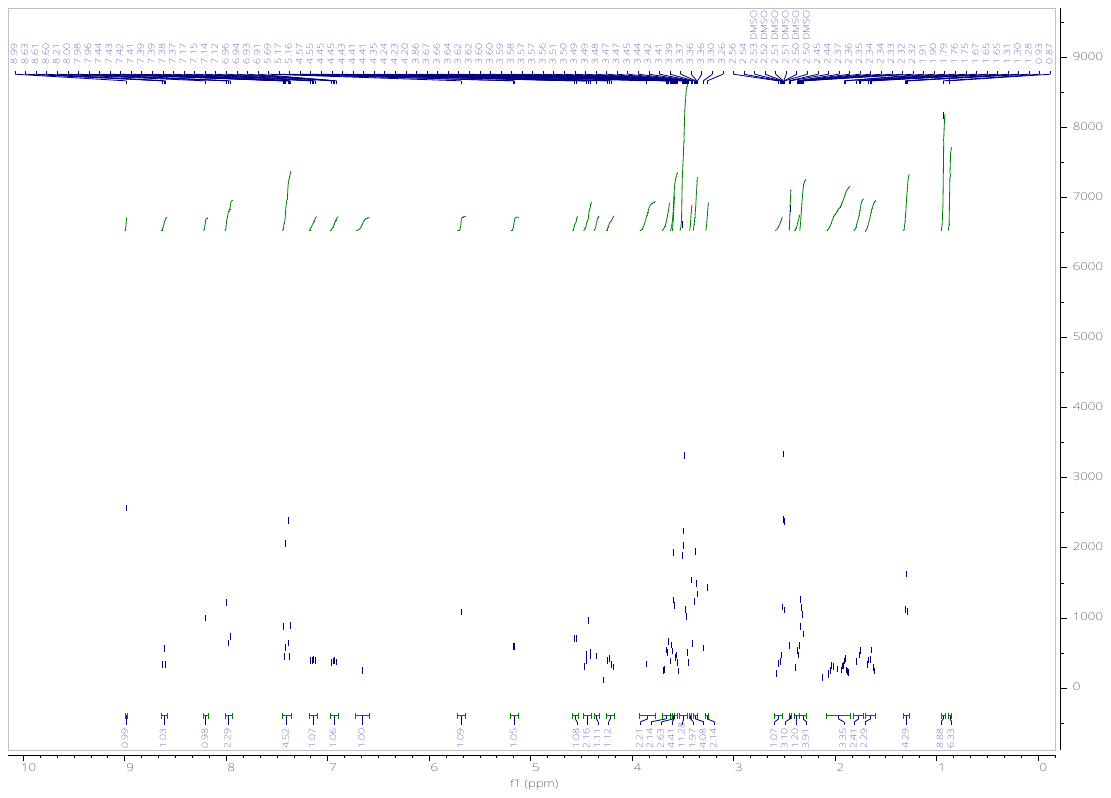

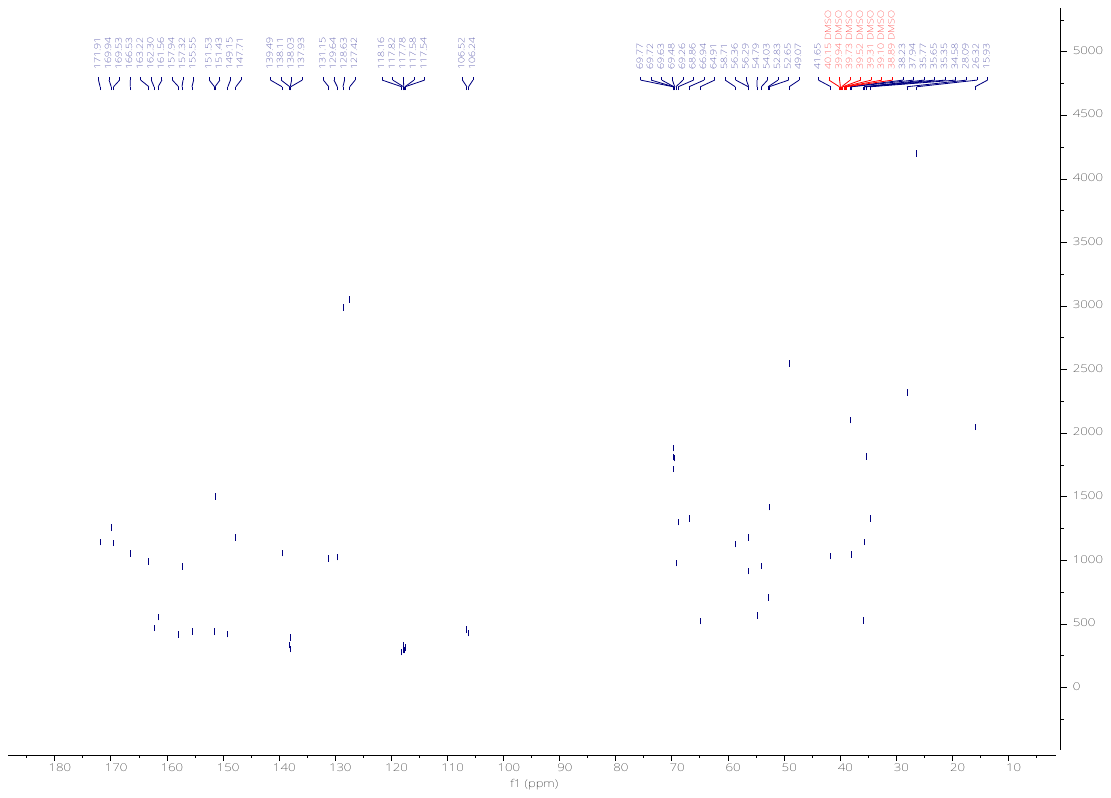

5-(4-(6-((6-(4-(4-((4,4-dimethylpiperidin-1-yl)methyl)-2,5-difluorophenyl)-2-oxo-1,4,9-triazaspiro[5.5]undecan-9-yl)pyrimidin-4-yl)amino)hexanoyl)piperazin-1-yl)-2-(2,6-dioxopiperidin-3-yl)isoindoline-1,3-dione (**KH21**).

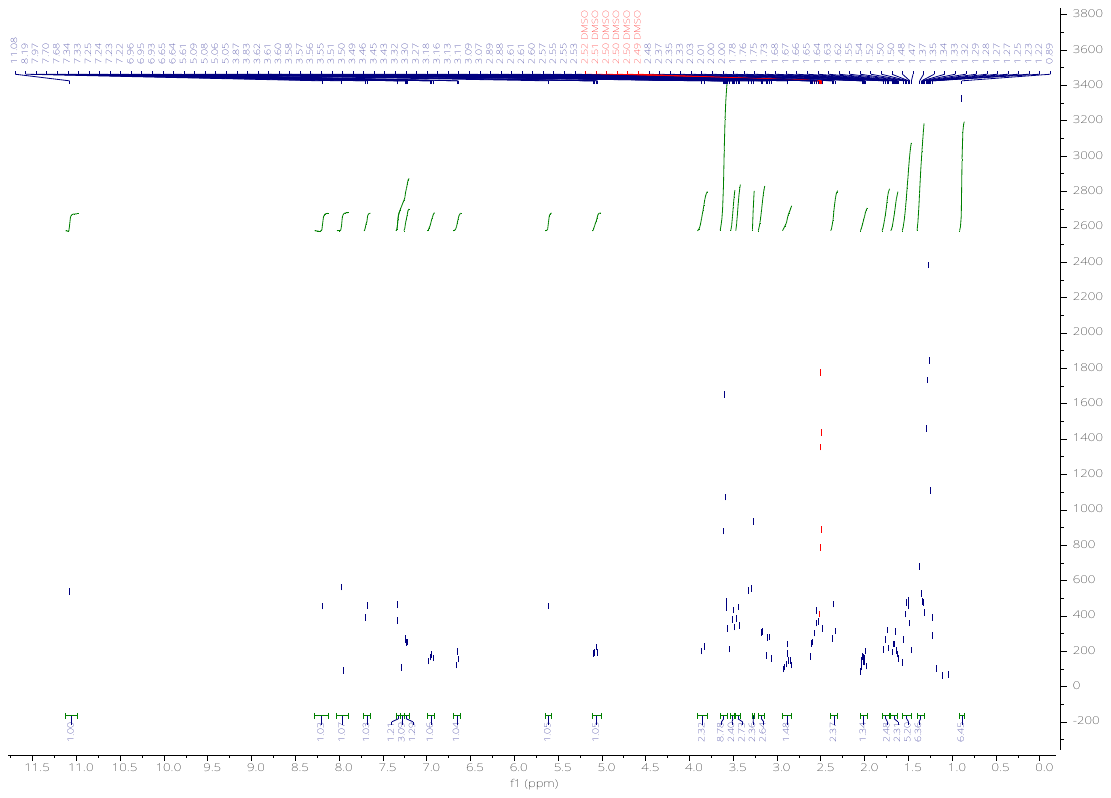

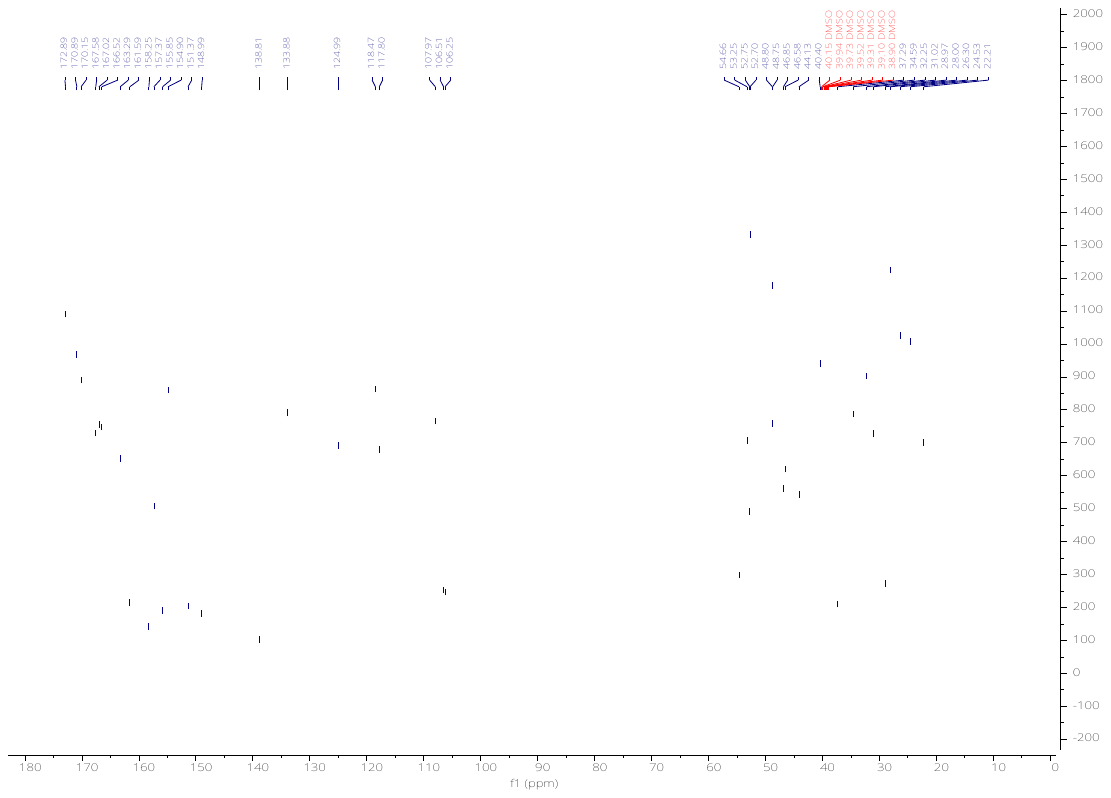

5-(4-(8-((6-(4-(4-((4,4-dimethylpiperidin-1-yl)methyl)-2,5-difluorophenyl)-2-oxo-1,4,9-triazaspiro[5.5]undecan-9-yl)pyrimidin-4-yl)amino)octanoyl)piperazin-1-yl)-2-(2,6-dioxopiperidin-3-yl)isoindoline-1,3-dione (**KH22**).

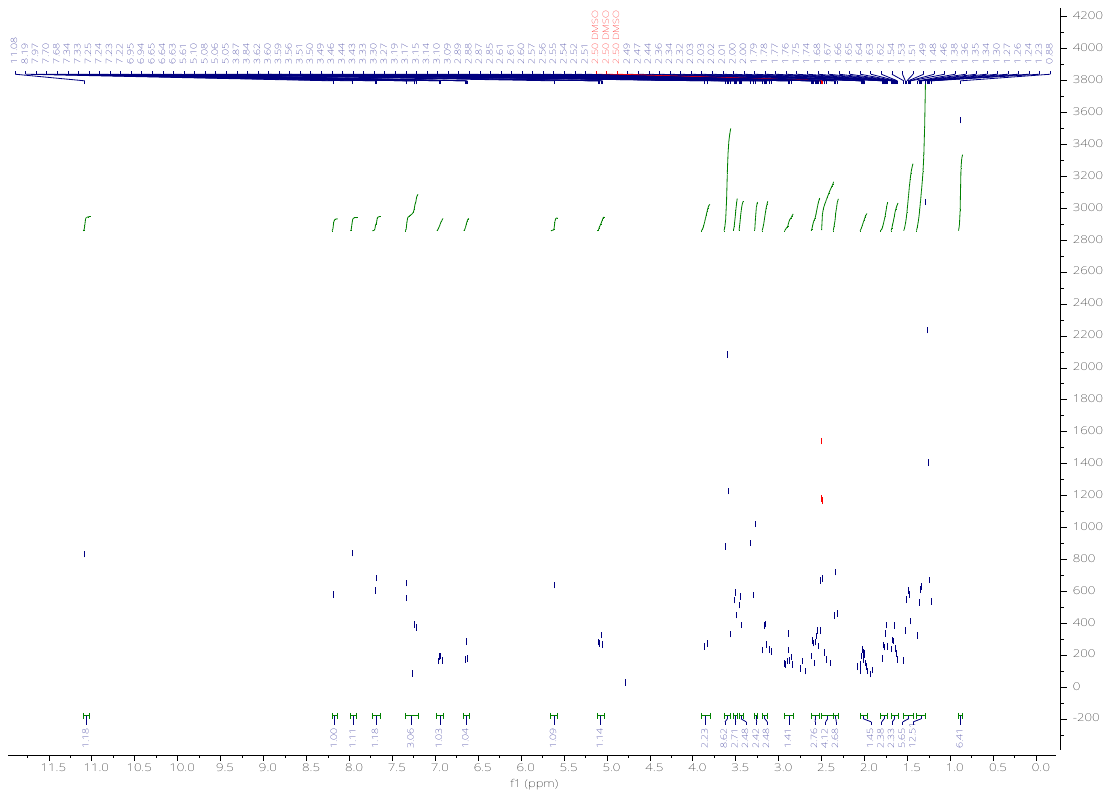

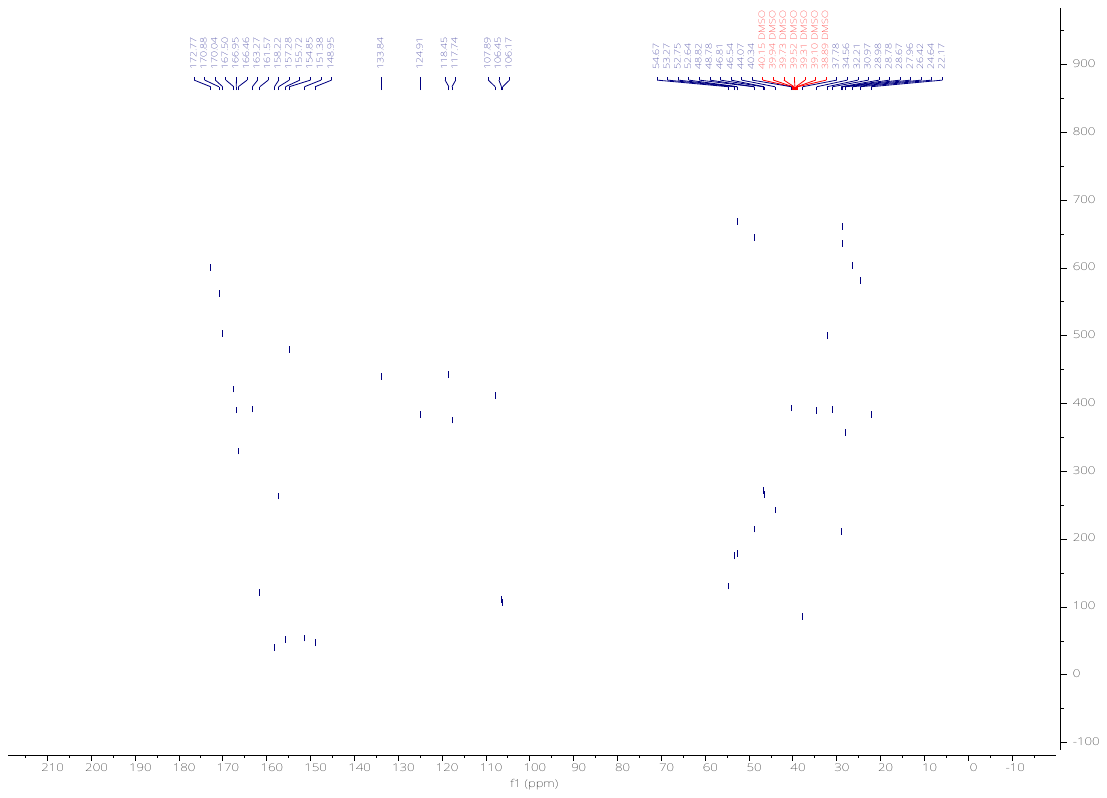

5-(4-(11-((6-(4-(4-((4,4-dimethylpiperidin-1-yl)methyl)-2,5-difluorophenyl)-2-oxo-1,4,9-triazaspiro[5.5]undecan-9-yl)pyrimidin-4-yl)amino)undecanoyl)piperazin-1-yl)-2-(2,6-dioxopiperidin-3-yl)isoindoline-1,3-dione (**KH23**).

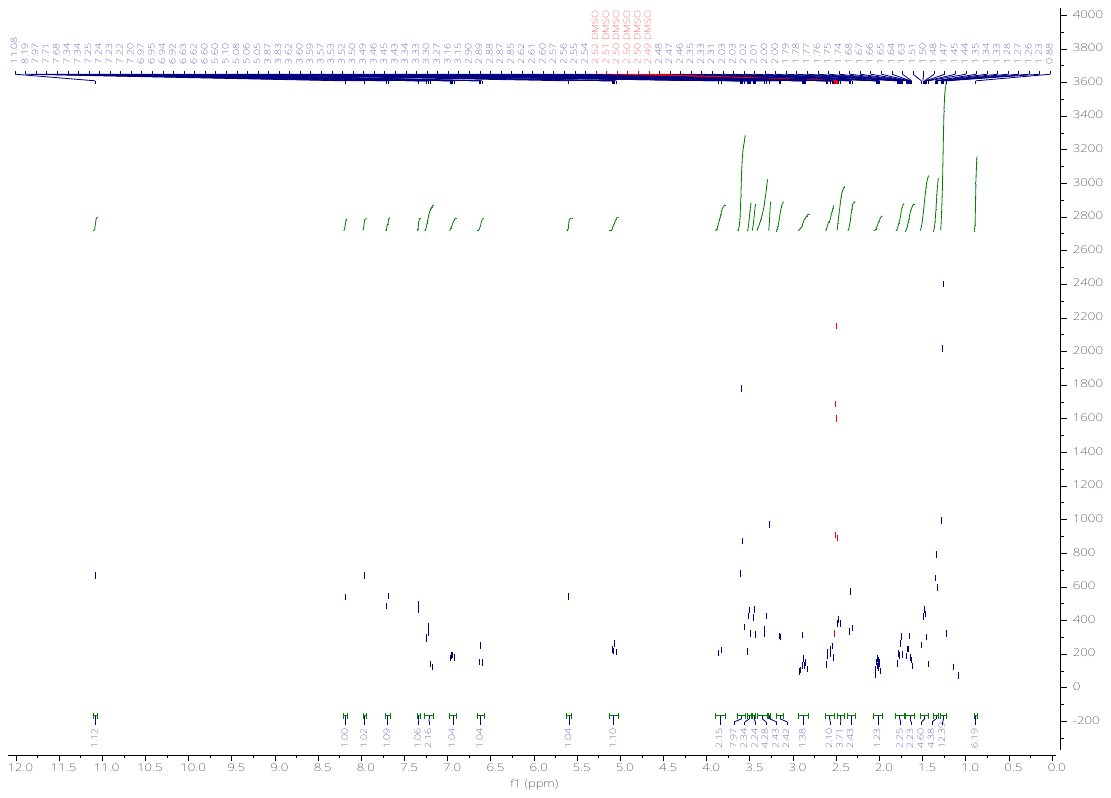

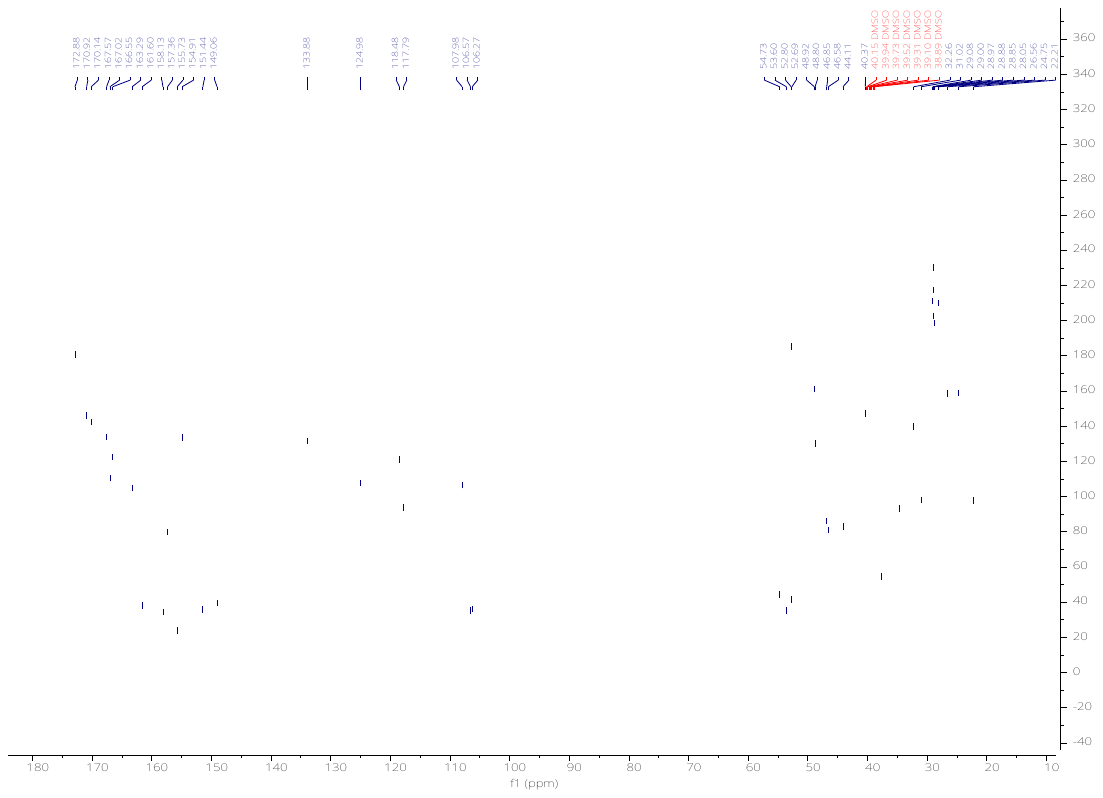

5-(4-(3-(2-((6-(4-(4-((4,4-dimethylpiperidin-1-yl)methyl)-2,5-difluorophenyl)-2-oxo-1,4,9-triazaspiro[5.5]undecan-9-yl)pyrimidin-4-yl)amino)ethoxy)propanoyl)piperazin-1-yl)-2-(2,6-dioxopiperidin-3-yl)isoindoline-1,3-dione (**KH24**).

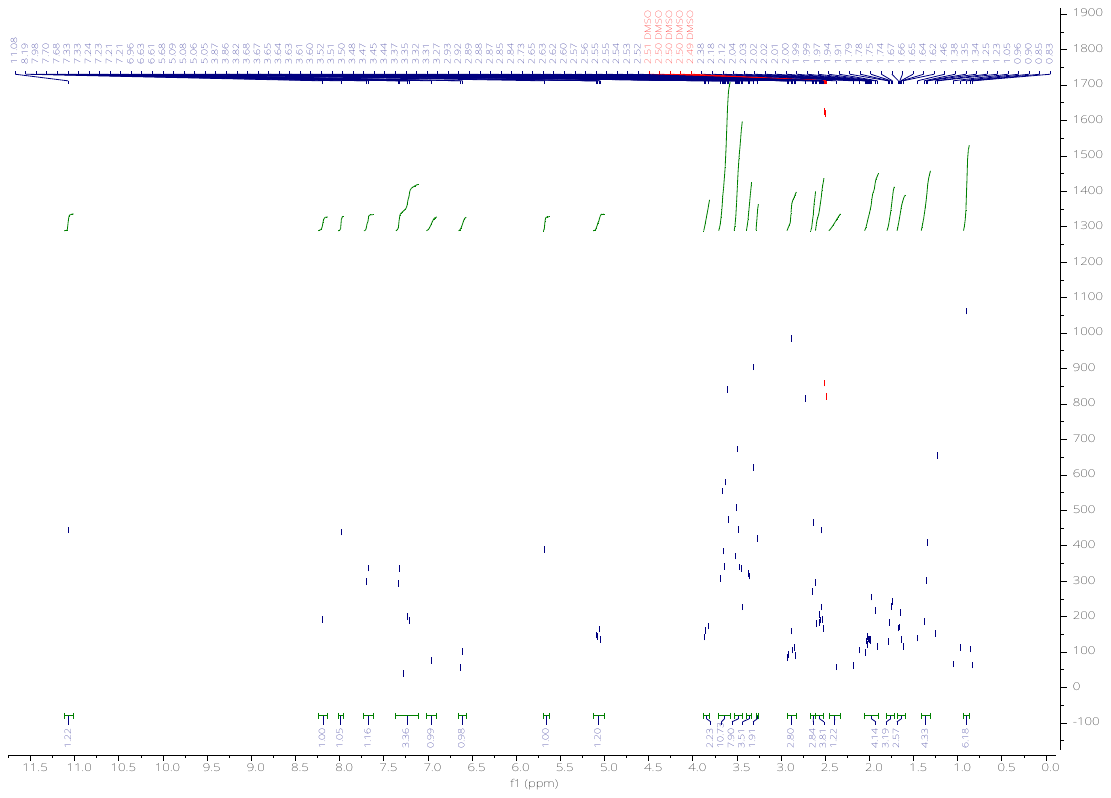

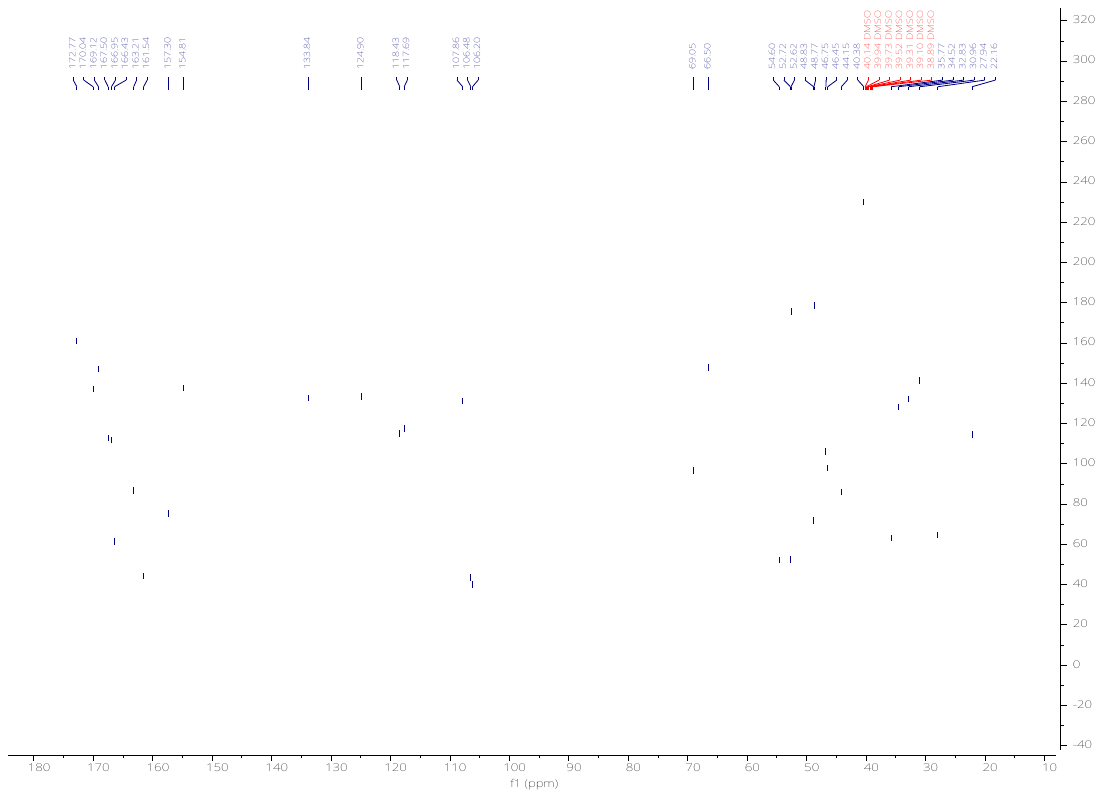

5-(4-(3-(2-(2-((6-(4-(4-((4,4-dimethylpiperidin-1-yl)methyl)-2,5-difluorophenyl)-2-oxo-1,4,9-triazaspiro[5.5]undecan-9-yl)pyrimidin-4-yl)amino)ethoxy)ethoxy)propanoyl)piperazin-1-yl)-2-(2,6-dioxopiperidin-3-yl)isoindoline-1,3-dione (**KH25**).

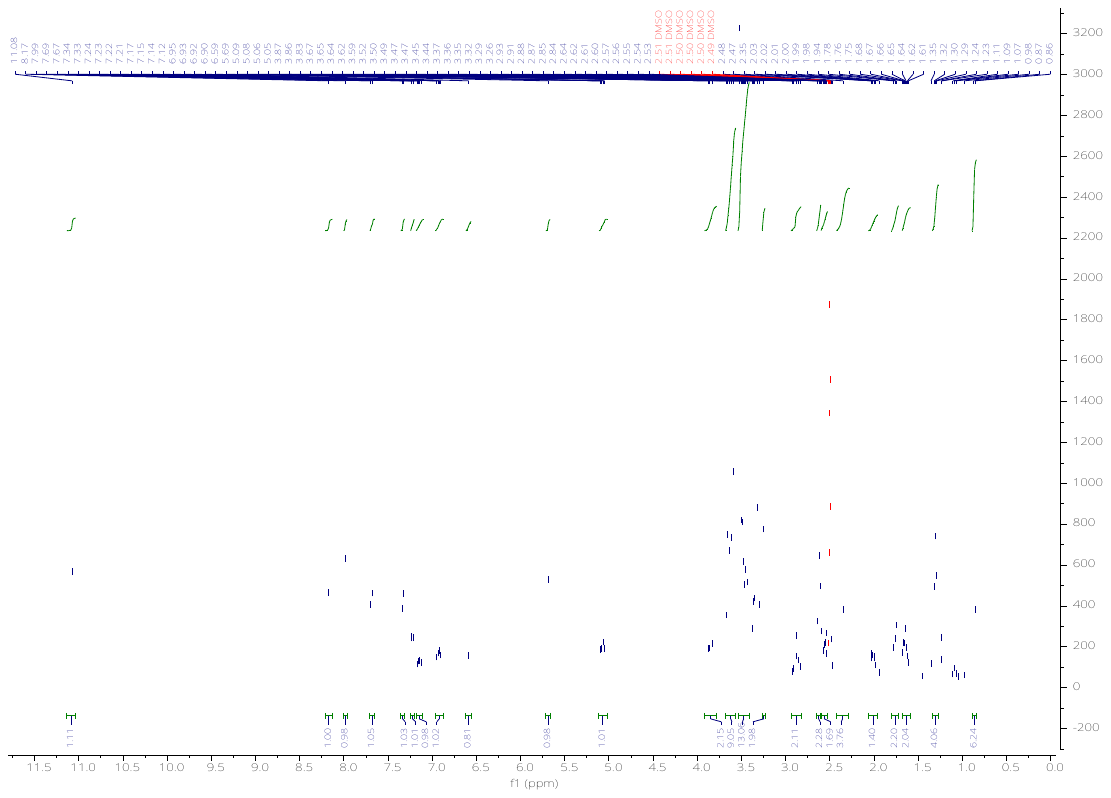

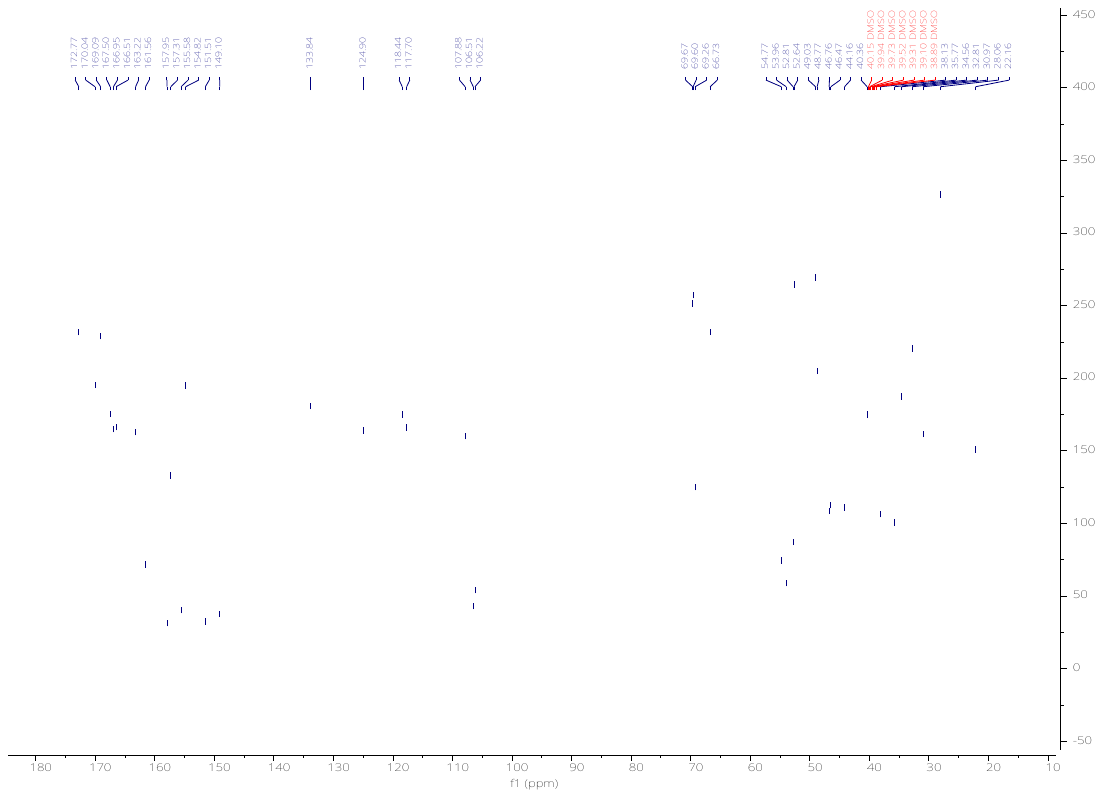

5-(4-(3-(2-(2-(2-((6-(4-(4-((4,4-dimethylpiperidin-1-yl)methyl)-2,5-difluorophenyl)-2-oxo-1,4,9-triazaspiro[5.5]undecan-9-yl)pyrimidin-4-yl)amino)ethoxy)ethoxy)ethoxy)propanoyl)piperazin-1-yl)-2-(2,6-dioxopiperidin-3-yl)isoindoline-1,3-dione (**KH26**)

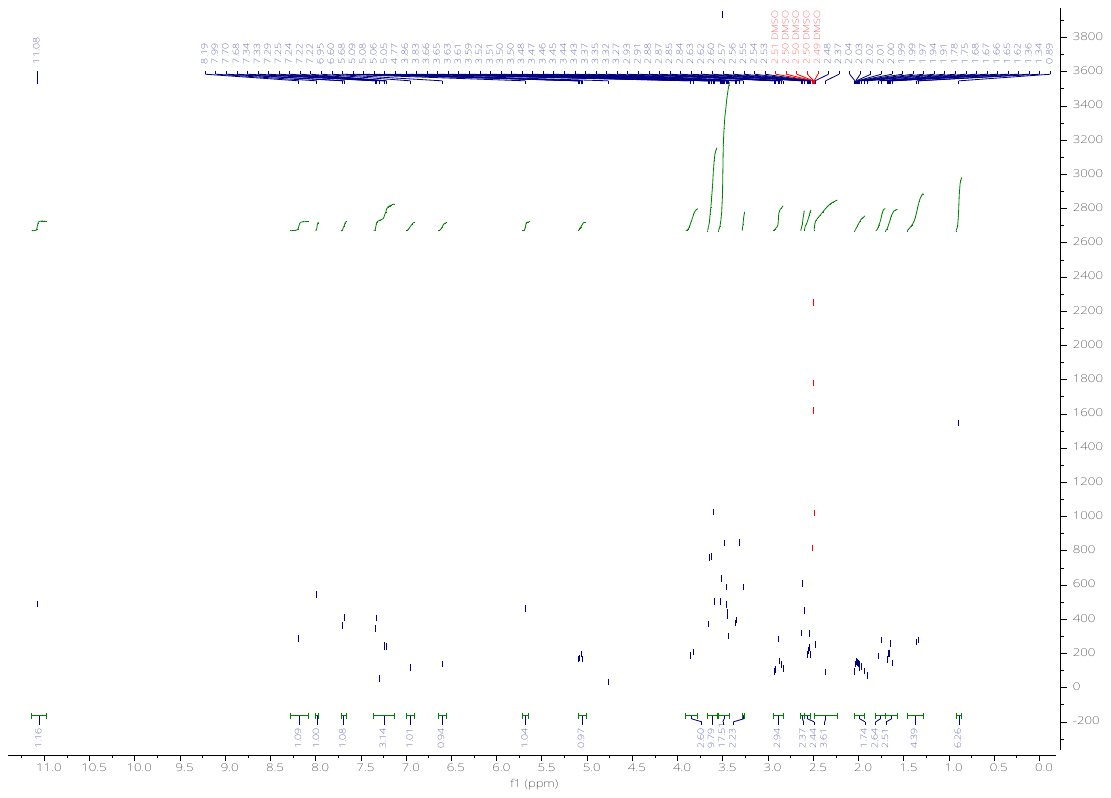

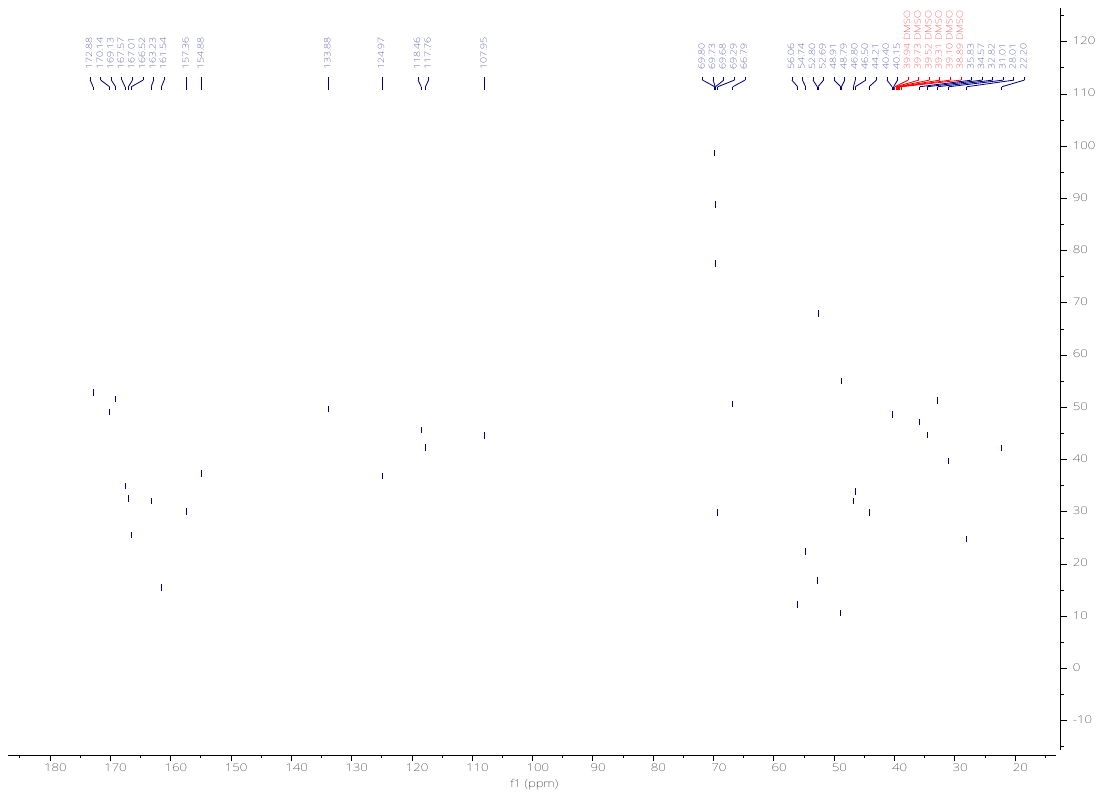

**HPLC traces of METTL3 PROTACs**

(2*S*,4*R*)-1-((*S*)-2-(6-((6-(4-(4-((4,4-Dimethylpiperidin-1-yl)methyl)-2,5-difluorophenyl)-2-oxo-1,4,9-triazaspiro[5.5]undecan-9-yl)pyrimidin-4-yl)amino)hexanamido)-3,3-dimethylbutanoyl)-4-hydroxy-N-(4-(4-methylthiazol-5-yl)benzyl)pyrrolidine-2-carboxamide (**KH11**).

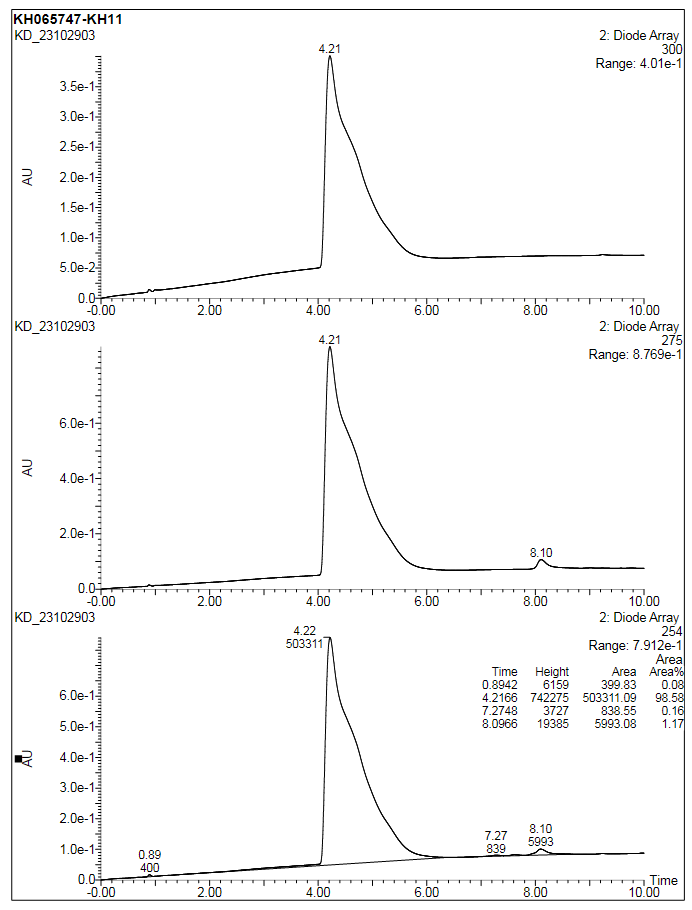

(2*S*,4*R*)-1-((*S*)-2-(8-((6-(4-(4-((4,4-Dimethylpiperidin-1-yl)methyl)-2,5-difluorophenyl)-2-oxo-1,4,9-triazaspiro[5.5]undecan-9-yl)pyrimidin-4-yl)amino)octanamido)-3,3-dimethylbutanoyl)-4-hydroxy-N-(4-(4-methylthiazol-5-yl)benzyl)pyrrolidine-2-carboxamide (**KH12**).

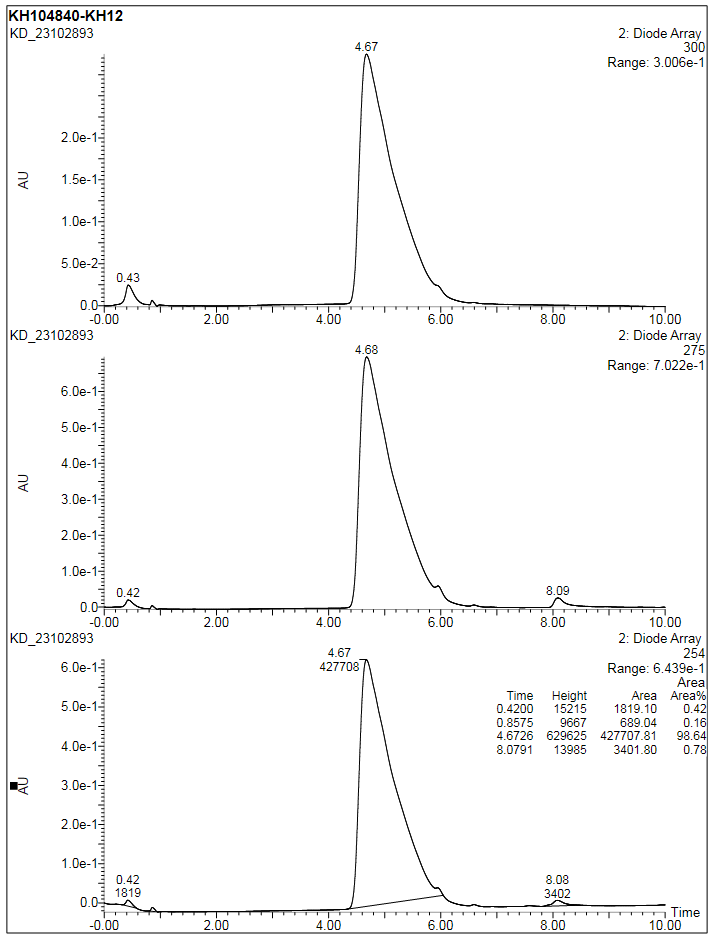

(2*S*,4*S*)-1-((*S*)-2-(8-((6-(4-(4-((4,4-dimethylpiperidin-1-yl)methyl)-2,5-difluorophenyl)-2-oxo-1,4,9-triazaspiro[5.5]undecan-9-yl)pyrimidin-4-yl)amino)octanamido)-3,3-dimethylbutanoyl)-4-hydroxy-N-(4-(4-methylthiazol-5-yl)benzyl)pyrrolidine-2-carboxamide (**KH12-NC**).

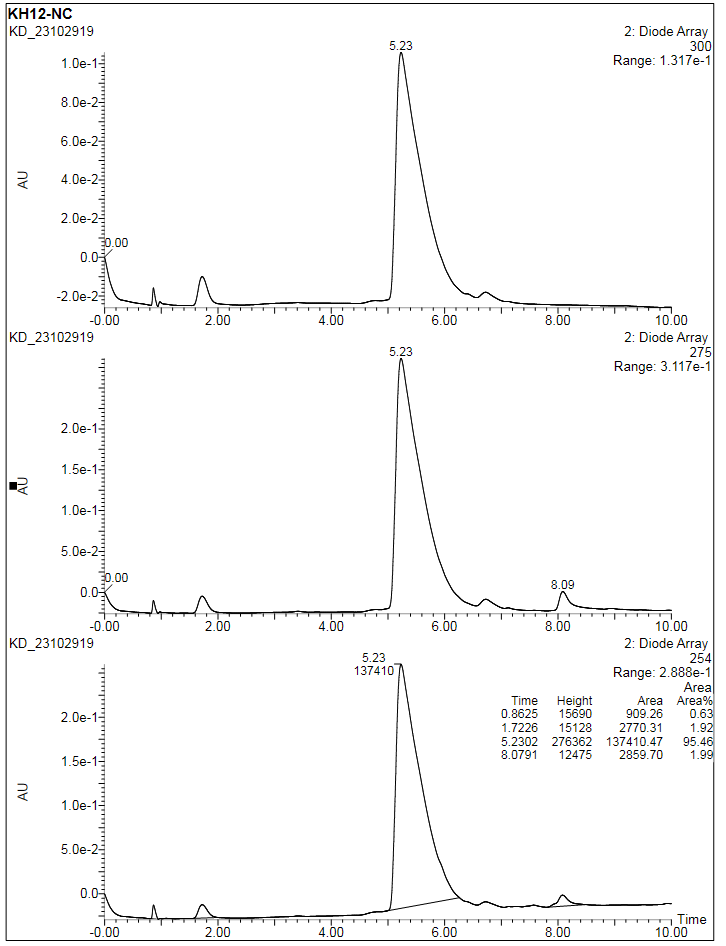

(2*S*,4*R*)-1-((*S*)-2-(11-((6-(4-(4-((4,4-dimethylpiperidin-1-yl)methyl)-2,5-difluorophenyl)-2-oxo-1,4,9-triazaspiro[5.5]undecan-9-yl)pyrimidin-4-yl)amino)undecanamido)-3,3-dimethylbutanoyl)-4-hydroxy-N-(4-(4-methylthiazol-5-yl)benzyl)pyrrolidine-2-carboxamide (**KH13**).

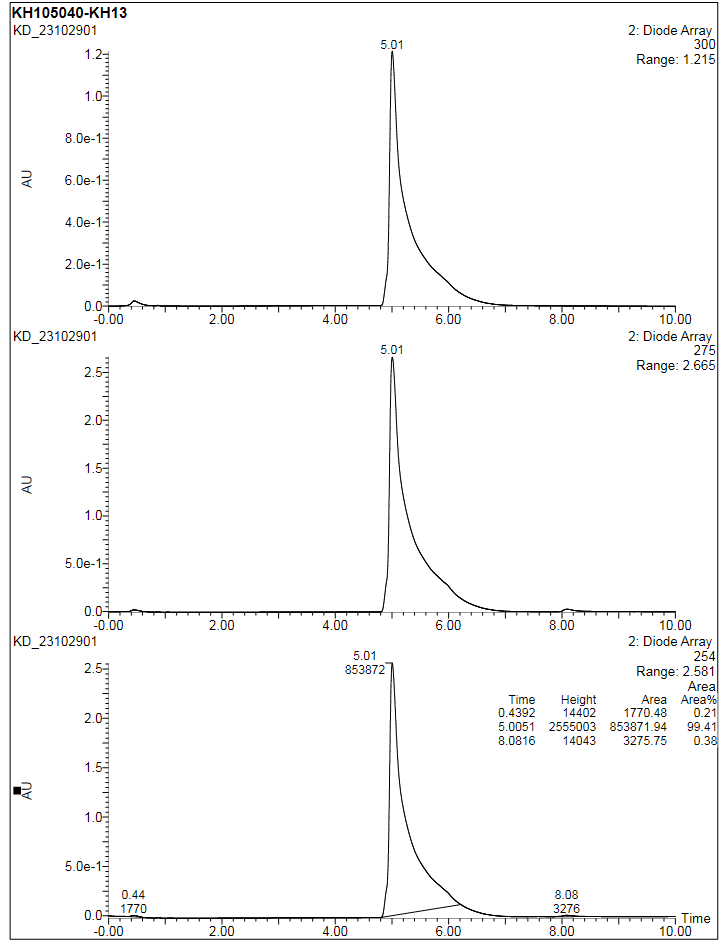

(2*S*,4*R*)-1-((*S*)-2-(3-(2-((6-(4-(4-((4,4-dimethylpiperidin-1-yl)methyl)-2,5-difluorophenyl)-2-oxo-1,4,9-triazaspiro[5.5]undecan-9-yl)pyrimidin-4-yl)amino)ethoxy)propanamido)-3,3-dimethylbutanoyl)-4-hydroxy-N-(4-(4-methylthiazol-5-yl)benzyl)pyrrolidine-2-carboxamide (**KH14**).

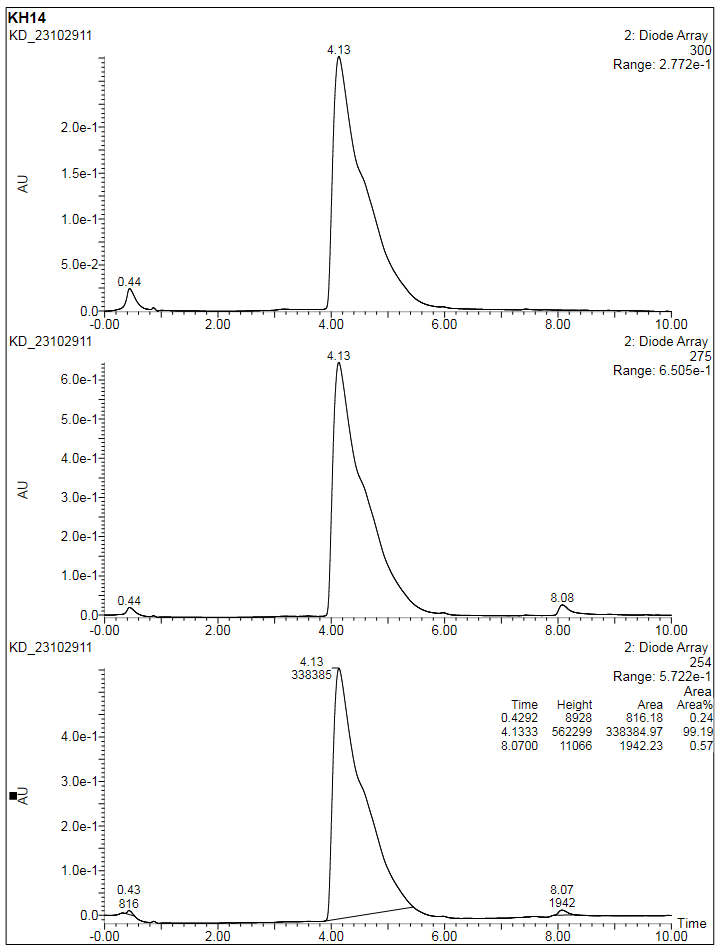

(2*S*,4*R*)-1-((*S*)-2-(3-(2-(2-((6-(4-(4-((4,4-dimethylpiperidin-1-yl)methyl)-2,5-difluorophenyl)-2-oxo-1,4,9-triazaspiro[5.5]undecan-9-yl)pyrimidin-4-yl)amino)ethoxy)ethoxy)propanamido)-3,3-dimethylbutanoyl)-4-hydroxy-N-(4-(4-methylthiazol-5-yl)benzyl)pyrrolidine-2-carboxamide (**KH15**).

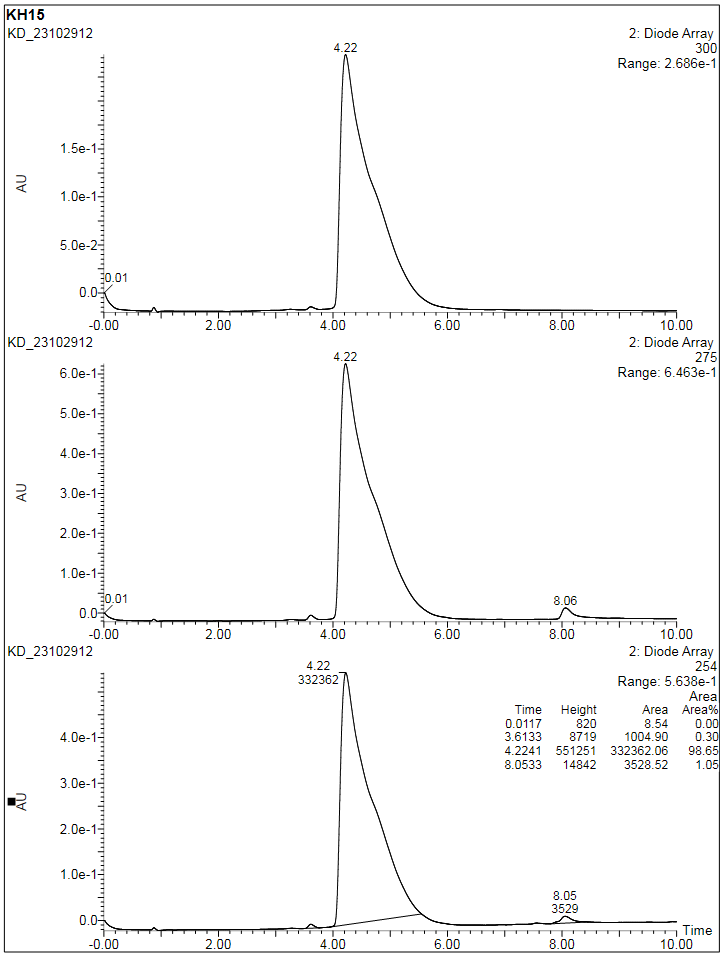

(2*S*,4*R*)-1-((*S*)-14-(tert-butyl)-1-((6-(4-(4-((4,4-dimethylpiperidin-1-yl)methyl)-2,5-difluorophenyl)-2-oxo-1,4,9-triazaspiro[5.5]undecan-9-yl)pyrimidin-4-yl)amino)-12-oxo-3,6,9-trioxa-13-azapentadecan-15-oyl)-4-hydroxy-N-(4-(4-methylthiazol-5-yl)benzyl)pyrrolidine-2-carboxamide (**KH16**).

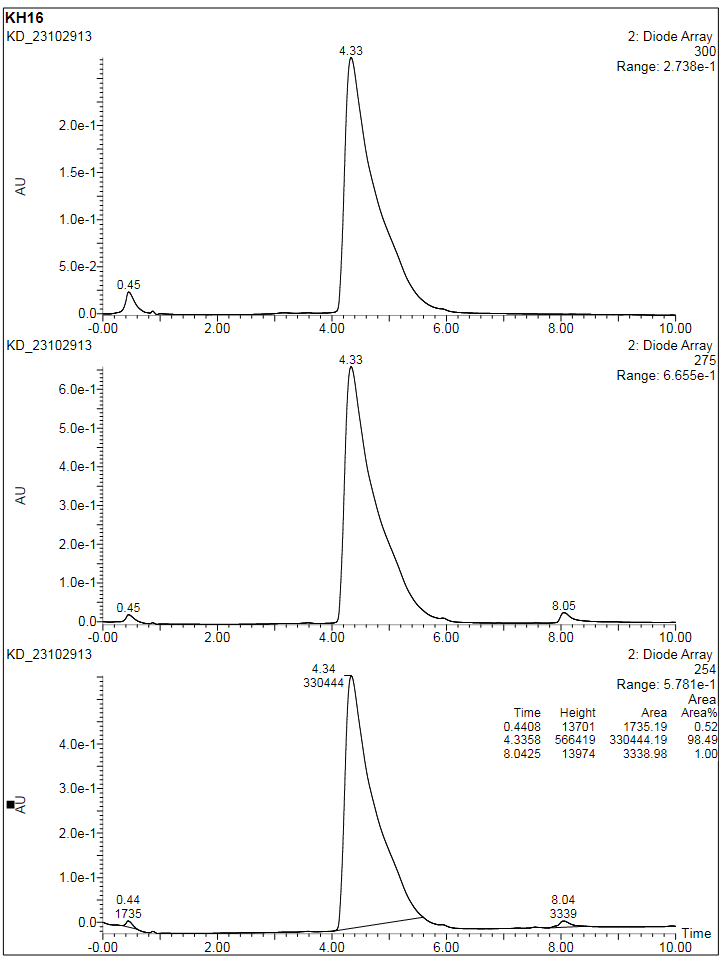

5-(4-(6-((6-(4-(4-((4,4-dimethylpiperidin-1-yl)methyl)-2,5-difluorophenyl)-2-oxo-1,4,9-triazaspiro[5.5]undecan-9-yl)pyrimidin-4-yl)amino)hexanoyl)piperazin-1-yl)-2-(2,6-dioxopiperidin-3-yl)isoindoline-1,3-dione (**KH21**).

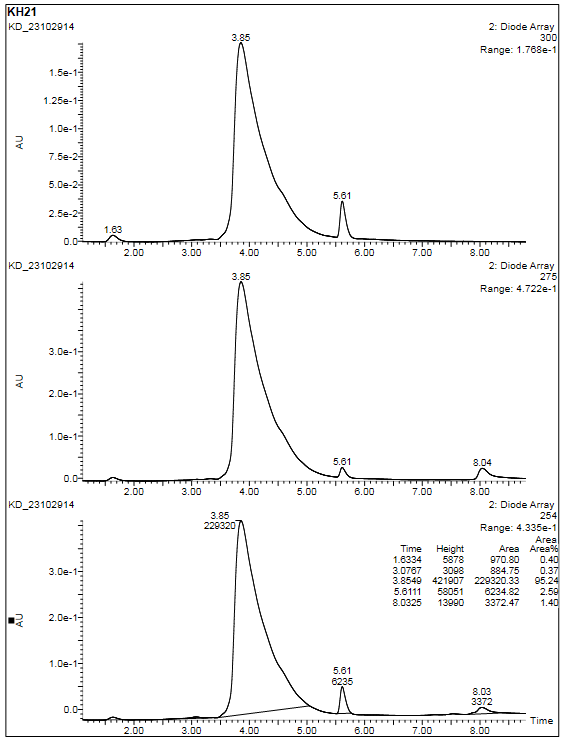

5-(4-(8-((6-(4-(4-((4,4-dimethylpiperidin-1-yl)methyl)-2,5-difluorophenyl)-2-oxo-1,4,9-triazaspiro[5.5]undecan-9-yl)pyrimidin-4-yl)amino)octanoyl)piperazin-1-yl)-2-(2,6-dioxopiperidin-3-yl)isoindoline-1,3-dione (**KH22**).

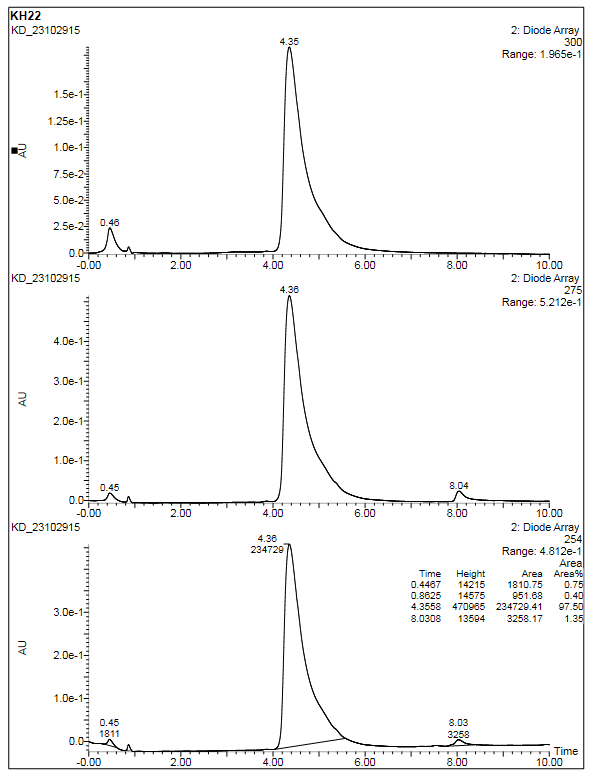

5-(4-(11-((6-(4-(4-((4,4-dimethylpiperidin-1-yl)methyl)-2,5-difluorophenyl)-2-oxo-1,4,9-triazaspiro[5.5]undecan-9-yl)pyrimidin-4-yl)amino)undecanoyl)piperazin-1-yl)-2-(2,6-dioxopiperidin-3-yl)isoindoline-1,3-dione (**KH23**).

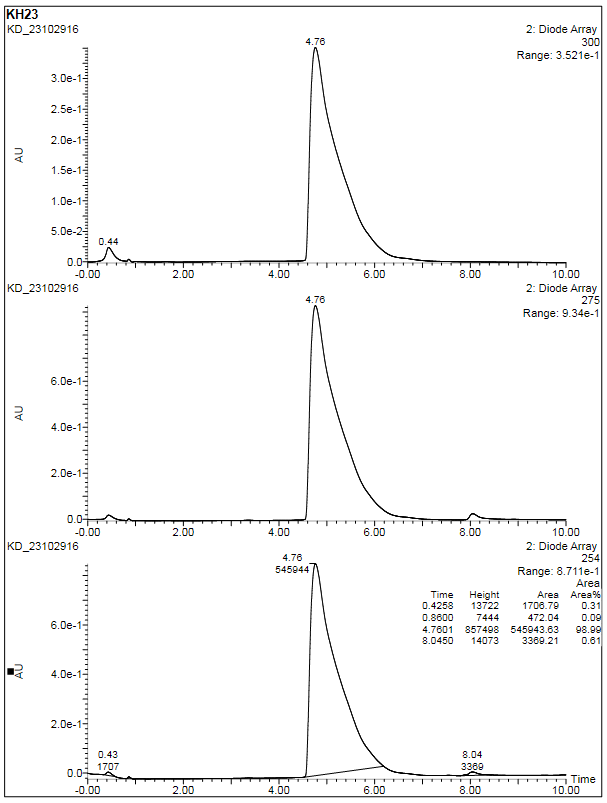

5-(4-(3-(2-((6-(4-(4-((4,4-dimethylpiperidin-1-yl)methyl)-2,5-difluorophenyl)-2-oxo-1,4,9-triazaspiro[5.5]undecan-9-yl)pyrimidin-4-yl)amino)ethoxy)propanoyl)piperazin-1-yl)-2-(2,6-dioxopiperidin-3-yl)isoindoline-1,3-dione (**KH24**).

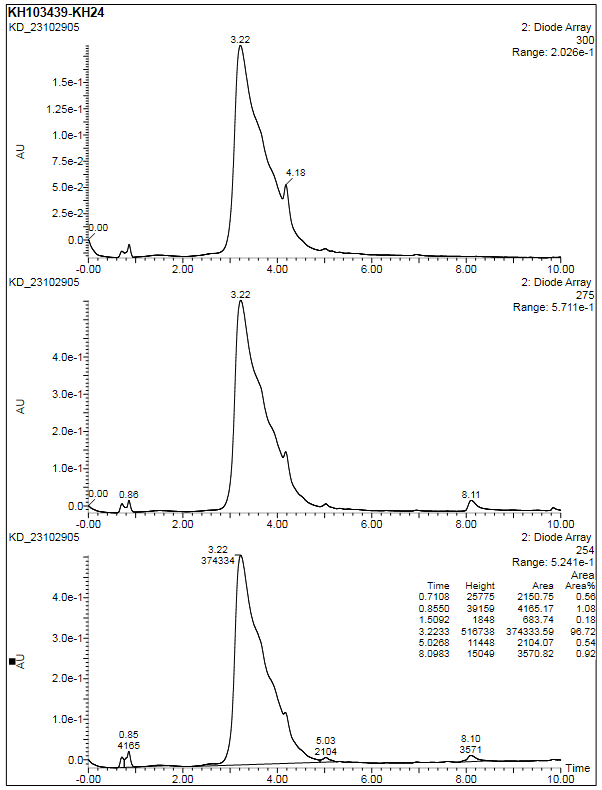

5-(4-(3-(2-(2-((6-(4-(4-((4,4-dimethylpiperidin-1-yl)methyl)-2,5-difluorophenyl)-2-oxo-1,4,9-triazaspiro[5.5]undecan-9-yl)pyrimidin-4-yl)amino)ethoxy)ethoxy)propanoyl)piperazin-1-yl)-2-(2,6-dioxopiperidin-3-yl)isoindoline-1,3-dione (**KH25**).

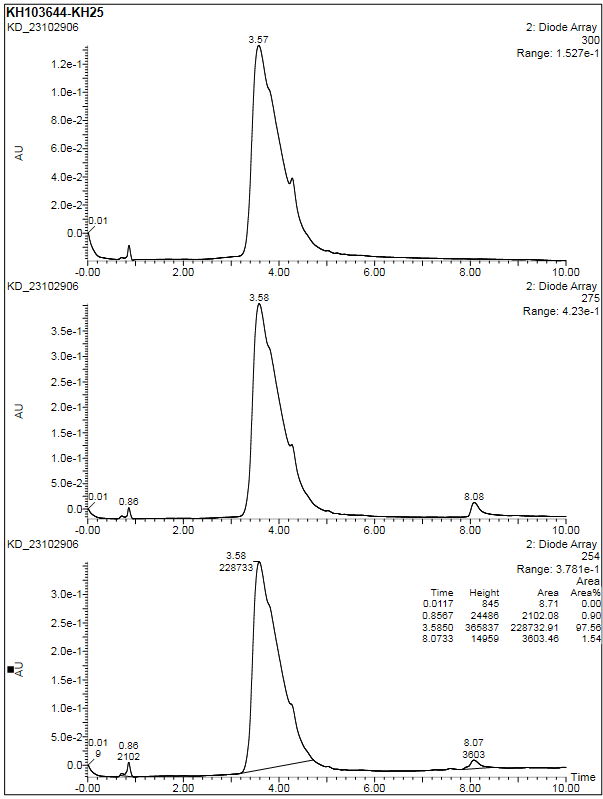

5-(4-(3-(2-(2-(2-((6-(4-(4-((4,4-dimethylpiperidin-1-yl)methyl)-2,5-difluorophenyl)-2-oxo-1,4,9-triazaspiro[5.5]undecan-9-yl)pyrimidin-4-yl)amino)ethoxy)ethoxy)ethoxy)propanoyl)piperazin-1-yl)-2-(2,6-dioxopiperidin-3-yl)isoindoline-1,3-dione (**KH26**)

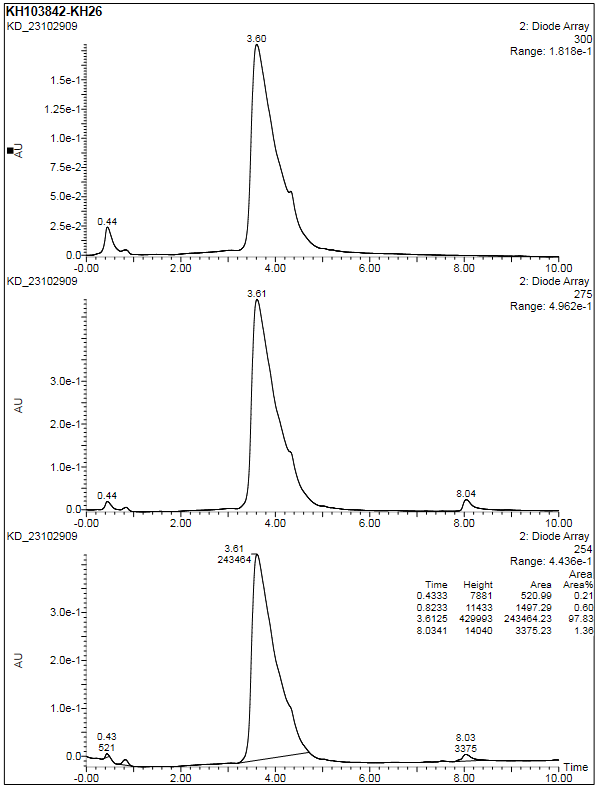

**ESI-HRMS spectra of METTL3 PROTACs**

(2*S*,4*R*)-1-((*S*)-2-(6-((6-(4-(4-((4,4-Dimethylpiperidin-1-yl)methyl)-2,5-difluorophenyl)-2-oxo-1,4,9-triazaspiro[5.5]undecan-9-yl)pyrimidin-4-yl)amino)hexanamido)-3,3-dimethylbutanoyl)-4-hydroxy-N-(4-(4-methylthiazol-5-yl)benzyl)pyrrolidine-2-carboxamide (**KH11**).

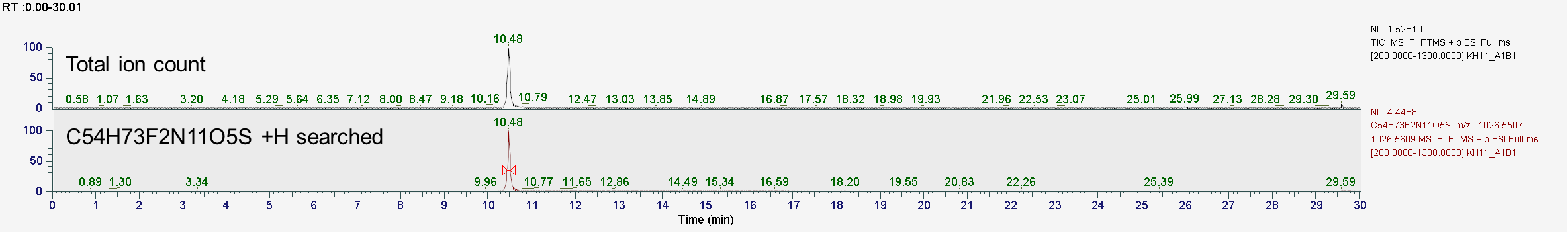

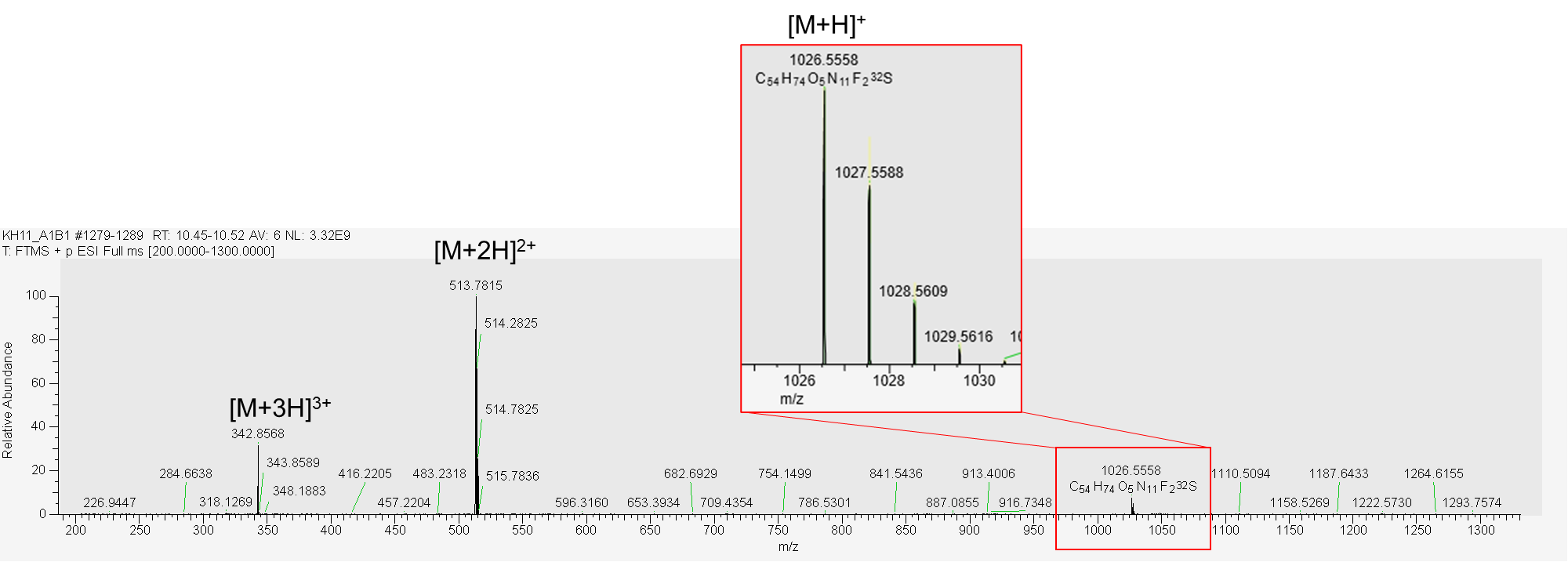

(2*S*,4*R*)-1-((*S*)-2-(8-((6-(4-(4-((4,4-Dimethylpiperidin-1-yl)methyl)-2,5-difluorophenyl)-2-oxo-1,4,9-triazaspiro[5.5]undecan-9-yl)pyrimidin-4-yl)amino)octanamido)-3,3-dimethylbutanoyl)-4-hydroxy-N-(4-(4-methylthiazol-5-yl)benzyl)pyrrolidine-2-carboxamide (**KH12**).

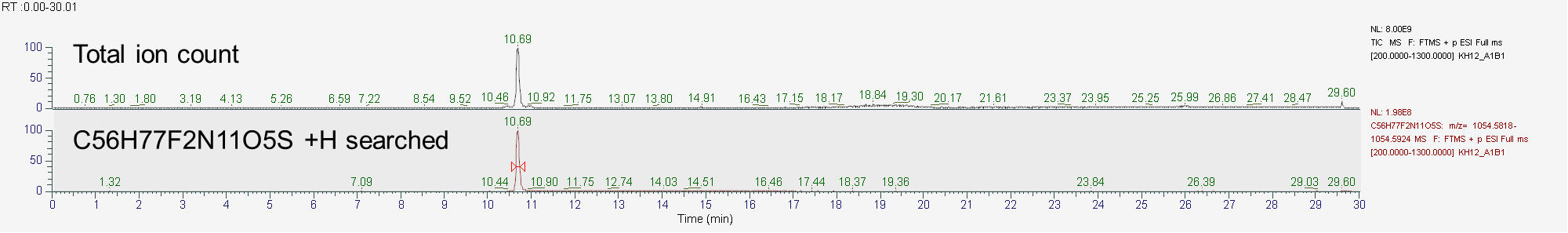

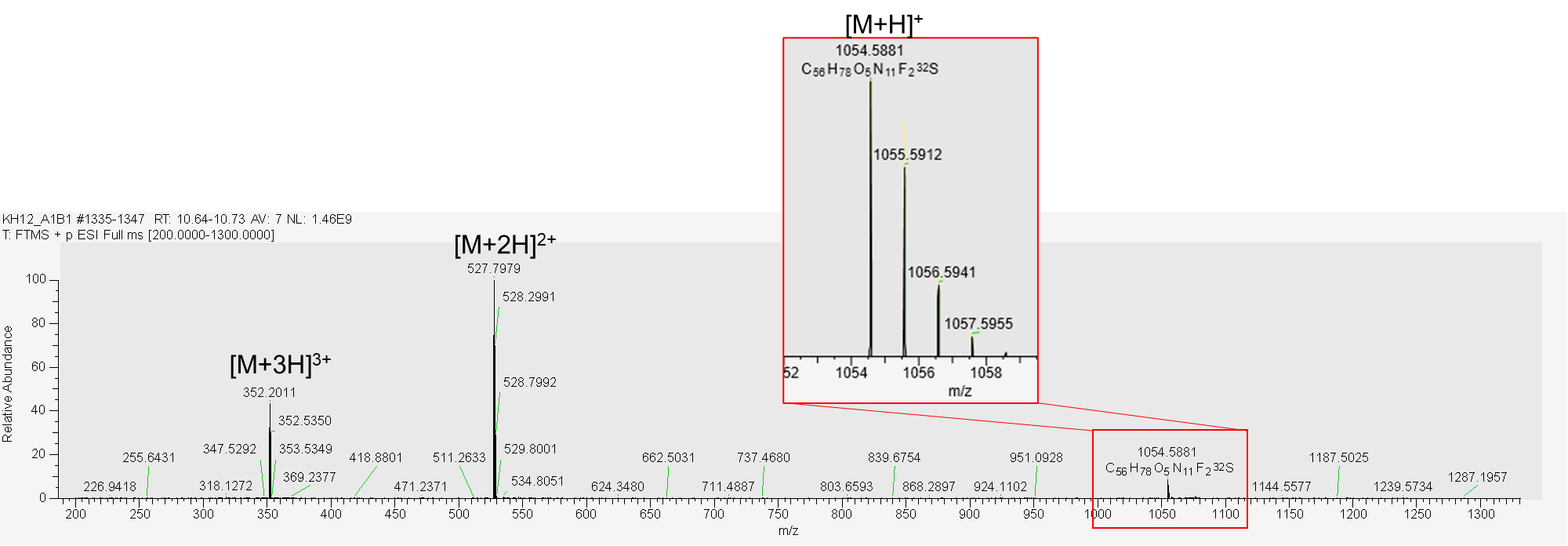

(2*S*,4*S*)-1-((*S*)-2-(8-((6-(4-(4-((4,4-dimethylpiperidin-1-yl)methyl)-2,5-difluorophenyl)-2-oxo-1,4,9-triazaspiro[5.5]undecan-9-yl)pyrimidin-4-yl)amino)octanamido)-3,3-dimethylbutanoyl)-4-hydroxy-N-(4-(4-methylthiazol-5-yl)benzyl)pyrrolidine-2-carboxamide (**KH12-NC**).

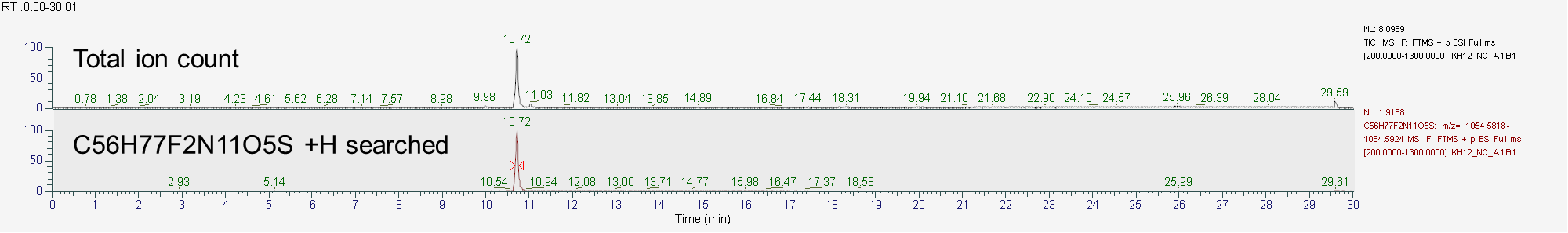

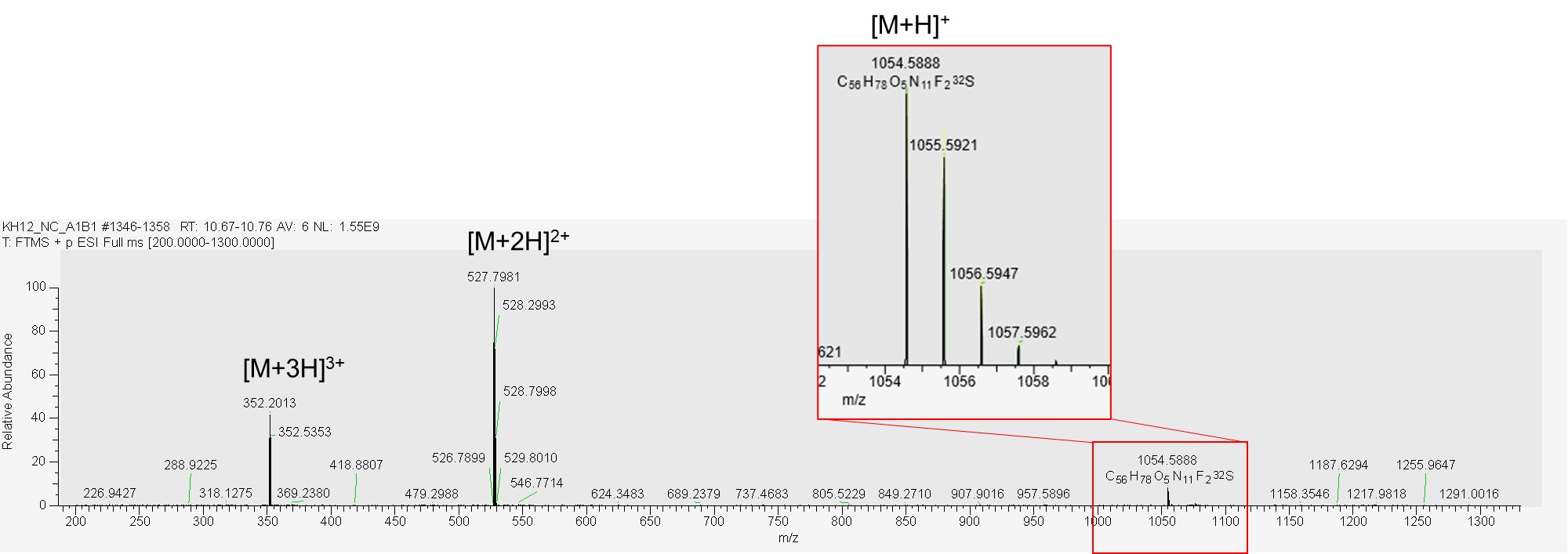

(2*S*,4*R*)-1-((*S*)-2-(11-((6-(4-(4-((4,4-dimethylpiperidin-1-yl)methyl)-2,5-difluorophenyl)-2-oxo-1,4,9-triazaspiro[5.5]undecan-9-yl)pyrimidin-4-yl)amino)undecanamido)-3,3-dimethylbutanoyl)-4-hydroxy-N-(4-(4-methylthiazol-5-yl)benzyl)pyrrolidine-2-carboxamide (**KH13**).

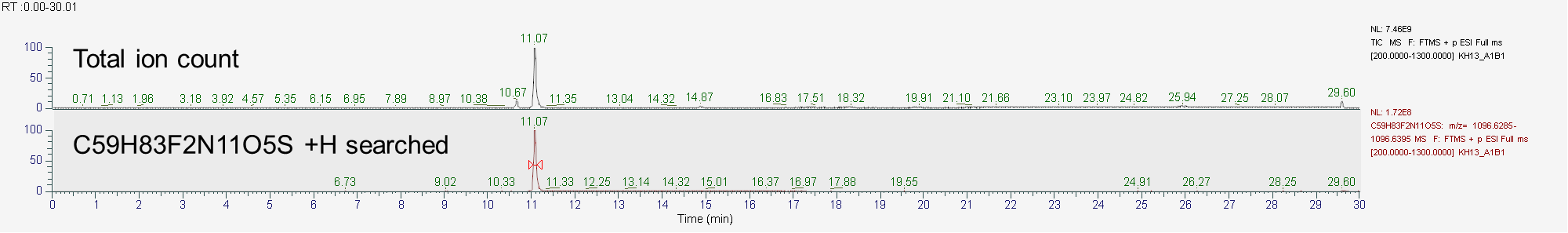

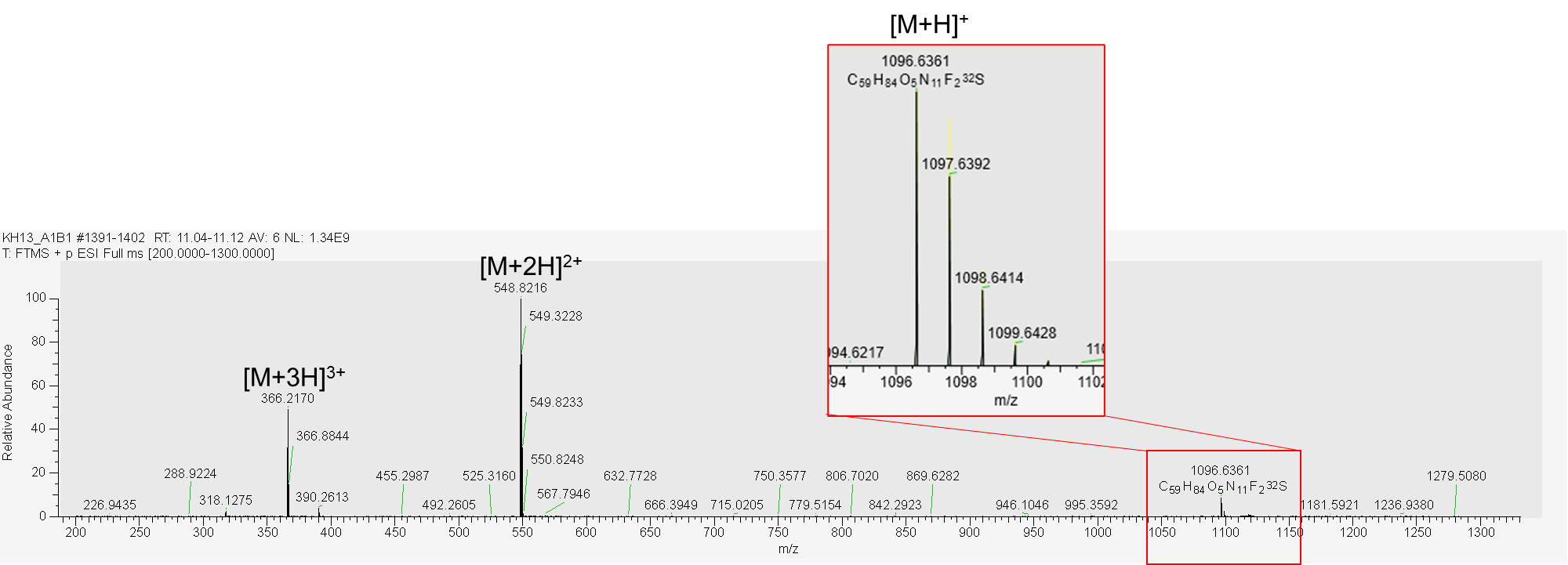

(2*S*,4*R*)-1-((*S*)-2-(3-(2-((6-(4-(4-((4,4-dimethylpiperidin-1-yl)methyl)-2,5-difluorophenyl)-2-oxo-1,4,9-triazaspiro[5.5]undecan-9-yl)pyrimidin-4-yl)amino)ethoxy)propanamido)-3,3-dimethylbutanoyl)-4-hydroxy-N-(4-(4-methylthiazol-5-yl)benzyl)pyrrolidine-2-carboxamide (**KH14**).

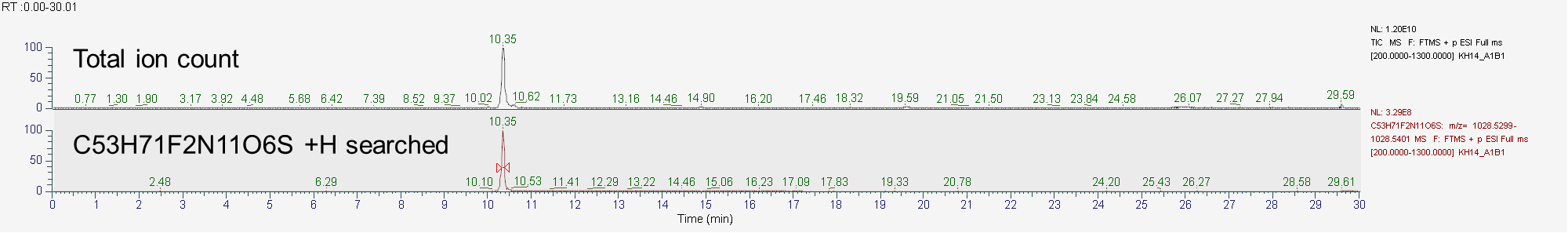

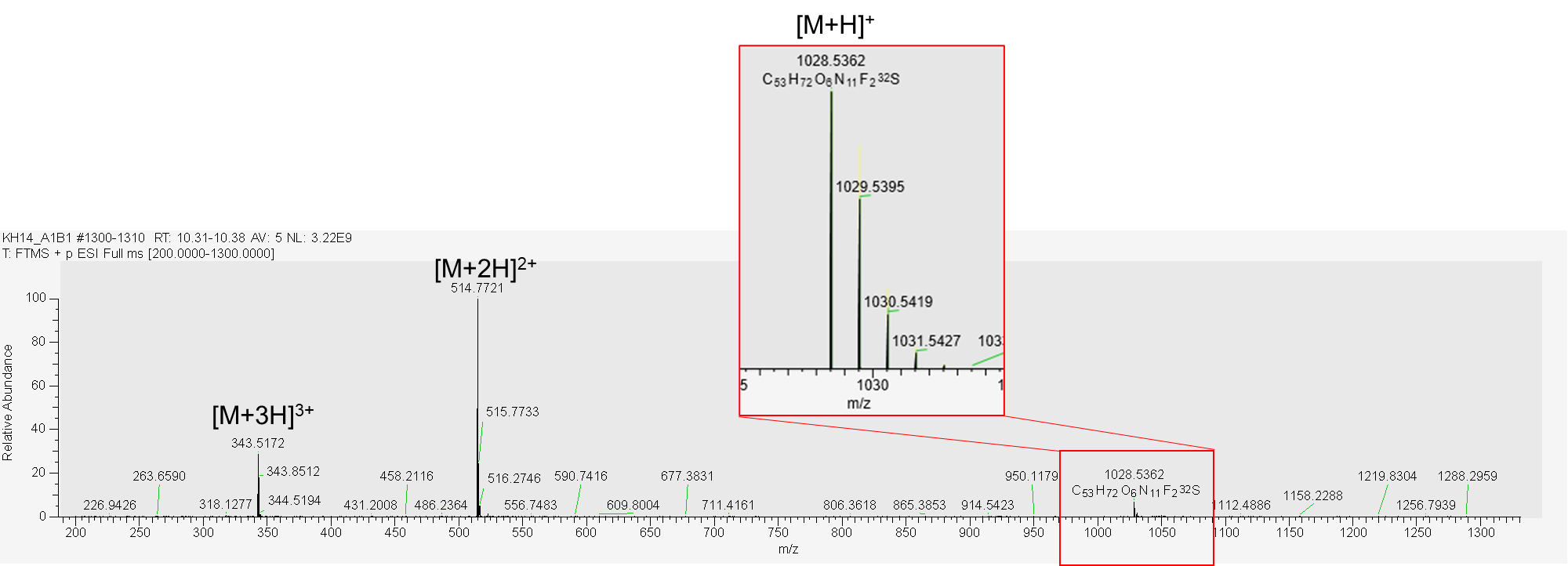

(2*S*,4*R*)-1-((*S*)-2-(3-(2-(2-((6-(4-(4-((4,4-dimethylpiperidin-1-yl)methyl)-2,5-difluorophenyl)-2-oxo-1,4,9-triazaspiro[5.5]undecan-9-yl)pyrimidin-4-yl)amino)ethoxy)ethoxy)propanamido)-3,3-dimethylbutanoyl)-4-hydroxy-N-(4-(4-methylthiazol-5-yl)benzyl)pyrrolidine-2-carboxamide (**KH15**).

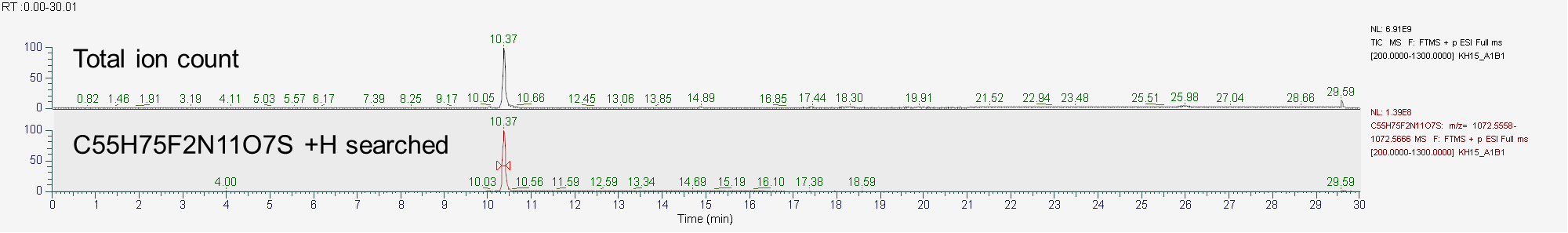

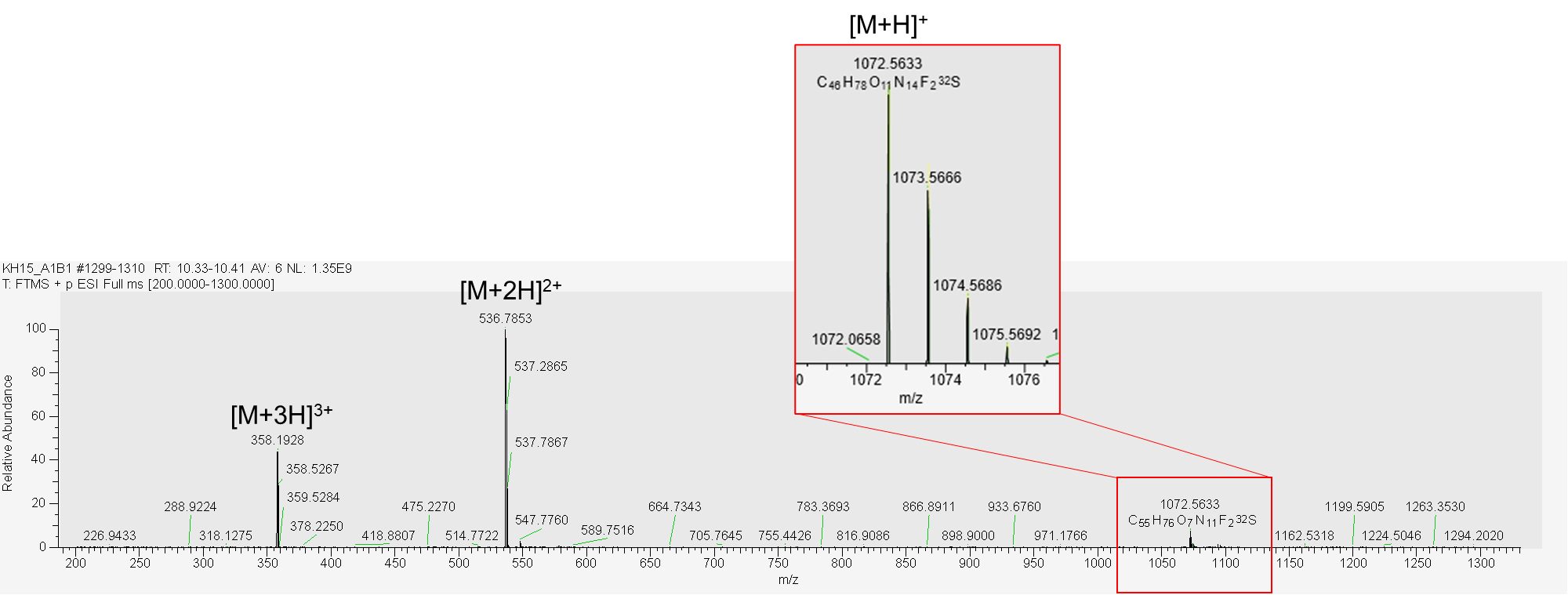

(2*S*,4*R*)-1-((*S*)-14-(tert-butyl)-1-((6-(4-(4-((4,4-dimethylpiperidin-1-yl)methyl)-2,5-difluorophenyl)-2-oxo-1,4,9-triazaspiro[5.5]undecan-9-yl)pyrimidin-4-yl)amino)-12-oxo-3,6,9-trioxa-13-azapentadecan-15-oyl)-4-hydroxy-N-(4-(4-methylthiazol-5-yl)benzyl)pyrrolidine-2-carboxamide (**KH16**).

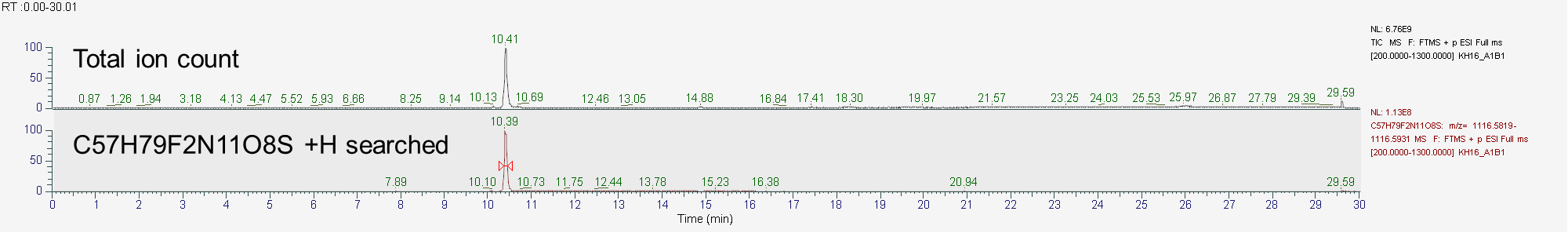

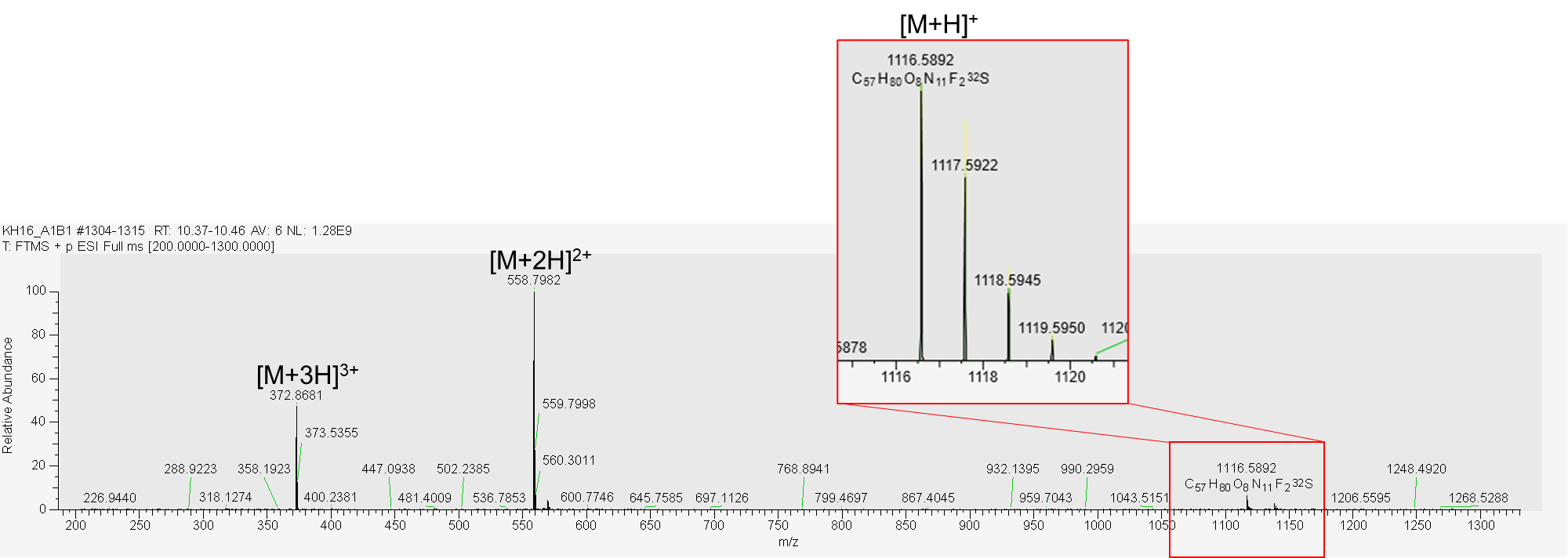

**References**

1	Dolbois, A. *et al.* 1,4,9-Triazaspiro[5.5]undecan-2-one Derivatives as Potent and Selective METTL3 Inhibitors. *J Med Chem* **64**, 12738-12760 (2021).
